# Age-related reorganization of locus coeruleus–cortical functional connectivity gradients

**DOI:** 10.64898/2026.02.05.704005

**Authors:** Arjun Dave, Shuer Ye, Xiaqing Lan, Alireza Salami, Heidi I.L. Jacobs, Maryam Ziaei

## Abstract

The locus coeruleus (LC) regulates attention, arousal, and adaptive behavior via widespread noradrenergic projections to cortex, yet its large-scale cortical organization and age-related vulnerability remain poorly understood. Using 7T MRI during naturalistic movie-viewing with neutral and negative emotional valence, study-specific LC delineation, and PET-derived receptor–transporter maps, we characterized LC–linked cortical organization in younger and older adults. Functional gradient analysis revealed two dominant axes: a stable primary gradient anchored by catecholaminergic receptor–transporter distributions, and a context-sensitive secondary gradient aligning with the visual–somatomotor (V–S) axis in neutral movie-viewing and the sensorimotor–association (S–A) axis in negative movie-viewing. During negative movie-viewing, older adults exhibited higher global and within-frontoparietal control network dispersion, indicating reduced functional differentiation, a pattern further associated with poorer emotional wellbeing. By revealing chemoarchitectural and context-sensitive LC–linked cortical organization, current work identifies mechanisms underlying age-related LC–linked cortical reorganization with consequences for mental health.

## Introduction

The locus coeruleus (LC) is the brain’s principal source of noradrenaline (NA). Through widespread projections to cortical and subcortical regions, the LC–NA system regulates attention, arousal, stress responses, and adaptive behavior by modulating large-scale brain dynamics^1–5^. LC neurons primarily operate in two firing modes: brief phasic bursts that enhance task-relevant processing and sustained tonic activity that maintains baseline arousal and behavioral flexibility^5,6^. By modulating neural gain, the LC–NA facilitates transitions between focused and exploratory behavioral states^5–9^. Convergent theoretical accounts, including the adaptive-gain^5^, network-reset^4,10^, and integration–segregation frameworks^11,12^, propose that the LC–NA system shapes large-scale functional organization to balance global integration and segregation in the cortex. Animal studies further demonstrate that elevated tonic LC activity can reshape long-range communication, supporting the LC’s role as a neuromodulatory hub for large-scale cortical dynamics^13^.

Given the LC–NA system’s capacity to influence cortex-wide patterns of activity^4,8,13–15^, a fundamental question arises: how is LC influence organized across the cortex? Specifically, do cortical regions couple to the LC in systematic patterns that reflect underlying biologically meaningful functional architecture? This question matters because the organization of LC–cortex connectivity has direct implications for cognitive and emotional function^16–20^. Prior LC connectivity work has largely treated cortical areas as discrete units or relied on static seed-to-voxel connectivity^20–22^, approaches that may not capture the broad organization of the cortex through the lens of the LC. Gradient-based mapping overcomes this limitation, positioning cortical regions according to their patterns of LC connectivity to reveal continuous transitions in LC–cortical functional organization^23–25^. Crucially, gradients are not merely functional abstractions. They align closely with underlying molecular architecture, including spatial distributions of neurotransmitter receptors and transporters, linking large-scale connectivity patterns to cortical chemoarchitecture^24,26,27^. Applying this framework to LC–cortical connectivity can reveal a systems-level map of how LC coupling is organized across the cortex, and how this organization relates to the underlying molecular architecture.

Aging provides a critical window during which cortical organization changes substantially. For instance, late adulthood is associated with reduced cortical differentiation, whereby functional networks become less distinct and less selectively organized^25,28–31^. This reduced differentiation is linked to poorer cognitive performance^25,28,30^ and greater vulnerability to mental health difficulties in later life^32^. According to a theoretical account developed by Li and colleagues across a sustained line of work^33–35^, these age-related changes reflect reduced neuromodulatory precision, which weakens neural gain control and reduces the selectivity of cortical responses. The LC is thus a strong mechanistic candidate, given its role as the primary source of cortical NA and its extensive projections to the cortex^1–7,10^. However, it remains unknown whether aging is associated with a reorganization of LC-cortical connectivity, and whether this reflects a general loss of cortical differentiation or network-specific alterations. Answering this question will provide important insight into the role of neuromodulatory systems in brain aging and inform prevailing theories of age-related neural reorganization.

Here, we address the gaps in three critical ways. First, we map the gradients of LC– cortical connectivity to characterize how the LC’s influence is embedded within large-scale cortical organization. Second, we assess age-related differences by quantifying dispersion within the LC–cortical functional connectivity gradient space and testing whether such changes are global or network-specific^36^. Finally, we test whether variation in LC–linked cortical dispersion relates to self-reported emotional wellbeing, assessing the behavioral relevance of LC-cortical connectivity topology in later life.

To address these objectives, we leveraged ultra-high field 7 Tesla Magnetic Resonance Imaging (7 T MRI), study-specific LC delineation, and a naturalistic movie-viewing paradigm to provide an ecologically valid framework to examine LC–linked cortical organization. Participants watched two movie clips differing in emotional valence (neutral and negative). We identified two dominant gradients of LC–cortical connectivity that explained the highest variance. The primary gradient remained stable across conditions and was tracked by cortical chemoarchitecture, specifically the density of catecholaminergic receptor–transporter distributions derived from publicly available PET-derived maps^37^. In contrast, the secondary gradient exhibited context-dependent organization, aligning with the canonical visual–somatomotor (V–S) axis in neutral movie-viewing and with the canonical sensorimotor–association (S–A) axis^23^ in negative movie-viewing. Analyses of the LC–linked cortical organization revealed that older adults exhibited higher LC–linked cortical dispersion specifically during negative movie-viewing, with the frontoparietal control network (FPN) emerging as the primary site of age-related vulnerability. Critically, higher global and FPN dispersion in older adults associated with poorer emotional wellbeing. Collectively, our findings provide a systems-level map of LC–linked cortical organization and reveal that age-related reduced functional differentiation within this organization carries relevance for mental health in later life.

## Results

### Two dominant gradients of LC–cortical connectivity organization

We characterized LC–cortical connectivity organization in 140 participants: 72 younger adults (mean age: 25.75 ± 4.25 years; range: 19–36 years; 34 females) and 68 older adults (mean age: 70.90 ± 3.83 years; range: 65–82 years; 36 females) during the two naturalistic movie-viewing conditions (**Figure 1a)** followed by a separate behavioral session assessing emotional wellbeing and cognitive performance **(Figure 1b)**. Post movie-viewing, self-reported valence and arousal ratings confirmed that negative movie-viewing elicited lower positive valence and higher arousal compared with baseline and neutral viewing, with comparable effects across age groups (**Supplementary Figure 1**).

**Figure 1:**
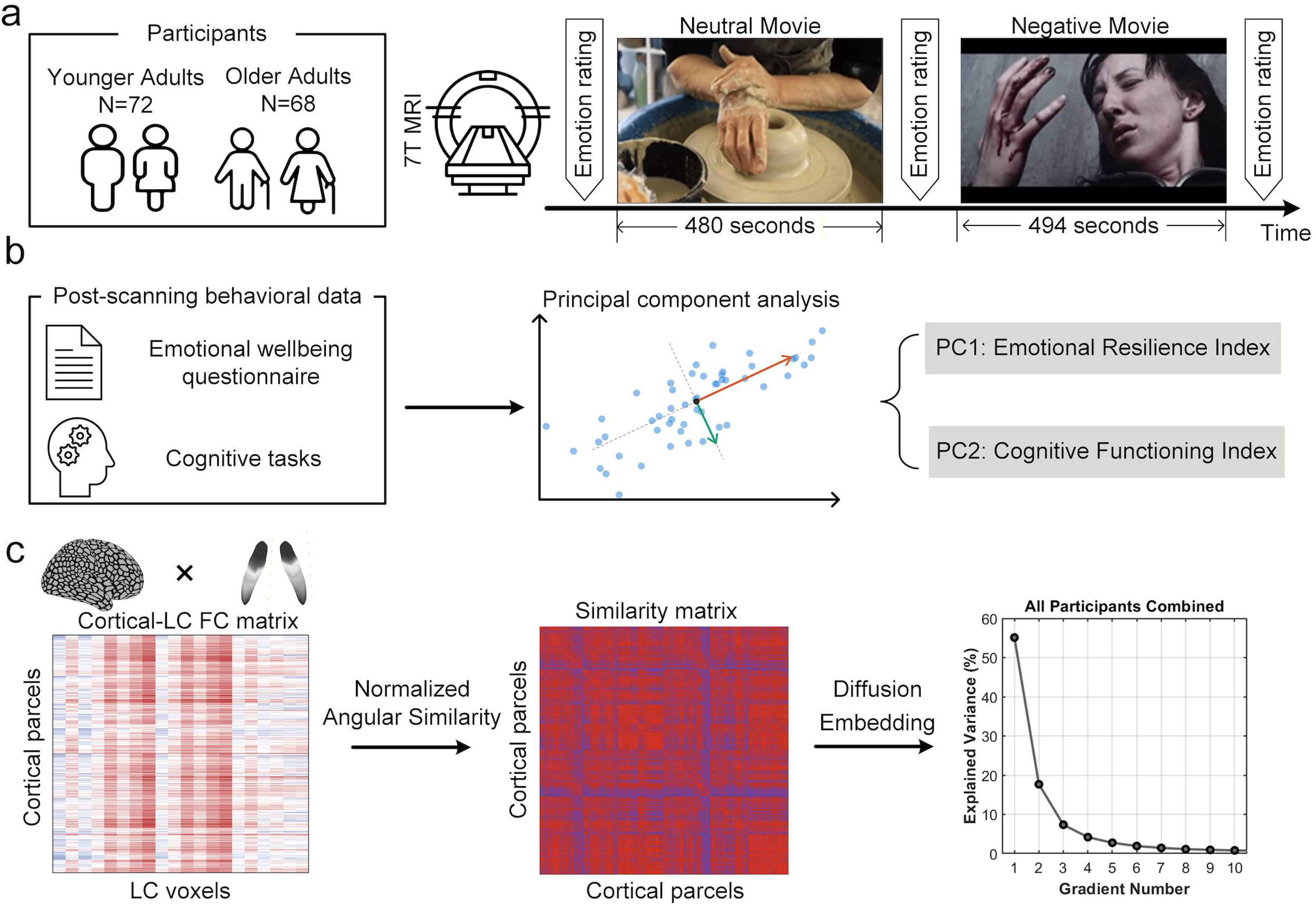
Overview of the study design and gradient analyses. **(a)** Ultra-high-field 7 T fMRI data were acquired from 72 younger and 68 older adults during naturalistic viewing of two movie clips differing in emotional valence and arousal: a neutral clip (480 s) followed by a negative clip (494 s). Participants rated their valence and arousal before scanning and immediately after each clip. **(b)** In a separate session outside the scanner, participants completed a battery of emotional wellbeing questionnaires and cognitive tasks. Principal component analysis (PCA) on the standardized measures yielded two composite indices: an Emotional Resilience Index (ERI) summarizing emotional wellbeing, and a Cognitive Functioning Index summarizing cognitive performance. **(c)** For each participant, movie condition and LC side, functional connectivity was computed between LC voxels and 1,000 cortical parcels of the Schaefer atlas^39^, yielding a cortical-parcel × LC-voxel connectivity *matrix*. A parcel-by-parcel similarity matrix was then constructed from these LC connectivity profiles using normalized angular (cosine) similarity^40^, and diffusion map embedding^41^was applied to derive low-dimensional gradients capturing the principal axes of variance in LC– cortical coupling. The scree plot shows the percentage of variance explained by the first ten gradients across all participants; gradients were ranked by variance explained, and individual gradients were aligned to the group-level template using Procrustes rotation prior to statistical analysis.

To derive LC–cortical functional connectivity gradients, we first computed subject-specific LC–cortical connectivity matrices (**Figure 1c**) separately for the neutral and negative movie-viewing conditions, and for the left and right LC. LC voxel-wise time series extracted from study-specific LC masks delineated on the magnetization transfer-weighted turbo flash (MT-TFL) sequence^38^ (**Supplementary Figure 2**), were correlated with the time series of Schaefer 1,000 parcels atlas^39^ with Yeo 7-network assignment^36^. Correlations were Fisher z-transformed to yield a per-participant LC-voxel × cortical-parcel connectivity matrix. For each condition, individual connectomes were averaged across participants to form a group-mean LC–cortical matrix. Using the BrainSpace toolbox^40^, a cortex × cortex affinity matrix was computed using the normalized angular (cosine) similarity between cortical parcels based on their LC connectivity; this quantifies how similarly cortical regions are coupled to the LC. Diffusion map embedding^41^ was then applied to this affinity matrix to derive low-dimensional gradient components^23,40^. The resulting components served as the condition-specific group reference template. Individual-participant gradients were further computed and aligned to this template using Procrustes rotation to ensure cross-subject comparability. Averaging the aligned dividual gradients yielded the group-level gradient map across all participants, as well as separate age-specific maps for younger and older adults.

Gradient 1 (G1) accounted for the largest proportion of variance in LC–cortical coupling across both movie-viewing conditions (**Figure 2; neutral movie-viewing**: Right LC = 48%, Left LC = 49%; **negative movie-viewing**: Right LC = 56%, Left LC = 61%). G1 exhibited highly consistent spatial topology across emotional conditions (**Supplementary Figure 3;** Right LC: *ρ* = 0.74, *p_spin_* < 0.001; Left LC: *ρ* = 0.83; *p_spin_* < 0.001) and between the right and left LC, with strong spatial correspondence in both movie-viewing conditions (**Supplementary Figure 4;** negative movie: *ρ* = 0.92; neutral movie: *ρ* = 0.95; both *p_spin_* < 0.001). Gradient 2 (G2) was less stable, with lower cross-condition correspondence than G1 (Right LC: *ρ* = 0.26; Left LC: *ρ* = 0.52). Because gradient orientation is arbitrary^40,42^, we examined LC–cortical coupling at the extremes of G1 to characterize its functional significance in both neutral and negative movie-viewing conditions. Parcels at the positive extreme (top 5% and top 10% of G1 values) showed strong positive coupling with the LC, whereas parcels at the negative extreme exhibited weak or negative coupling **(Supplementary Figure 5)**.

**Figure 2:**
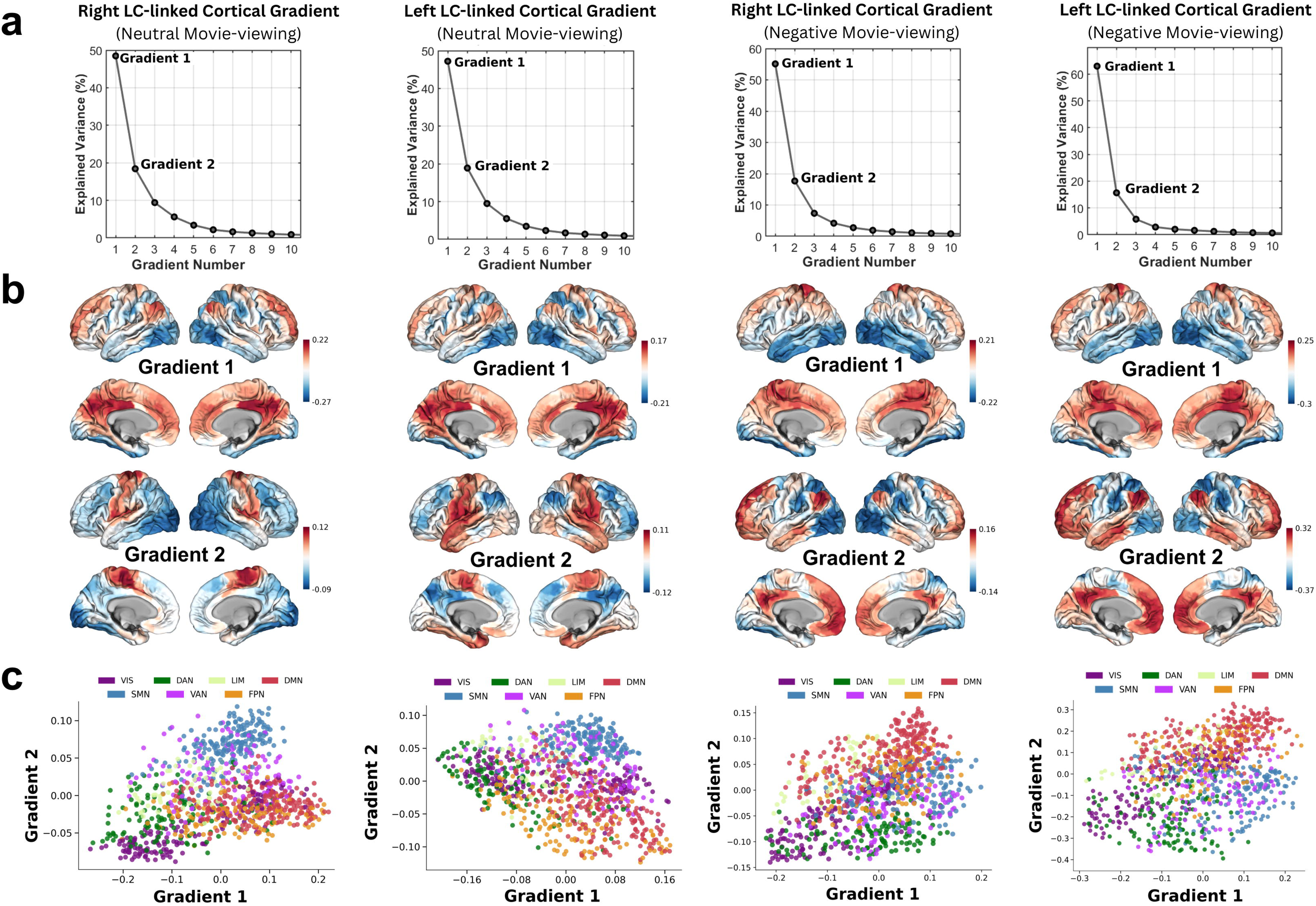
Group-level LC–linked cortical gradients shown separately for the right and left LC under neutral and negative movie-viewing conditions. (columns, left to right: right LC–cortex neutral, left LC–cortex neutral, right LC–cortex negative, left LC–cortex negative). (a) Scree plots showing the percentage of explained variance for the first ten gradients; the first two gradients (G1, G2) capture the largest proportion of explained variance in all four embeddings and are retained for subsequent analyses. (b) Cortical surface projections of Gradient 1 (top) and Gradient 2 (bottom), shown as lateral and medial views of both hemispheres; color reflects gradient loading, with colorbar ranges scaled per panel. (c) Joint G1 × G2 embedding of cortical parcels, colored by Yeo seven-network assignment36. Network abbreviations: visual (VIS), somatomotor (SMN), dorsal attention (DAN), ventral attention (VAN), limbic (LIM), frontoparietal (FPN), and default mode (DMN).

The spatial distribution of G1 revealed a graded pattern of LC–cortical coupling across large-scale networks. Regions within the default mode network (DMN), FPN, and somatomotor network (SMN) including precuneus, posterior cingulate cortex, medial prefrontal cortex, and motor regions occupied the positive end of G1, indicating stronger LC coupling. This pattern is consistent with anatomical and physiological evidence of broad noradrenergic projections from the LC to association and sensorimotor cortices, supporting the LC’s role in influencing distributed cortical systems^43,44^. In contrast, visual, limbic, and dorsal attention regions occupied the negative end, reflecting weaker LC coupling. Comparison with canonical cortical gradients^23^ showed nonsignificant alignment of G1 with canonical S–A and V–S axes across both movie-viewing conditions (**Supplementary Figure 6**). Neurosynth meta-analytic decoding (Hungarian spin-permutation test^45^, 10,000 spins; *p_spin_* < 0.05) revealed consistent functional associations across both emotional conditions. Parcels at the positive end of G1 were associated with cognitive control, salience, and affective-arousal processes, including decision-making, inhibition, monitoring, anticipation, hyperactivity, salience, risk, stress, and pain. Parcels at the negative end were linked to perceptual and language-related processes, including object and word recognition, facial expression processing, gaze, discrimination, reading, and language. This pattern indicates that the primary LC–linked cortical gradient captures a functional axis from higher-order control and salience processing to perception, recognition, and language (**Extended Data Figure 1**).

Gradient 2 (G2) explained a smaller proportion of variance (**Figure 2; neutral movie-viewing**: Right LC = 20%, Left LC = 19%; **negative movie-viewing**: Right LC = 19%, Left LC = 18%) and captured a distinct and condition-dependent organizational axis. During neutral movie-viewing, G2 primarily differentiated visual and somatosensory cortices, aligning with the canonical V–S axis (**Figure 3b**; Left LC *ρ* = 0.57, Right LC *ρ* = 0.48; both *p_spin_* < 0.001). In contrast, during negative movie-viewing, G2 was anchored by higher-order association regions, including posterior cingulate cortex, prefrontal cortex, and temporal pole on one extreme and by occipital and dorsal attention regions at the other, resembling the canonical S–A axis (**Figure 3b**; Left LC *ρ* = 0.75, Right LC *ρ* = 0.63; both *p_spin_* < 0.001). Additionally, G1 showed significantly stronger cross-condition similarity than G2 (Right LC: z = 4.10, *p* < 0.001; Left LC: z = 3.08, *p* = 0.002), confirming that G1 captures a stable organizational pattern while G2 exhibits context-dependent reorganization. Neurosynth meta-analytic decoding of G2 for both conditions is presented in **Extended Data Figure 1**.

**Figure 3:**
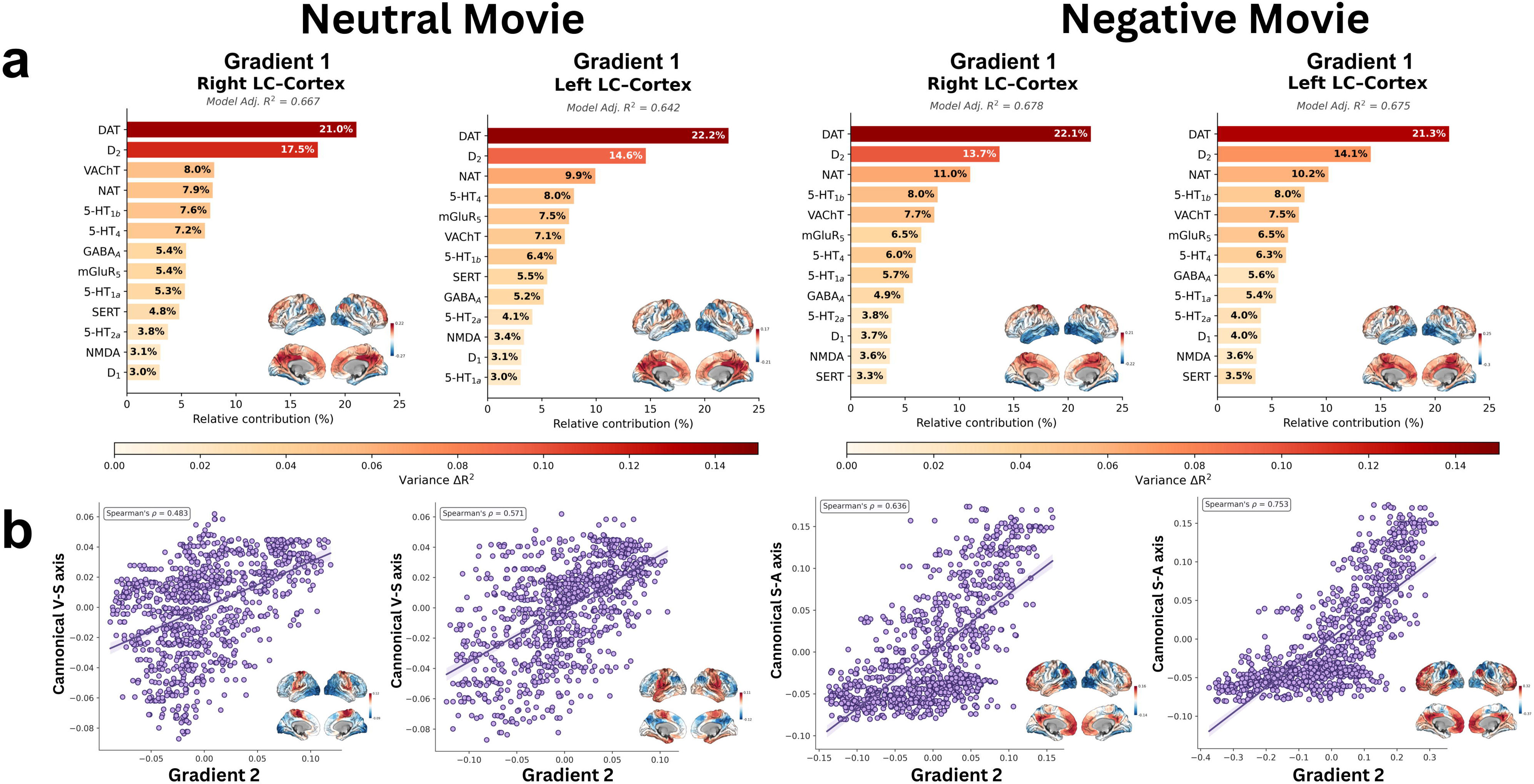
Chemoarchitectural and canonical-gradient alignment of LC–linked cortical Gradients 1 and 2 across emotional conditions and hemispheres. **(a)** Dominance analysis quantifying the relative contribution of 13 neurotransmitter receptor and transporter density maps to the spatial topography of Gradient 1, derived separately for the right and left LC under neutral (left two columns) and negative (right two columns) movie-viewing conditions. Bars show the relative contribution (%) of each receptor or transporter to the total explained variance, with bar color encoding absolute variance contribution (ΔR², shaded colorbar). Model-level adjusted R² is reported above each panel. **(b)** Scatter plots showing the spatial alignment between Gradient 2 and canonical cortical gradients^23^ across the same four hemisphere × condition combinations (Spearman’s ρ; significance assessed via spin permutation, see Methods). During neutral movie-viewing, G2 aligned with the canonical visual–somatomotor (V–S) axis, whereas during negative movie-viewing, G2 aligned with the canonical sensorimotor–association (S–A) axis. Inset cortical surfaces show the corresponding Gradient 1 and Gradient 2 group-level maps. Receptor/transporter abbreviations: DAT, dopamine transporter; D₁/D₂, dopamine receptors; NAT, noradrenaline transporter; 5-HT_1a_ / 5-HT_1b_ / 5-HT_2a_ / 5-HT_4_ , serotonin receptor subtypes; SERT, serotonin transporter; mGluR₅, metabotropic glutamate receptor 5; NMDA, N-methyl-D-aspartate receptor; GABAₐ, γ-aminobutyric acid type A receptor; VAChT, vesicular acetylcholine transporter.

### G1 aligns with catecholaminergic architecture

Functional connectivity gradients have been shown to align with underlying neuromodulatory architectures^24,26,27^. To characterize the chemoarchitectural basis of LC–linked cortical organization, we included a broader set of receptors–transporter distribution maps. Specifically, we used an extension of multiple regression to link parcel-wise G1 values to thirteen PET-derived receptor–transporter density maps (**Supplementary Table 1**) spanning major neurotransmitter systems from JuSpace toolbox^37^. Because neurotransmitter receptors and transporters show substantial spatial overlap^26,46^, we applied dominance analysis^47^ to quantify the unique contribution of each receptor–transporter profile to the model’s adjusted explained variance (R²_adj_), providing estimates of their independent effects.

The receptor–transporter density model explained a large proportion of G1 variance across both emotional conditions (**Figure 3a**; **neutral movie-viewing**: Right LC R²_adj_ = 0.667; Left LC R²_adj_ = 0.642; **negative movie-viewing**: Right LC R²_adj_ = 0.678; Left LC R²_adj_ = 0.675). Critically, three catecholaminergic markers: dopamine transporter (DAT), dopamine D2 receptor (D2), and NAT (noradrenaline transporter) emerged as dominant predictors, jointly accounting for 47% (right LC–linked cortical G1) and 46% (left LC–linked cortical G1) of total explained variance across conditions. This consistency across emotional conditions supports the interpretation of G1 as a stable axis of LC–cortical connectivity tracked by density of cortical catecholaminergic chemoarchitectural organization.

To further characterize these relationships, we examined associations between G1 and the dominant catecholaminergic markers. NAT exhibited a positive association with G1 scores, with strongest associations within the FPN, DMN, and SMN (**Supplementary Figure 7a**; Right LC: *ρ = 0.29, p < 0.001; Left LC: ρ = 0.22, p < 0.001*), whereas DAT showed a negative association (Right LC: *ρ* = −0.46, *p <* 0.001; Left LC: *ρ* = −0.45, *p <* 0.001; **Supplementary Figure 7b**). No significant correlation was observed between G1 scores and D2 receptor density (*p* > 0.05). In contrast, G2 was poorly explained by the receptor–transporter density profiles, with substantially lower model fits (Right LC–cortex: R²_adj_ = 0.23; Left LC–cortex: R²_adj_ = 0.11).

### LC functional organization reflects rostro-caudal differentiation

To validate whether our gradient approach detects meaningful spatial structure within the LC, we transposed, for each subject, the LC-voxel × cortical-parcel matrix and computed normalized angular (cosine) similarity between LC voxels based on their cortical connectivity fingerprints and applied diffusion map embedding. The primary within-LC gradient (G1) accounted for a substantial proportion of variance (Right LC: 75.7%; Left LC: 71.0%) and was spatially correlated with the rostro-caudal axis, approximated by the MNI Z coordinate (**Supplementary Figure 8;** Right LC: *r* = 0.68, *p* < 0.001; Left LC: *r* = 0.73, *p* < 0.001). Together, these results replicate recent reports of rostro-caudal functional differentiation within the LC^17^.

### Age-related differences in LC–linked cortical gradient organization

To characterize age-related differences in LC–linked cortical functional organization, we compared group-level primary (G1) and secondary (G2) cortical gradient maps derived from right and left LC seeds between younger and older adults, separately for the neutral and negative movie-viewing conditions. Age effects were quantified at two complementary levels: (i) positioning of canonical functional networks^36^ within the 2D gradient manifold^25,31^, and (ii) spatial similarity of gradient topography in younger and older adults.

During negative movie-viewing, age-related position shifts were largest in sensory and attentional networks — the visual network (VIS) for both right and left LC–linked cortical gradients, and in the DAN, particularly for the left LC–linked cortical gradient (**Figure 4; Supplementary Table 2**). During neutral movie-viewing, age-related shifts were largest in the SMN for both right and left LC–linked cortical gradients (**Supplementary Figure 9; Supplementary Table 3)**. In both conditions, these shifts occurred primarily along the G1, while higher-order association networks (FPN, VAN, DMN) showed smaller repositioning.

**Figure 4:**
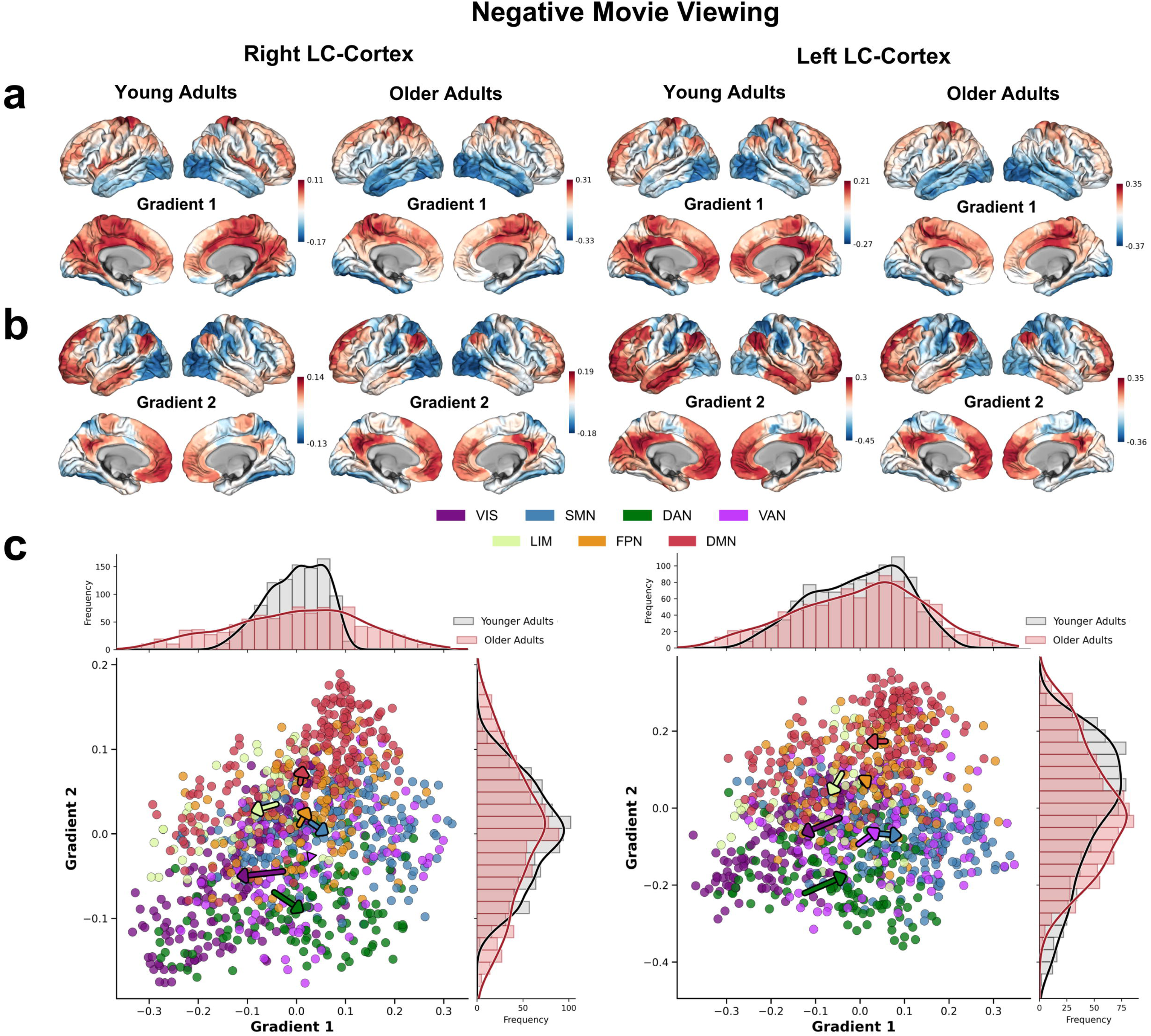
Age-related network shifts within the two-dimensional (G1 × G2) manifold of LC–linked cortical organization during negative movie-viewing. (a, **b)** Group-averaged cortical maps of **(a)** Gradient 1 and **(b)** Gradient 2 for younger and older adults, derived separately from the right LC–linked cortical gradient (left columns) and left LC–linked cortical gradient (right columns); color reflects gradient loadings. **(c)** Joint G1 × G2 embedding of cortical parcels for the right LC–linked cortical gradient (left panel) and left LC–linked cortical gradient (right panel) of older adults, with parcels colored by their Yeo seven-network assignment^36^ (VIS, SMN, DAN, VAN, LIM, FPN, DMN). Marginal histograms above and to the right of each scatter plot show the distributions of younger (grey) and older (red) adult gradient scores along G1 (top) and G2 (right). Arrows depict age-related centroid displacement: each arrow originates at the younger-adult network centroid and terminates at the older-adult centroid, with arrow length indicating Euclidean displacement and direction indicating the relative ΔG1 and ΔG2 contributions. Network abbreviations: VIS, visual; SMN, somatomotor; DAN, dorsal attention; VAN, ventral attention; LIM, limbic; FPN, frontoparietal; DMN, default mode.

Additionally, the overall topography of LC–linked cortical gradients remained largely preserved across age groups. Across both gradients, LC side, and movie-viewing conditions, younger-and older-adult group-level gradient maps exhibited high spatial correlation (**Supplementary Figure 10**).

Together, these analyses indicate that the macroscale topography of LC–linked cortical organization is largely preserved with age at the group level, consistent with the previous findings^25,31^. To further examine whether age-related differences emerge at the individual level, despite preserved group-level organization, we next quantified dispersion in 2D (G1 × G2) gradient space, following an established multivariate framework^25,48^.

### Age-related increase in LC–linked cortical dispersion during negative movie-viewing

For each participant, the first two gradients (G1 and G2) defined the embedding space for computing three complementary dispersion metrics: global dispersion was defined as the mean Euclidean distance of all cortical parcels from the whole-cortex centroid; within-network dispersion as the mean Euclidean distance of parcels within a given functional network from that network’s centroid; and between-network dispersion as the mean Euclidean distance between the centroids of each network pair^25,48^ (*schematic and empirical intuitive illustration in **Supplementary Figure 11** and **Supplementary Figure 12**, respectively).* To account for potential confounds related to age-related cortical atrophy^25^, all statistical models included mean cortical thickness, sex, and head motion as covariates.

Older adults exhibited significantly higher global dispersion than younger adults during negative movie-viewing for the right LC–linked global cortical dispersion (**Figure 5**; *t* = −2.94, *p* = 0.004, *d* = 0.50; 95% CI [−0.145, −0.028]) indicating a more heterogeneous cortical embedding in gradient space. The left LC–linked global cortical dispersion showed a directionally consistent but non-significant trend (*t* = −1.70, *p* = 0.092, *d* = 0.29, 95% CI [−0.152, 0.012]).

**Figure 5:**
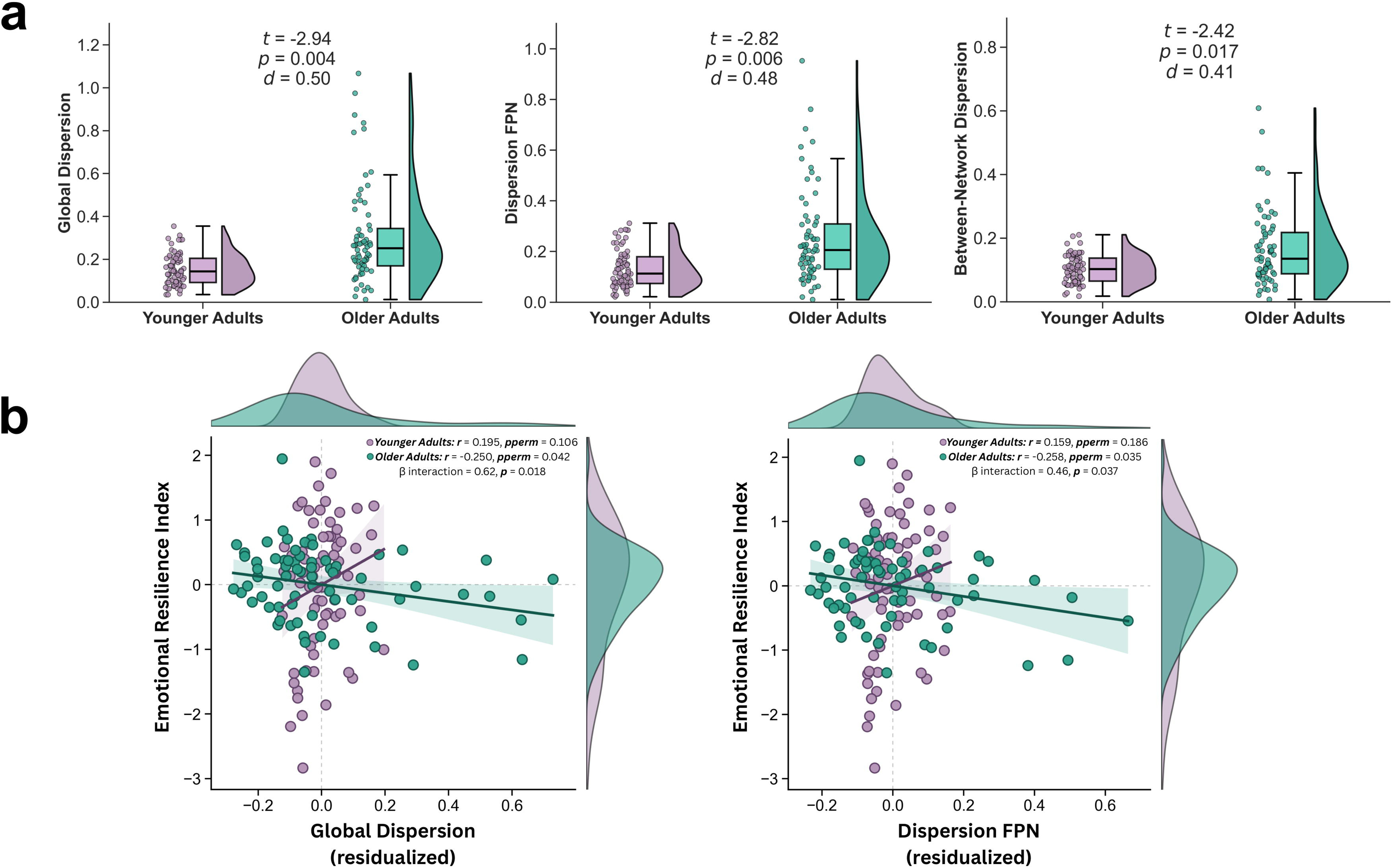
Age-related differences in LC–linked cortical dispersion during negative movie-viewing and its associations with emotional wellbeing. **(a)** Raincloud plots showing the distribution of subject-level dispersion values in younger adults (purple) and older adults (teal) across three complementary dispersion metrics derived from the right LC–linked cortical gradient manifold (G1 × G2). Left: global dispersion, computed as the mean Euclidean distance of all cortical parcels from the overall centroid. Middle: within-network dispersion for the frontoparietal network (FPN), computed as the mean Euclidean distance of FPN parcels from the FPN centroid. Right: between-network dispersion, computed as the mean Euclidean distance between all pairs of Yeo seven-network centroids. Each dot represents one participant; box plots display the median and interquartile range, and density curves show the distribution shape. Inset values report independent-samples t-statistics, p-values, and Cohen’s d for the age-group comparison. **(b)** Scatter plots showing the association between covariate-adjusted dispersion and the Emotional Resilience Index in younger adults (purple) and older adults (teal). Left: global dispersion. Right: FPN dispersion. Solid lines and shaded bands show the linear fit and 95% confidence interval per group, and each dot represents one participant. Network abbreviations: FPN, frontoparietal network.

At the network level, we examined within-network cortical dispersion separately for each of the seven functional networks^36^. For the right LC–linked within-network cortical dispersion, older adults showed significantly higher within FPN dispersion than younger adults during negative movie-viewing (**Figure 5**; *t* = −2.82, *p* = 0.006, *d* = 0.48; 95% CI [−0.117, −0.021]), which survived Bonferroni correction across the seven networks (*p_adj_* < 0.05). The left LC–linked FPN dispersion showed a directionally consistent but non-significant trend (*t* = −1.70, *p* = 0.091, *d* = 0.29, 95% CI [−0.136, 0.010]). At the uncorrected threshold, additional age-related effects were observed for the right LC–linked visual (VIS; *t* = −2.41, *p* = 0.017, *d* = 0.41), ventral attention/salience (VAN; *t* = −2.45, *p* = 0.015, *d* = 0.41), and limbic (LIM; *t* = −2.51, *p* = 0.013, *d* = 0.42) networks, while no significant age differences were observed in the DMN, SMN, or DAN (all *p*s > 0.05).

Between-network dispersion was also significantly higher in older adults for the right LC during negative movie-viewing (**Figure 5**; *t* = −2.42, *p* = 0.017, *d* = 0.41; 95% CI [−0.066, −0.007]), with the left LC–linked between-network dispersion showing a non-significant effect mirroring the pattern seen for global and within-network dispersion (*t* = −1.44, *p* = 0.152, *d* = 0.24; 95% CI [−0.090, 0.014]).

No age effects were detected during neutral movie-viewing for any dispersion measure or LC side (**Supplementary Figure 13**), suggesting that age-related differences in LC–cortical organization emerge specifically under conditions of heightened arousal and negative emotional valence.

To complement the dispersion analyses, we computed the subject-level range (max − min) of parcel gradient scores along each axis, which captures the distance between the cortical regions with the most dissimilar LC–connectivity profiles or in simple words occupying opposite ends of the gradient. Older adults showed a significantly wider G1 range than younger adults for the right LC during negative movie-viewing (*t* = −2.94, *p* = 0.004, *d* = 0.50), with a directionally consistent but non-significant effect on the left LC (*t* = −1.51, *p* = 0.133, *d* = 0.26). Right LC G2 range was also greater in older adults during the negative condition (*t* = −1.96, *p* = 0.052, *d* = 0.34). Under the neutral condition, age-related range differences were attenuated and largely non-significant (**Supplementary Figure 14**). This converges with the dispersion findings, indicating that cortical regions become more heterogeneous in their LC connectivity profiles with age along both the primary (G1) and secondary (G2) gradient axes, specifically during negative movie-viewing.

### Higher dispersion links with worse emotional wellbeing in aging

Participants completed a comprehensive cognitive and emotional assessment battery. Principal component analysis of standardized multiple measures of emotional wellbeing and cognitive task performance yielded two composite indices^38^: an Emotional Resilience Index (ERI) and a Cognitive Functioning Index (see Methods) with higher scores reflecting better emotional wellbeing and intact cognitive function. Given that significant age-related differences in dispersion were selectively observed for right LC–linked cortical organization, we next examined whether these specific alterations carry behavioral relevance by testing age × dispersion interactions on ERI scores. A significant interaction between age and right LC–linked cortical global dispersion was observed (**Figure 5***; β* = 0.62, *p* = 0.018), indicating that age moderates the relationship between global dispersion and emotional resilience index. Follow-up permutation-based correlation analyses (10,000 permutations) showed that higher right LC–linked cortical global dispersion was associated with lower scores in ERI among older adults (**Figure 5***; r* = −0.250, *p_perm_* = 0.042, *95% CI* = [–0.461, –0.012]), whereas no significant association was observed among younger adults (*r* = 0.195, *p_perm_* = 0.106, *95% CI* = [−0.039, 0.408]).

A parallel age × right LC–linked within-network FPN dispersion interaction was observed (**Figure 5**; *β* = 0.46, *p* = 0.037). Permutation tests confirmed that higher right LC–linked FPN dispersion was significantly associated with lower ERI scores among older adults (**Figure 5***; r* = −0.258, *p_perm_* = 0.035, *95% CI* = [−0.468, −0.021]) with no significant association among younger adults (*r* = 0.159, *p_perm_* = 0.186, *95% CI* = [– 0.076, 0.376]). We additionally tested moderation effects for the other networks that showed uncorrected age-related dispersion differences but did not survive Bonferroni correction (VIS, VAN, LIM). Age × dispersion interactions reached significance in the VIS (*β* = 0.47, *p* = 0.033) and LIM (*β* = 0.61, *p* = 0.029) networks, suggesting broader age-dependent moderation between right LC–linked network dispersion and ERI. However, follow-up permutation-based correlations did not reveal significant within-group associations in either network (VIS: younger adults *r* = 0.192, *p_perm_*= 0.139; older adults *r* = −0.139, *p_perm_*= 0.276; LIM: younger adults *r* = 0.199, *p_perm_*= 0.093; older adults *r* = −0.201, *p_perm_*= 0.104).

Associations between right LC–linked cortical dispersion and cognitive functioning index scores were limited. In older adults, higher dispersion within the VIS was associated with lower cognitive functioning index scores (*r* = −0.31, *p_perm_* = 0.013, *95% CI* = [−0.513, −0.073]). This association was not observed in younger adults (*r* = 0.002, *p* = 0.98, *95% CI* = [−0.23, 0.24]), and the age × dispersion interaction was not significant (*p* > 0.05). No other measures of dispersion i.e., global, within-network, or between-network were significantly associated with cognitive functioning index scores in either age group (all *ps* > 0.05).

### Validation and Sensitivity Analyses

To assess the robustness and specificity of our findings, we conducted a series of validation and sensitivity analyses, organized into four categories: (1) methodological robustness (seed definition, parcellation scheme), (2) gradient reliability (split-half, 5-fold cross-validation, leave-one-out individual-to-group correspondence, and inter-individual variability), (3) dispersion-specific sensitivity (gradient dimensionality, high-dispersion exclusion), and (4) confound controls (neuroanatomical specificity, physiological and visual artifacts).

#### Bilateral LC seed

Replicating the gradient analyses using a combined bilateral LC seed yielded topographies highly consistent with hemisphere-specific solutions (**Supplementary Figure 15;** G1 negative: left LC *ρ* = 0.94, right LC *ρ* = 0.97; G2 negative: left LC *ρ* = 0.98, right LC *ρ* = 0.93; G1 neutral: left LC *ρ* = 0.95, right LC *ρ* = 0.97). Within-age-group correlations during negative movie-viewing also remained high (**Supplementary Figure 16;** *ρ* = 0.85–0.98 across all gradients × hemisphere combinations), confirming that the gradient architecture is robust to seed definition.

#### Parcellation replications

Primary analyses were repeated using an independent 980-region cortical parcellation^49^. Parcel-wise Spearman correlations between the two parcellation-derived maps during negative movie-viewing were high for both G1 (right LC *ρ* = 0.97; left LC *ρ* = 0.98) and G2 (right LC *ρ* = 0.96; left LC *ρ* = 0.95; **Supplementary Figure 17**). To further assess replicability across parcel resolution and parcellation type, we repeated the analysis using the Schaefer-400 functional parcellation^39^ and the structurally defined Destrieux atlas^50^. Gradients derived from the Schaefer-400 parcellation closely aligned with Schaefer-1000 (G1: right LC *ρ* = 0.95, left LC *ρ* = 0.96; G2: right LC *ρ* = 0.96, left LC *ρ* = 0.97). Moreover, the primary gradient was reliably recovered with the Destrieux atlas during both negative (right LC *ρ* = 0.82, left LC *ρ* = 0.86) and neutral movie-viewing (right LC *ρ* = 0.83, left LC *ρ* = 0.85), demonstrating that the observed gradient organization was robust across parcellation scheme, resolution, and modality.

#### Reliability and inter-individual consistency

We evaluated gradient reliability using split-half (1,000 iterations) and 5-fold (200 iterations) cross-validation for the full sample (*N* = 140), younger adults (*n* = 72), and older adults (*n* = 68) across all four LC × condition combinations. Gradients showed high reliability in the combined sample (**Supplementary Figure 18;** right LC–cortex G1: split-half *r* = 0.88, 5-fold *r* = 0.83; right LC–cortex G2: split-half *r* = 0.92, 5-fold *r* = 0.88; and left LC–cortex G1: split-half *r* = 0.89, 5-fold *r* = 0.84; left LC–cortex G2: split-half *r* = 0.92, 5-fold *r* = 0.88). Younger adults consistently showed greater reliability than older adults during negative movie-viewing (younger adults right LC–cortex: split-half G1 *r* = 0.90, G2 *r* = 0.92; left LC– cortex: G1 *r* = 0.88, G2 *r* = 0.88; 5-fold: right LC–cortex G1 *r* = 0.85, G2 *r* = 0.89; left LC–cortex: G1 *r* = 0.82, G2 *r* = 0.83; older adults right LC–cortex: split-half G1 *r* = 0.80, G2 *r* = 0.82; left LC–cortex: G1 *r* = 0.80, G2 *r* = 0.81; 5-fold: right LC–cortex G1 *r* = 0.73, G2 *r* = 0.75; left LC–cortex: G1 *r* = 0.72, G2 *r* = 0.74; **Supplementary Table 4**).

To further assess gradient reliability at the individual level, we computed leave-one-out (LOO) spatial Spearman correlations between each participant’s parcel-wise gradient vector and the group-mean gradient derived from all the remaining participants. Using the full-sample as the reference level, mean correspondences ranged from *ρ* = 0.37–0.49. Similar levels of correspondence were observed within age groups, with younger adults showing *ρ* = 0.36–0.54 and older adults *ρ* = 0.37– 0.47, indicating reliable recovery of the LC–linked cortical gradients at the individual-level in both groups. Correspondence was generally similar in both movie-viewing conditions (**Supplementary Figure 19**; right LC G2: younger *ρ* = 0.52 neutral vs 0.54 negative; older *ρ* = 0.45 neutral vs 0.39 negative; **Supplementary Table 5**). Importantly, individual gradient reliability was not associated with dispersion in either age group (**Supplementary Table 6**), and the age-group differences in dispersion remained significant after additionally adjusting for G1 and G2 reliability (**Supplementary Table 7**), indicating that greater dispersion in older adults reflects a genuine difference in cortical organization rather than noisier gradient estimation. The age × dispersion interactions on emotional resilience were likewise unaffected by this adjustment. Specifically, significant age × dispersion interactions were observed for both global dispersion (β = 0.62, *p* = 0.018) and FPN dispersion (β = 0.46, *p* = 0.037). These effects remained significant after inclusion of the additional leave-one-out (LOO) covariates (global: β = 0.61, *p* = 0.021; FPN: β = 0.45, p = 0.045; **Supplementary Table 8**).

Pairwise Spearman correlations between individual gradient vectors were positive across all conditions and LC seeds (G1: *ρ* = 0.14–0.24; G2: *ρ* = 0.14–0.29), reflecting substantial but expected inter-individual variability in functional connectivity organization^51^. Younger-adult pairs exhibited greater similarity than older-adult pairs, during negative movie-viewing (**Supplementary Figure 20;** right LC G1: *ρ* = 0.23 vs 0.14; G2: *ρ* = 0.29 vs 0.18; **Supplementary Table 4**).

Together, these analyses indicate that the LC–linked cortical gradient topographies are reproducible and present at the individual level in both age groups. A key question for future work is whether the inter-individual variability observed, especially during negative movie-viewing in older adults is meaningfully linked to individual differences in cognition, mental health, and physiology (e.g., heart rate variability).

#### Dispersion-specific sensitivity

To verify that the group-level gradient is not dominated by high-dispersion individuals, we recomputed gradients after excluding participants with the top 5% (*n* = 7) and top 10% (*n* = 14) of global dispersion scores (all older adults) for the right LC–linked cortical gradient during negative movie-viewing. Spatial correlations between full-sample and excluded-sample gradients remained uniformly high (**Supplementary Table 9**), indicating that the reported gradient was not dominated by extreme individuals.

#### Gradient dimensionality

We retained the first two gradients for all primary analyses for two reasons. First, G1 and G2 together explained 68–79% of variance across conditions and hemispheres, whereas G3 consistently accounted for less than 10% (7–9%) which is below the variance level commonly used to guide gradient retention in the cortical-gradient literature^25,52,53^. Second, gradients explaining relatively less variance tend to have lower reliability and weaker predictive validity^54^, placing G3 within a less-informative range. Importantly, when repeating all dispersion analyses (global, within-network, between-network) using a three-gradient solution, the same pattern of results was observed with modestly attenuated effect sizes (right LC: global *d* = 0.41, between-network *d* = 0.35, FPN *d* = 0.38; **Supplementary Table 10 and Supplementary Table 11**).

#### Confound control

To verify the neuroanatomical specificity of the dominance analysis, we replaced the LC with the 10 × 10 voxel pontine tegmentum (PT), a neighboring brainstem structure frequently used as a reference for assessing LC structural integrity^55–60^. The PT explained only a negligible proportion of variance (adjusted R² = 0.07), confirming that the chemoarchitectural associations identified are LC specific. We extended the same control analysis to the age comparison. Cortical dispersion computed relative to the PT seed showed no age-group differences for any metric (**Supplementary figure 21**). Additionally, given the LC’s proximity to the fourth ventricle and the potential influence of physiological noise from cardiac and respiratory pulsations, we performed a post-hoc control analysis examining correlations between left and right LC signals and fourth ventricle signal fluctuations. No significant correlations were observed (*r* < 0.01), ruling out contamination from ventricular artifacts and supporting the physiological validity of the LC signal used in our analyses. Lastly, to assess the potential influence of low-level visual confounds, we tested the associations between luminance and LC–cortex functional connectivity and observed no reliable associations in any movie, LC hemisphere, or age group (**Supplementary Tables 12 and 13**). This is consistent with prior evidence that the LC is preferentially activated by arousal and emotional salience rather than by ambient luminance per se^61–63^ (**see Supplementary Methods for luminance extraction methods**).

## Discussion

We identified two dominant gradients that provide a low-dimensional representation of cortical organization through the lens of LC connectivity. The primary gradient remained consistent across both movie-viewing conditions and tracked the cortical distribution of neurotransmitter receptors and transporters, linking it to the brain’s underlying chemoarchitecture, whereas the secondary gradient aligned with canonical functional axes^23^ in a context-dependent manner. This organization was particularly sensitive to aging. During negative movie-viewing, older adults exhibited significantly higher global and network-specific dispersion, a hallmark of functional dedifferentiation. This reorganization was most prominent within the FPN and carried behavioral relevance, as higher FPN dispersion was associated with poorer emotional wellbeing in later life.

One major observation from this work is the association of the primary LC–linked cortical gradient with chemoarchitectural organization in both movie-viewing conditions. This finding extends prior work showing that macroscale cortical gradients align with the spatial distribution of neurotransmitter receptors and transporters^24,26,27^. Using dominance analysis, we found that dopaminergic and noradrenergic receptor– transporter densities explained the largest unique proportion of variance, identifying G1 as a catecholaminergic axis along which cortical regions differ in neuromodulatory profile. This interpretation is consistent with accumulating evidence that catecholaminergic receptor–transporter distributions shape cortical gain, responsiveness to neuromodulatory input, and communication across large-scale cortical networks^64–67^. Notably, the spatial polarity of G1, positively associated with NAT density and negatively associated with DAT density, mirrors established catecholaminergic gradients across the cortex. NAT-dominant regions, particularly frontal and somatomotor cortices^68,69^, are positioned to sustain LC influence, as catecholamine clearance in these regions relies primarily on NAT, which exhibits slower reuptake and lower affinity than DAT^70,71^. As a result, NA and dopamine co-released from LC terminals persist longer in the extracellular space^72^, supporting prolonged neuromodulatory effects. Critically, G1 remained stable across both movie-viewing conditions, reflecting a stable, chemoarchitecturally anchored axis within the low-dimensional manifold of LC–cortical connectivity. Supporting its specificity, a neighboring PT control region did not show comparable explained variance in the dominance analysis, confirming that this organization is LC-specific.

In contrast, G2 exhibited condition-dependent organization. During neutral movie-viewing, LC–cortical coupling was organized along the canonical V–S axis, consistent with engagement of perceptual systems under low emotional demand. During negative movie-viewing, however, G2 reflected the canonical S–A axis, indicating stronger LC coupling with higher-order associative networks. These findings are consistent with models in which increased noradrenergic tone elevates cortical gain and biases processing toward networks that support appraisal, regulation, and adaptive behavior^5,73,74^. These results indicate that LC–linked cortical organization comprises a stable axis anchored in cortical chemoarchitecture and a flexible secondary axis that aligns with distinct canonical axes in the two emotional movie-viewing conditions. This organization emerged only when LC–cortical connectivity was probed across emotional contexts, suggesting that the low-dimensional manifold of neuromodulatory connectivity to cortex may dynamically reorganize with emotional salience. Such dynamics may therefore be best characterized through naturalistic paradigms. Further, the within-LC gradient aligned with the rostro-caudal axis replicating prior findings^17^. This provides independent confirmation that our gradient approach recovers established functional organization.

Beyond these organizational principles, a marked reorganization within the LC–linked cortical manifold was observed in late life. Older adults showed a significantly higher global dispersion, indicating that cortical regions were more widely spread across gradient space. In other words, regions that formed relatively compact clusters in younger adults occupied a more diffuse, less cohesive arrangement in older adults, consistent with age-related functional dedifferentiation^25,28–30^. Importantly, the overall gradient topography was preserved across age groups, with parcels occupying broadly similar positions along each gradient in younger and older adults^25,31^. Thus, what changed with age was not where regions sit in the manifold, but how widely they spread. The observation that age-related effects emerged selectively during negative movie-viewing suggests that emotionally salient contexts can amplify age-related differences in LC–linked cortical organization. This condition-dependent effect aligns with arousal-biased competition^75^ and gain-modulation models^5,6^, which posit that heightened arousal increases the influence of LC signaling and, in turn, cortical gain. In younger adults, such modulation may facilitate coordinated LC–cortical coupling. In older adults, by contrast, age-related reductions in neuromodulatory precision^33–35^, combined with shifts toward elevated tonic and reduced phasic LC activity as observed in aging animal models^76^, may produce more diffuse, less selective neuromodulatory signaling to the cortex. Notably, these age-related effects emerged only under an emotionally salient condition and would have been obscured in a resting-state design. Emotional arousal may therefore serve as a functional probe that unmasks instability in LC–linked cortical organization in later life.

Network-level analyses further localized these effects. Although several networks exhibited elevated within-network dispersion in older adults, only the FPN survived correction for multiple comparisons. This finding is notable given the FPN’s relatively high noradrenergic receptor expression ^77,78^ and evidence from rodent studies demonstrating FPN sensitivity to LC-mediated gain modulation under cognitively demanding conditions^79^. While translating rodent findings to human functional networks requires caution, the convergence across species supports the interpretation that the FPN may be particularly vulnerable to age-related changes in LC neuromodulation^21,80^. Our results show that LC–linked FPN organization becomes more diffuse in older adults. This pattern is consistent with neuromodulatory aging frameworks proposing that reduced receptor–transporter specificity may compromise the LC’s ability to deliver selective gain modulation to distributed control circuits^33,35^, resulting in more variable connectivity. Future studies combining LC–cortical gradient analyses with individual-level PET measures of receptor and transporter availability, or longitudinal multimodal designs, will be critical for directly testing this proposed mechanism.

Notably, higher age-related dispersion was more pronounced in the right LC–linked cortical organization. A direct test of the age × hemisphere interaction did not support a reliable hemispheric difference (**Supplementary Table 14**); the effect is therefore best described as bilateral with a right-sided emphasis. One possible explanation, which we offer as a hypothesis to be tested in larger samples, is that the asymmetry reflects the specific demands of the current paradigm, in which participants observed another individual experiencing acute distress without direct personal threat (a form of vicarious emotional processing) that recent rodent work has linked to preferential right-LC engagement^81^. This right-lateralization is thought to arise not from the LC itself but from asymmetric afferent input via the bed nucleus of the stria terminalis, which selectively mediates vicarious fear^81^. The LC–NA system may then further regulate fear responses by modulating theta oscillations in cingulate–amygdala circuits^82^. While prior human work has emphasized left-LC involvement in introspective and resting-state contexts^17,83^, our findings are consistent with, and extend, prior work by pointing to a right-sided emphasis of age-related differences in LC–linked cortical dispersion during observational emotional processing. Future work combining neuroimaging with electrophysiology could test whether the left and right LC are differentially recruited during personal versus vicarious emotional processing, and whether lateralized oscillatory dynamics can explain the age effects we observed.

Importantly, this reorganization in late life was behaviorally meaningful. Among older adults, higher global and LC–linked FPN dispersion during negative movie-viewing was associated with lower self-reported emotional wellbeing, suggesting that a more diffuse LC–linked cortical organization, particularly within the FPN, may compromise emotional regulation networks precisely when emotional demands are greatest. This observation is consistent with recent findings linking higher FPN dispersion to emotion-regulation difficulties and vulnerability to mental health difficulties in aging^32^. Several lines of neurobiological evidence provide a plausible framework for this interpretation. For instance, studies in non-human primates demonstrate that excessive or dysregulated NA activity destabilizes prefrontal networks by biasing signaling from high-affinity α2A receptors to lower-affinity α1 and β receptors, weakening persistent activity and top-down control^84,85^. Age-related alterations in LC firing dynamics may further exacerbate this vulnerability^76^, reducing the capacity of frontoparietal systems to regulate emotional responses effectively^32^. These findings align with broader theories of age-related dedifferentiation, which posit that reduced neural specificity contributes to declines in cognitive performance^25,30,86^. Extending these frameworks, our results suggest that LC–linked global and particularly FPN dedifferentiation may reflect diminished neuromodulatory precision that undermines emotional wellbeing under emotionally demanding affective conditions in later life.

Several limitations should be noted. First, analyses were restricted to cortex. Although the LC also innervates subcortical structures^2^, the availability of well-established cortical gradients and chemoarchitectural maps provides a principled foundation for characterizing large-scale LC–cortical connectivity organization. This provides a necessary first step toward extending the framework to subcortical systems. Second, each valence condition was represented by a single movie clip. Although the neutral and negative clips were matched on duration, character composition, and absence of dialogue through a pilot study^87^, clip-specific semantic influences cannot be completely ruled out. Future work could employ multiple exemplars per valence category to strengthen stimulus generalizability. Third, LC measurements are susceptible to physiological artifacts due to the LC’s proximity to the fourth ventricle. Nonetheless, our post-hoc analyses revealed no significant correlation between LC activity and fourth-ventricle fluctuations, supporting the robustness of our findings. Fourth, the fixed presentation order of the movie clips, neutral followed by negative, was chosen to prevent emotional carry-over from the negative film contaminating the neutral condition. However, condition differences may still not be fully disentangled from time-in-scanner effects, such as scanner drift, progressive head motion, and fatigue, which may affect older adults. Although dispersion correlated modestly with head motion (Global dispersion: *r* = 0.16 overall, *r* = 0.22 in older adults; between-network dispersion: *r* = 0.17 overall, *r* = 0.34 in older adults; FPN dispersion: *r* = 0.14 overall, *r* = 0.24 in older adults**; Supplementary Table 15**), the age-related effects in dispersion and their associations with emotional wellbeing remained significant after controlling for head motion, suggesting that the reported effects are unlikely to be explained by motion-related artifacts.

Fifth, our cross-sectional design precludes inferences about longitudinal change. Whether the observed age differences reflect developmental trajectories, cohort effects, or both remains uncertain. Longitudinal studies tracking LC–cortical connectivity across the adult lifespan will be essential for distinguishing these possibilities.

In summary, this study demonstrates that LC-cortical organization is tracked by the brain’s molecular and canonical functional architecture and becomes increasingly dispersed with age. Emotional arousal selectively unmasked this reorganization, most prominently within the FPN, with implications for emotional wellbeing in later life. These findings extend prevailing theories of neural dedifferentiation and neuromodulatory aging, indicating that age-related declines in neuromodulatory precision may propagate to the LC-cortical manifold. Critically, our work advances beyond the current state of the art. Whereas existing accounts treat locus coeruleus decline and cortical dedifferentiation as separate processes and rely largely on static structural or resting-state measures, we show that they are interrelated, and that negative-valence, high-arousal movie-viewing unmasks this age-related reorganization otherwise invisible at rest. More broadly, by positioning LC-linked FPN organization as a potential neural substrate connecting noradrenergic aging to emotional vulnerability, our results point to new targets for the early detection and prevention of mental health difficulties in later life.

## Materials and Methods

The current study used data from the Trondheim Aging Brain Study (TABS). The study design and procedures have been described in detail^32,38,87^. We have added a detailed description in the **Supplementary Methods** outlining how the present study differs from and extends the prior TABS-based publications. Here, we include the materials and methods relevant to the current study. TABS was approved by the regional Ethics Committee NTNU, Norway (Midt-REC). All participants provided written informed consent prior to testing.

### Participants

Eighty younger and 75 older adults were scanned as part of TABS and completed two separate sessions: one imaging and one behavioral-neuropsychological testing sessions. However, due to incomplete behavioral datasets, technical issues during data collection, order of stimuli reversed and excessive head motion^88^ (mean framewise displacement ≥0.3 mm), the final sample in the current study included a total of 140 participants (72 Younger and 68 Older adults). Sex was determined by self-report and was approximately equally divided within each age group (younger: 34 females of 72; older: 36 females of 68). Sex differences are reported in **Supplementary Figure 22**.

Participants were recruited from the local community through paper flyers and online advertisements. Exclusion criteria included the presence of metallic implants, claustrophobia, or a history of neurological disorder (e.g., neurodegenerative diseases), psychiatric disorders, or autism spectrum disorder. All participants were right-handed, had normal or corrected-to-normal vision, and reported normal hearing abilities. All older participants underwent cognitive screening using the Mini-Mental State Examination^89^ (MMSE), conducted either in person or via phone using the 22-point Braztel-MMSE^90^, and participants scored above the respective thresholds of 24 and 15, confirming lack of cognitive deficits. Specifically, MMSE scores ranged from 25 to 30 (*mean* = 29.05, SD = 1.22), and telephonic Braztel-MMSE scores ranged from 17 to 22 (*mean* = 21.34, SD = 0.98). All participants received compensation of 500 Norwegian Kroner in gift cards for their participation.

### Movie clips

During fMRI, participants watched two clips: a neutral clip and a negative-valence clip. The neutral movie-viewing ‘Pottery’ (480 seconds) depicts a woman making pottery and was rated as low negative valence and low arousal in a pilot study^87^. The negative clip ‘Curve’ (494 seconds) shows a woman struggling to avoid falling from a cliff and was rated high in negative valence and arousal. The clips were selected from a pilot study in which participants rated four candidate videos on valence and arousal; Pottery and Curve were chosen to represent neutral and negative conditions, respectively. Prior work from our group shows that Curve reliably elicits high arousal and negative valence self-reported responses, whereas Pottery functions as an emotionally minimal control^32,87^. To reduce confounds across clips, we matched key features: (1) one human character appears on screen at a time; (2) all characters are female; (3) no dialogue, written text, or subtitles; and (4) both clips are in color.

### Neuropsychological assessments

During the behavioral testing session, participants completed self-report questionnaires assessing mental health and emotion-related functioning: Depression, Anxiety, and Stress Scale^91^ (DASS-21), the Difficulties in Emotion Regulation Scale^92^, the General Health Questionnaire^93^, the Hospital Anxiety and Depression Scale^94^, the Connor-Davidson Resilience Scale^95^, the Intolerance of Uncertainty Scale^96,97^, the Perceived Stress Scale^98^, and the State-Trait Anxiety Inventory^99^.

Cognitive performance was assessed using Stroop test^100^, the Trail Making Test^101^, and the COWAT Phonemic and Semantic Fluency Task^102^, and forward and backward digit span tests. As multiple emotional and cognitive measures were collected, we applied principal component analysis (PCA) to reduce dimensionality and derived two indices: an Emotional Resilience Index (ERI) capturing emotional resilience and mental wellbeing, and a Cognitive Functioning Index capturing overall cognitive performance.

### Procedure

Before scanning, participants received written and verbal instructions. The MRI session began with an ∼8-min anatomical scan, followed by movie-fMRI. The neutral clip always preceded the negative clip. Participants rated valence (1 = very unpleasant to 9 = very pleasant) and arousal (1 = very calm to 9 = very excited) using the 9-point Self-Assessment Manikin before and after each clip^103^; they also rated baseline emotions before entering the scanner. Participants were instructed to watch the videos naturally while minimizing movement; foam padding stabilized the head. Audio was delivered via MRI-compatible earphones (BOLDfonic, Cambridge Research Systems Ltd), with volume adjusted individually. The behavioral session was completed within the same week, typically 3–4 days from scanning session.

### Imaging acquisition and preprocessing

Data were acquired on a 7 T Siemens MAGNETOM Terra scanner with a 32-channel head coil at the Norwegian 7 T MR Center (St. Olav’s Hospital). High-resolution anatomical T1-weighted images were collected using MP2RAGE^104^ (voxel size 0.75 × 0.75 × 0.75 mm; 224 slices; TR = 4300 ms; TE = 1.99 ms; flip angles = 5°/6°; field-of-view (FOV) = 240 × 240 × 175 mm; slice thickness = 0.75 mm). To achieve high-resolution imaging of the LC, we acquired magnetization transfer-weighted turbo flash images^105^ (MT-TFL; voxel size 0.4 × 0.4 × 0.5 mm; 60 slices; TR = 400 ms; TE = 2.55 ms; flip angle = 8°). The MT-TFL FOV was positioned approximately perpendicular to the pons, covering the region between the inferior colliculus and the inferior border of the pons. Functional data were acquired using multiband EPI (92 interleaved slices; multiband factor = 2; voxel size = 1.25 × 1.25 × 1.25 mm; TR = 2000 ms; TE = 19 ms; matrix = 160 × 160; FOV = 200 mm; slice thickness = 1.25 mm; flip angle = 80°), yielding 243 volumes (neutral movie) and 250 volumes (negative movie).

Preprocessing used fMRIPrep v22.0.2^106^, followed by postprocessing in XCP-D v0.3.053^107^. Steps included removal of the first three volumes, motion and slice-timing correction, co-registration, normalization to MNI space, nuisance regression, and band-pass filtering (details in Supplementary Methods). Due to small size of the LC, smoothing was performed with a 2 mm full width at half maximum Gaussian kernel, following the recommendation to use a kernel of at least 1.5 times the native voxel size^108^, maintaining spatial precision to avoid signal blurring^109^.

### Locus coeruleus delineation and spatial transformation

Detailed procedures for LC delineation and spatial transformation are described in previously published work^38^ and LC segmentation followed the established published protocol^58,59^. Briefly, MT-TFL images were intensity-normalized by dividing voxel values by the mean signal from a 10 × 10 pontine tegmentum (PT) ROI on the slice containing the LC peak voxel, reducing inter-subject variability while preserving LC– PT contrast^55^. A group template was created from normalized MT-TFL images using Greedy SyN registration with cross-correlation^110,111^. LC was manually segmented on the template in ITK-SNAP based on voxel intensity and anatomical landmarks, repeated after 4 weeks, and thresholded to retain overlapping voxels (Dice: left LC = 0.959; right LC = 0.962).

For spatial transformation, MT-TFL images and LC segmentations were resampled to isotropic T1w resolution using FreeSurfer mri_convert (v7.1). MT-TFL images were rigidly registered to native T1w space using antsRegistrationSyN^110^. T1w images were then nonlinearly registered to MNI152 space using SyN^110,111^, and the T1w image served as an intermediate bridge for transforming MT-TFL data into MNI space. Transformations were combined and applied to LC masks using antsApplyTransforms with nearest-neighbor interpolation to preserve voxel boundaries, minimizing interpolation error and maintaining LC localization in MNI space.

### Locus coeruleus-cortical gradient analysis

For each participant, movie-viewing condition (neutral and negative), and LC side (left and right LC), we constructed an LC–cortical functional connectivity matrix. LC voxel-wise time series were extracted from study-specific LC masks delineated on the MT-TFL sequence (**Supplementary Figure 2**), and cortical time series were extracted from 1,000 parcels of the Schaefer atlas^39^ spanning seven canonical functional networks^36^. Pairwise Pearson correlations between each LC voxel and cortical parcel were Fisher z-transformed, yielding a participant-level LC-voxel × cortical-parcel connectivity matrix. For each movie-viewing condition, individual connectivity matrix was averaged across participants to form a group-mean LC–cortical matrix. Using the BrainSpace toolbox^40^, a cortex × cortex affinity matrix was computed using normalized angular (cosine) similarity between cortical parcels based on their LC connectivity; this quantifies how similarly cortical regions are coupled to the LC. Diffusion map embedding^41^ was then applied to this affinity matrix to derive low-dimensional gradient components capturing the principal axes of variation in LC–cortical connectivity. The manifold-learning parameter α was set to 0.5, consistent with prior gradient-mapping work^23,40,112^, balancing the contribution of local versus global connectivity density to the embedding geometry. The resulting components served as the condition-specific group reference template. Individual-participant gradients were estimated using the same procedure and aligned to the corresponding group template via Procrustes rotation to ensure cross-subject comparability. Aligned individual gradients were then averaged to obtain a group-level gradient map across all participants, and separately for younger and older adults. Gradients were ranked by their explained variance. To characterize functional organization within the LC itself, we transposed the LC-voxel × cortical-parcel matrix and computed normalized angular (cosine) similarity between LC voxels based on their cortical connectivity profiles. Diffusion map embedding was then applied to the resulting affinity matrix. All gradient estimation procedures were implemented in MATLAB using the BrainSpace toolbox^40^.

### Age-related differences in dispersion

To quantify age-related differences in LC–linked cortical organization, we computed dispersion in the two-dimensional gradient space (G1-G2), adapting the framework introduced in previous study^25,48^. All dispersion measures were calculated at the individual level within Procrustes-aligned gradient spaces on the unnormalized gradient embeddings to preserve the variance structure of the decomposition.

Global dispersion was defined as mean Euclidean distance of all cortical parcels to the global centroid, reflecting whole-cortex spread.

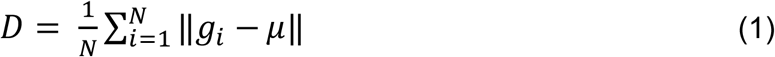

Within-network dispersion was calculated as the mean Euclidean distance of parcels within a given network from that network’s centroid.

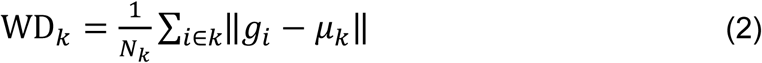

Between-network dispersion was computed as the Euclidean distance between pairs of network centroids. Because cortical atrophy may affect functional embedding, cortical thickness was included as a covariate^25^.

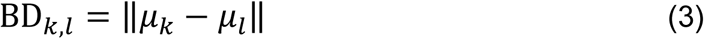

*g_i_*, *Coordinates of ROI i in the 2-D gradient space (Gradient 1 and Gradient 2)*.

*μ*, *global centroid across all parcels in 2D gradient space. μ_k_*, *centroid of network k in gradient space*

Linear regression models also adjusted for sex and head motion. To examine whether effects were specific to a gradient dimension, we also computed within-network dispersion separately along G1 and G2. Additional measures included the numerical range of each gradient (min–max distance) and the standard deviation of eigenvector values across parcels (**Supplementary Figure 14**). Control analyses confirmed that LC–linked dispersion reflected both gradient axes, with G1 and G2 range each correlating with global dispersion within both age groups, indicating that dispersion captures spread along both axes rather than a single dominant one (**Supplementary Figure 23**). Association of global dispersion with within-network dispersion is presented in **Supplementary Table 16**.

### Meta-analytic functional decoding with Neurosynth

We used Neurosynth^113^ volumetric association test maps (https://github.com/neurosynth/neurosynth) to derive probabilistic associations between brain voxels and cognitive/affective terms based on large-scale coordinate-based meta-analysis (>14,000 studies). For each voxel, the association reflects the probability that a term is reported when activation occurs at that voxel. Terms were selected from the Cognitive Atlas ontology^114^, yielding 123 terms spanning broad constructs (e.g., “attention”) and specific processes (e.g., “visual attention”, “episodic memory”) as well as affective states (e.g., “fear”, “pain”). Neurosynth maps were parcellated using the Schaefer 1,000-parcel atlas^39^ and z-scored.

Statistical inference was corrected at two levels. First, to control for spatial autocorrelation^115,116^, significance was assessed using the Hungarian spin-permutation test^117^, yielding two-sided spin-corrected p-values (*p_spin_*). Second, because each gradient × condition × hemisphere combination tests ∼123 terms, we applied Benjamini–Hochberg FDR correction^118^ across terms within each combination (*q* < 0.05); full term lists with *p_spin_* and FDR-corrected values (*p_spin_* FDR) are reported in **Supplementary Tables 17–24**. For descriptive functional decoding, terms surviving *p_spin_* < 0.05 are visualized in **Extended Data Figure 1**, consistent with prior gradient studies^26,119^.

### Neurotransmitter receptors and transporters

Receptor and transporter density maps were obtained from JuSpace library^37^. We included 13 maps spanning major neuromodulatory systems (**Supplementary Figure 24**; **Supplementary Table 1)**: dopamine D1, D2 receptors^120,121^, dopamine transporter^122^, noradrenaline (noradrenaline transporter NAT^123^), serotonin^124,125^ (5-HT1a, 5-HT1b, 5-HT2a, 5-HT4 receptors; serotonin transporter, SERT), acetylcholine (vesicular acetylcholine transporter^126^, VAChT), glutamate (metabotropic glutamate receptor 5, mGluR5^26^), gamma-aminobutyric acid type A receptor^37^ (GABAa), and N-methyl-D-aspartate receptor^127^ (NMDA). Maps were parcellated using the Schaefer 1,000-parcel atlas^39^ to match the functional gradients ensuring direct comparability between molecular and functional spaces.

### Dominance analysis

To assess how LC–linked cortical gradient maps are related to neurotransmitter receptor and transporter distributions, we applied dominance analysis^47^. This method quantifies the relative contribution of each receptor and transporter density map to the fit (*adjusted R²*) of a multiple linear regression model. It does so by fitting models across all possible combinations of predictors (2^ᵖ^ − 1 submodels for *p* predictors). For each variable, total dominance is defined as the average increase in explained variance (R²) when that variable is added to each submodel, such that the sum of all predictors’ dominance equals the adjusted R² of the full model. This approach accounts for collinearity among predictors and provides interpretable estimates of relative importance. We also computed interactional dominance, defined as the incremental variance explained when adding a predictor to the model containing all other predictors, indexing unique variance in the presence of shared effects for negative (**Supplementary Figure 25)** and neutral movie-viewing (**Supplementary Figure 26**). Total and interactional dominance values were normalized by the full model fit (R²_adj_) to facilitate comparisons across analyses.

### Spatial null model

All spatial correspondence analyses between parcel-wise cortical maps including alignment of LC–linked cortical gradients with canonical functional gradients, Neurosynth association maps, condition-to-condition gradient comparisons, and hemisphere-to-hemisphere gradient comparisons used a consistent spatial-null pipeline. In brief, spatial associations were quantified using Spearman correlations, and significance was assessed against a spatial-autocorrelation-preserving null^115,116^ using the Hungarian spin test^45^ with 10,000 spins. Two-sided spin-corrected p-values (*p_spin_*) were computed as the proportion of null statistics equaling or exceeding the empirical statistic in absolute value, with 1 added to both numerator and denominator (10,001), giving a minimum possible value of *p_spin_* ≈ 0.0001.

## Acknowledgments

We would like to thank radiographers and MR physicists at the 7T MR center at NTNU for their help with this project. We also would like to thank our participants for their time and effort during the experiment. We thank Stian Framvik, Avneesh Jain, Jae Hong, Leona Rahel Bätz and Karina Tømmerdal for their help during data collection. We also thank Carol Barnes for careful reading of the manuscript and Martin Dahl, Lynn Nadel, and Mary Peterson for helpful discussions on the results. This project was supported by the Research Council of Norway through its Centers of Excellence scheme, project number 332640 and National Infrastructure grant from the Research Council of Norway (NORBRAIN, project number 245904/350201). We would like to thank Kavli Foundation and Hjerneforskningsfondet for their support.

## Data availability

PET-derived receptor–transporter density maps were obtained from the JuSpace toolbox (https://github.com/juryxy/JuSpace; Dukart et al., 2021). The Schaefer 1000-parcel cortical atlas was obtained from the Computational Brain Imaging Group repository (https://github.com/ThomasYeoLab/CBIG/tree/master/stable_projects/brain_parcellation/Schaefer2018_LocalGlobal; Schaefer et al., 2018). The Yeo 7-network functional parcellation was obtained from (https://surfer.nmr.mgh.harvard.edu/fswiki/CorticalParcellation_Yeo2011; Yeo et al., 2011). Neurosynth data are available at https://neurosynth.org/, and the Cognitive Atlas is available at https://www.cognitiveatlas.org/. Source data are publicly available through https://github.com/Ziaei-Group/Age-Differences-in-LC-linked-Cortical-Gradients. Raw neuroimaging data are not publicly available due to ethical and data-protection restrictions. The data will meanwhile remain available upon request from the corresponding last author (M.Z), subject to the necessary institutional and data-sharing agreements.

## Code availability

Gradient analyses were conducted using BrainSpace, an open-access toolbox for macroscale gradient mapping and analysis (https://github.com/MICA-MNI/BrainSpace). Code for computing gradient dispersion measures is available at https://github.com/rb643/GradientDispersion. Movie stimuli and custom analysis scripts are available through https://github.com/Ziaei-Group/Age-Differences-in-LC-linked-Cortical-Gradients

## Contributions

A.D.: conceptualization, methodology, formal analysis, investigation, writing—original draft. S.Y.: conceptualization, methodology, data curation, software, investigation, writing—review & editing. X.L.: formal analysis. A.S.: formal analysis, writing—review & editing. H.I.L.J.: methodology, writing—review & editing. M.Z.: conceptualization, methodology, funding acquisition, data curation, supervision, writing—review & editing.

---

**Extended Data Figure 1:**
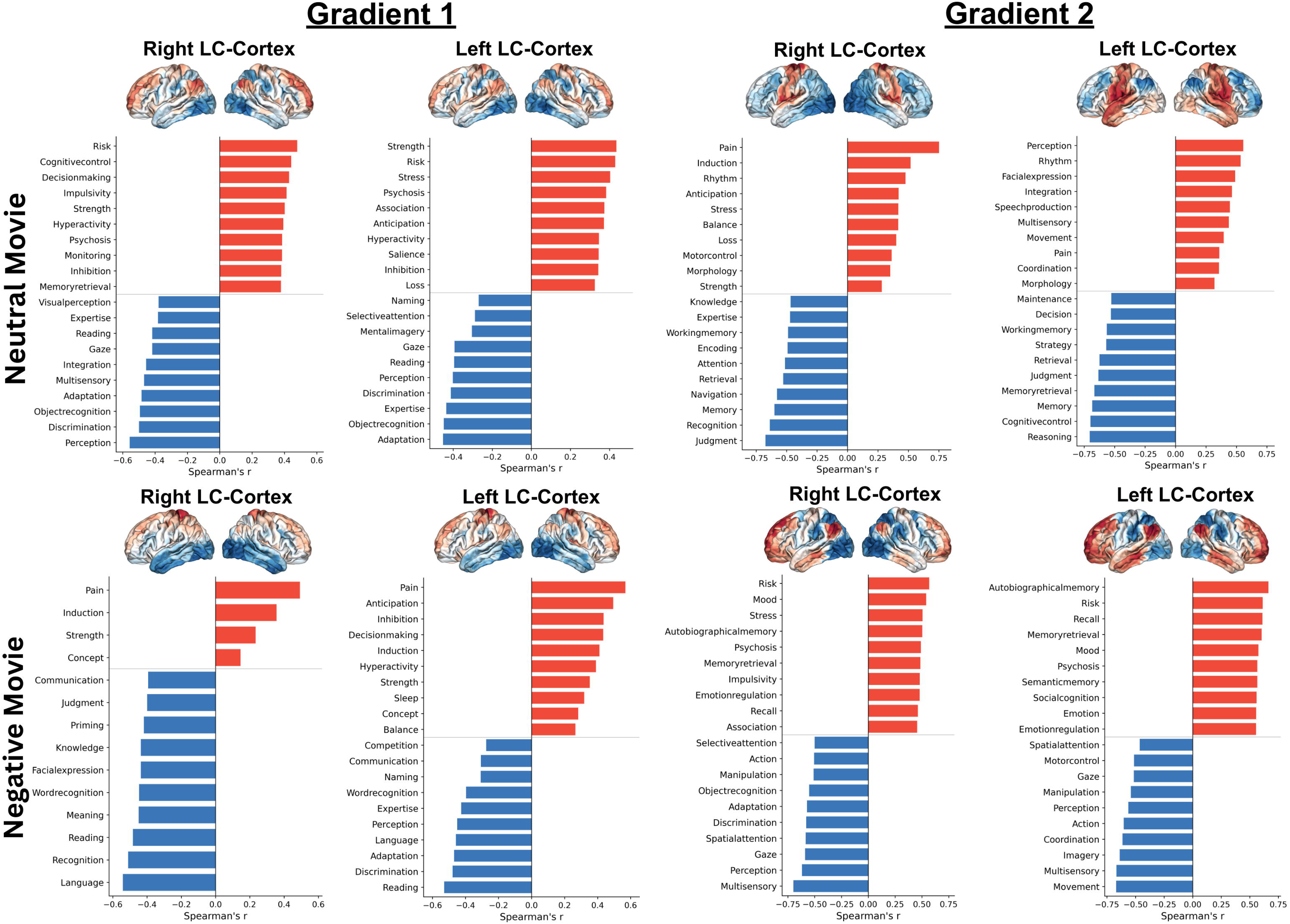
Neurosynth functional decoding of LC–linked cortical gradients 1 and 2 across emotional conditions and hemispheres. Cortical surface projections of LC–linked cortical gradients and their corresponding Neurosynth meta-analytic association maps are shown for Gradient 1 and Gradient 2, separately for the right and left LC, under neutral and negative movie-viewing conditions. Top row: neutral movie; bottom row: negative movie. Within each row, panels show, from left to right, Gradient 1 (right LC, left LC) and Gradient 2 (right LC, left LC). For each panel, inset cortical surfaces display the gradient map and bar plots show the top-ranked Neurosynth association terms ranked by Spearman’s ρ. Red bars indicate positive correlations between the gradient map and meta-analytic association maps; blue bars indicate negative correlations. Statistical significance was assessed against a spin-test null using the Hungarian method (10,000 spherical rotations with optimal parcel reassignment^117^), which preserves the empirical spatial autocorrelation of the cortex; only terms surviving p_spin_ < 0.05 are shown.

