## Supplementary document for "Age-related reorganization of locus coeruleus–cortical functional connectivity gradients"

**This file includes:**

Supplementary Methods

Supplementary Figures 1 to 26

Supplementary Tables 1 to 24

**Supplementary Information**

**Supplementary Methods**

**Methods**	Data Preprocessing and Postprocessing	3

**Supplementary TexT**

**Text**	Luminance Analysis	10

TABS Based Finding Summary	11

**Supplementary Figures**

**Figure 1**	Self-reported valence and arousal across movie conditions, by age group	13

**Figure 2**	Locus coeruleus delineation pipeline	14

**Figure 3**	Cross-condition correspondence of the primary gradient (G1)	15

**Figure 4**	Cross-hemisphere correspondence of the primary gradient (G1)	16

**Figure 5**	Cortical parcels expressed along Gradient 1	17

**Figure 6**	Alignment of G1 with canonical macroscale cortical axes	18

**Figure 7**	Correlation of noradrenaline and dopamine transporter density with G1	19

**Figure 8**	Topographic gradient of the LC based on cortical connectivity	20

**Figure 9**	Age-related network shifts in the 2D manifold (neutral movie)	21

**Figure 10**	Age-group correlations of LC–cortex gradient topographies	22

**Figure 11**	Schematic of the three dispersion measures	23

**Figure 12**	Empirical dispersion measures embedding	24

**Figure 13**	Age-related differences in cortical dispersion (neutral movie)	25

**Figure 14**	Age-related differences in per-axis gradient range, by hemisphere	26

**Figure 15**	Bilateral consistency of the primary and secondary gradients	27

**Figure 16**	Bilateral consistency of the gradients, by age group	27

**Figure 17**	LC–cortex connectivity patterns in 980 cortical parcels	28

**Figure 18**	Cross-validation reliability of LC–linked cortical gradients	29

**Figure 19**	Individual-to-group gradient similarity at the participant level	30

**Figure 20**	Inter-individual similarity of gradient organization	31

**Figure 21**	Cortical gradient dispersion relative to the pontine control seed between age groups	31

**Figure 22**	Sex differences in gradient dispersion	32

**Figure 23**	Per-axis gradient range and 2D dispersion	33

**Figure 24**	PET-derived receptor/transporter density maps	34

**Figure 25**	Interactional dominance results (negative movie)	35

**Figure 26**	Interactional dominance results (neutral movie)	36

**Supplementary Tables**

**Table 1**	Details of receptor–transporter density maps	37

**Table 2**	Age-related centroid displacement of Yeo networks (negative)	38

**Table 3**	Age-related centroid displacement of Yeo networks (neutral)	39

**Table 4**	Reliability of gradients across validation methods	40

**Table 5**	Leave-one-out individual-to-group gradient similarity	41

**Table 6**	Correlations between Right-LC dispersion and individual gradient reliability	43

**Table 7**	Age-group differences in right-LC dispersion with and without individual gradient reliability as a covariate	43

**Table 8**	Associations between right-LC dispersion and the Emotional Resilience Index with individual gradient reliability as an additional covariate	44

**Table 9**	Sensitivity: full- vs. excluded sample gradients	44

**Table 10**	Sensitivity: 2D vs. 3D dispersion metrics (right LC)	45

**Table 11**	Sensitivity: 2D vs. 3D dispersion metrics (left LC)	45

**Table 12**	LC–cortex FC correlation with framewise luminance	46

**Table 13**	LC–cortex FC correlation with luminance, by age group	46

**Table 14**	Age × hemisphere interaction for LC–cortical gradient dispersion during negative movie-viewing	47

**Table 15**	Correlations between right LC–cortical dispersion and mean FD during negative movie-viewing	47

**Table 16**	Within-network vs. global dispersion correlations (negative)	48

**Table 17**	Neurosynth decoding — left LC, G1, neutral movie	49

**Table 18**	Neurosynth decoding — left LC, G2, neutral movie	50

**Table 19**	Neurosynth decoding — right LC, G1, neutral movie	53

**Table 20**	Neurosynth decoding — right LC, G2, neutral movie	55

**Table 21**	Neurosynth decoding — left LC, G1, negative movie	57

**Table 22**	Neurosynth decoding — left LC, G2, negative movie	58

**Table 23**	Neurosynth decoding — right LC, G1, negative movie	61

**Table 24**	Neurosynth decoding — right LC, G2, negative movie	62

**Supplementary Methods**

**Imaging data preprocessing**

Results included in this manuscript come from preprocessing performed using *fMRIPrep* 22.0.2 (Esteban, Markiewicz, et al. (2018); Esteban, Blair, et al. (2018); RRID:SCR_016216), which is based on *Nipype* 1.8.5 (K. Gorgolewski et al. (2011); K. J. Gorgolewski et al. (2018); RRID:SCR_002502).

**Preprocessing of B0 inhomogeneity mappings**

A total of 2 fieldmaps were found available within the input BIDS structure. A B0-nonuniformity map (or fieldmap) was estimated based on two (or more) echo-planar imaging (EPI) references with topup (Andersson, Skare, and Ashburner (2003); FSL 6.0.5.1:57b01774).

**Anatomical data preprocessing**

A total of 1 T1-weighted (T1w) images were found within the BIDS dataset input. The T1-weighted (T1w) image was corrected for intensity non-uniformity (INU) with N4BiasFieldCorrection (Tustison et al. 2010), distributed with ANTs 2.3.3 (Avants et al. 2008, RRID:SCR_004757), and used as T1w-reference throughout the workflow. The T1w-reference was then skull-stripped with a *Nipype* implementation of the antsBrainExtraction.sh workflow (from ANTs), using OASIS30ANTs as target template. Brain tissue segmentation of cerebrospinal fluid (CSF), white-matter (WM) and gray-matter (GM) was performed on the brain-extracted T1w using fast (FSL 6.0.5.1:57b01774, RRID:SCR_002823, Zhang, Brady, and Smith 2001). Brain surfaces were reconstructed using recon-all (FreeSurfer 7.2.0, RRID:SCR_001847, Dale, Fischl, and Sereno 1999), and the brain mask estimated previously was refined with a custom variation of the method to reconcile ANTs-derived and FreeSurfer-derived segmentations of the cortical gray-matter of Mindboggle (RRID:SCR_002438, Klein et al. 2017). Volume-based spatial normalization to one standard space (MNI152NLin2009cAsym) was performed through nonlinear registration with antsRegistration (ANTs 2.3.3), using brain-extracted versions of both T1w reference and the T1w template. The following template was selected for spatial normalization: *ICBM 152 Nonlinear Asymmetrical template version 2009c* [Fonov et al. (2009), RRID:SCR_008796; TemplateFlow ID: MNI152NLin2009cAsym].

**Functional data preprocessing**

For each of the 2 BOLD runs found per subject (across all tasks and sessions), the following preprocessing was performed. First, a reference volume and its skull-stripped version were generated using a custom methodology of *fMRIPrep*. Head-motion parameters with respect to the BOLD reference (transformation matrices, and six corresponding rotation and translation parameters) are estimated before any spatiotemporal filtering using mcflirt (FSL 6.0.5.1:57b01774, Jenkinson et al. 2002). BOLD runs were slice-time corrected to 0.978s (0.5 of slice acquisition range 0s-1.96s) using 3dTshift from AFNI (Cox and Hyde 1997, RRID:SCR_005927). The BOLD time-series (including slice-timing correction when applied) were resampled onto their original, native space by applying the transforms to correct for head-motion. These resampled BOLD time-series will be referred to as *preprocessed BOLD in original space*, or just *preprocessed BOLD*. The BOLD reference was then co-registered to the T1w reference using bbregister (FreeSurfer) which implements boundary-based registration (Greve and Fischl 2009). Co-registration was configured with six degrees of freedom. Several confounding time-series were calculated based on the *preprocessed BOLD*: framewise displacement (FD), DVARS and three region-wise global signals. FD was computed using two formulations following Power (absolute sum of relative motions, Power et al. (2014)) and Jenkinson (relative root mean square displacement between affines, Jenkinson et al. (2002)). FD and DVARS are calculated for each functional run, both using their implementations in *Nipype* (following the definitions by Power et al. 2014). The three global signals are extracted within the CSF, the WM, and the whole-brain masks. Additionally, a set of physiological regressors were extracted to allow for component-based noise correction (*CompCor*, Behzadi et al. 2007). Principal components are estimated after high-pass filtering the *preprocessed BOLD* time-series (using a discrete cosine filter with 128s cut-off) for the two *CompCor* variants: temporal (tCompCor) and anatomical (aCompCor). tCompCor components are then calculated from the top 2% variable voxels within the brain mask. For aCompCor, three probabilistic masks (CSF, WM and combined CSF+WM) are generated in anatomical space. The implementation differs from that of Behzadi et al. instead of eroding the masks by 2 pixels on BOLD space, a mask of pixels that likely to contain a volume fraction of GM is subtracted from the aCompCor masks. This mask is obtained by dilating a GM mask extracted from the FreeSurfer’s *aseg* segmentation, and it ensures components are not extracted from voxels containing a minimal fraction of GM. Finally, these masks are resampled into BOLD space and binarized by thresholding at 0.99 (as in the original implementation). Components are also calculated separately within the WM and CSF masks. For each CompCor decomposition, the *k* components with the largest singular values are retained, such that the retained components’ time series are sufficient to explain 50 percent of variance across the nuisance mask (CSF, WM, combined, or temporal). The remaining components are dropped from consideration. The head-motion estimates calculated in the correction step were also placed within the corresponding confounds file. The confound time series derived from head motion estimates and global signals were expanded with the inclusion of temporal derivatives and quadratic terms for each (Satterthwaite et al. 2013). Frames that exceeded a threshold of 0.5 mm FD or 1.5 standardized DVARS were annotated as motion outliers. Additional nuisance timeseries are calculated by means of principal components analysis of the signal found within a thin band (*crown*) of voxels around the edge of the brain, as proposed by (Patriat, Reynolds, and Birn 2017). The BOLD time-series were resampled into standard space, generating a *preprocessed BOLD run in MNI152NLin2009cAsym space*. First, a reference volume and its skull-stripped version were generated using a custom methodology of *fMRIPrep*. All resamplings can be performed with *a single interpolation step* by composing all the pertinent transformations (i.e. head-motion transform matrices, susceptibility distortion correction when available, and co-registrations to anatomical and output spaces). Gridded (volumetric) resamplings were performed using antsApplyTransforms (ANTs), configured with Lanczos interpolation to minimize the smoothing effects of other kernels (Lanczos 1964). Non-gridded (surface) resamplings were performed using mri_vol2surf (FreeSurfer).

Many internal operations of *fMRIPrep* use *Nilearn* 0.9.1 (Abraham et al. 2014, RRID:SCR_001362), mostly within the functional processing workflow. For more details of the pipeline, see [the section corresponding to workflows in *fMRIPrep*’s documentation](https://fmriprep.readthedocs.io/en/latest/workflows.html).

**Supplementary Text**

**Luminance analysis**

Mean framewise luminance was computed for each movie clip by converting each video frame to grayscale using the ITU-R BT.709 luma formula (Y' = 0.2126R + 0.7152G + 0.0722B), applied to gamma-encoded 8-bit RGB channels (scale 0–255). The coefficients reflect the photopic luminous efficiency/ sensitivity across the visible spectrum for high-definition content. For each movie, luminance was first computed at the frame level by averaging luma values across all pixels within each frame, yielding one luminance value per frame. These frame-level values were then averaged across all frames falling within each fMRI acquisition window (TR = 2 s), yielding one luminance value per acquired volume (247 volumes for the negative clip; 240 volumes for the neutral clip). This down sampling aligned the luminance time series with the BOLD time series for subsequent association testing.”

The two movies differed in overall luminance. The negative movie (Curve) had considerably lower luminance (M = 50.9, SD = 18.8) than the neutral movie (Pottery; M = 114.9, SD = 23.3; Cohen's d = 3.05). This difference is readily explained by the visual content of each film. Curve depicts a woman stranded on the edge of a cliff in a sustained low-light environment dominated by dark overcast skies, shadowed rock faces, and muted tones, with scenes involving dark blood against already dim surroundings and the character dressed in dark clothing throughout, contributing to a consistently dull and visually restrained palette. In contrast, Pottery follows artists crafting pottery in a well-lit studio setting, with bright white clay, warm natural lighting, and an array of colorful objects and paints visible throughout, producing an overall visual impression of warmth and high luminance across the majority of frames.

**TABS-based Findings Summary**

The Trondheim Aging Brain Study (TABS) dataset has supported several recent publications from our group, each addressing distinct questions. Below, we outline how the present study differs from and extends these prior works in terms of both biological insight and methodological approach.

Dave et al. (2025, Journal of Neuroscience) examined LC task-evoked activity and task-dependent generalized psychophysiological interactions (gPPI) during a facial emotion paradigm that manipulated emotional ambiguity. That study used a GLM-based approach to quantify LC BOLD responses to static morphed face stimuli (ranging from unambiguous to fully ambiguous happy–fearful blends) and tested how LC–prefrontal cortex (PFC) connectivity changes during cognitively demanding absolute ambiguous faces. The central finding was that older adults show increased LC activation and strengthened LC–dorsolateral PFC coupling specifically during ambiguity processing, and that this heightened engagement was linked to better emotional wellbeing. Critically, this work focused on LC activation during a discrete cognitive-emotional challenge using static face stimuli, treating the LC as a task-responsive node and examining its coupling with specific prefrontal targets. However, the use of static faces inherently limits ecological validity, as real-world emotional processing unfolds dynamically and involves sustained, context-dependent engagement of neuromodulatory systems that cannot be captured by brief, trial-locked evoked responses. Importantly, the study also established high-resolution, subject-specific LC masks derived from 7T magnetization transfer imaging, which provide the anatomical foundation for the LC seed definitions used in the present work.

Ye et al. (2025, Neurobiology of Aging) took a purely cortical perspective, examining functional dedifferentiation of the frontoparietal network (FPN) during naturalistic movie-viewing using gradient mapping techniques. That study quantified FPN dispersion in a multidimensional gradient space and linked increased cortical dedifferentiation in older adults to poorer mental wellbeing (anxiety and depression), with emotion regulation serving as a mediating mechanism. Importantly, that work did not examine the LC at all, nor did it investigate which neuromodulatory mechanisms might contribute to the observed cortical dedifferentiation patterns.

The present manuscript bridges and extends both prior efforts in several keyways.

First, rather than examining LC task-evoked responses to static faces (as in Dave et al.) or cortical gradients in isolation from any neuromodulatory source (as in Ye et al.), we use gradient-based mapping to characterize the spatial topography of LC-cortical functional connectivity gradients during naturalistic movie-viewing. By employing dynamic emotional movie stimuli rather than static faces, we capture how the LC–cortex system is organized under ecologically valid, sustained emotional processing conditions, moving beyond trial-locked seed-to-target analyses. This work also directly benefits from the subject-specific, high-resolution 7T LC masks established in Dave et al. (2025, J. Neuroscience) ensuring anatomically precise LC seed definitions that are critical for reliable connectivity estimation from a structure of this size.

Second, we demonstrate that the primary LC-cortical gradient is anchored by catecholaminergic chemoarchitectural receptor distributions, providing a neurochemical basis for LC–cortex functional organization that neither prior study addressed.

Third, we show that aging selectively increases FPN dispersion during negative emotional processing, offering a complementary perspective to the cortical dedifferentiation reported by Ye et al. by situating it within the context of LC-cortical connectivity. While the Ye et al. findings characterized dedifferentiation at the cortical level, the present work extends this by examining how the LC-noradrenaline system may contribute to such patterns, suggesting that age-related changes in neuromodulatory input represent one potential pathway through which cortical dedifferentiation emerges during emotional processing. Finally, we demonstrate that this LC-linked FPN dispersion is associated with emotional wellbeing derived from the principal component analysis based approach as executed in Dave et al., connecting neuromodulatory organization to affective outcomes.

In summary, the three papers address complementary but distinct questions. Dave et al. established that the LC is functionally engaged during emotionally demanding tasks in aging and provided the high-resolution LC masks that enabled the present analyses; Ye et al. characterized cortical network dedifferentiation during emotional processing; and the present work examines how LC-cortical connectivity gradients are neurochemically organized, context-dependently altered in aging under naturalistic conditions, and associated with affective outcomes, offering a neuromodulatory perspective on cortical dedifferentiation that complements the prior findings.

**Supplementary Figures**

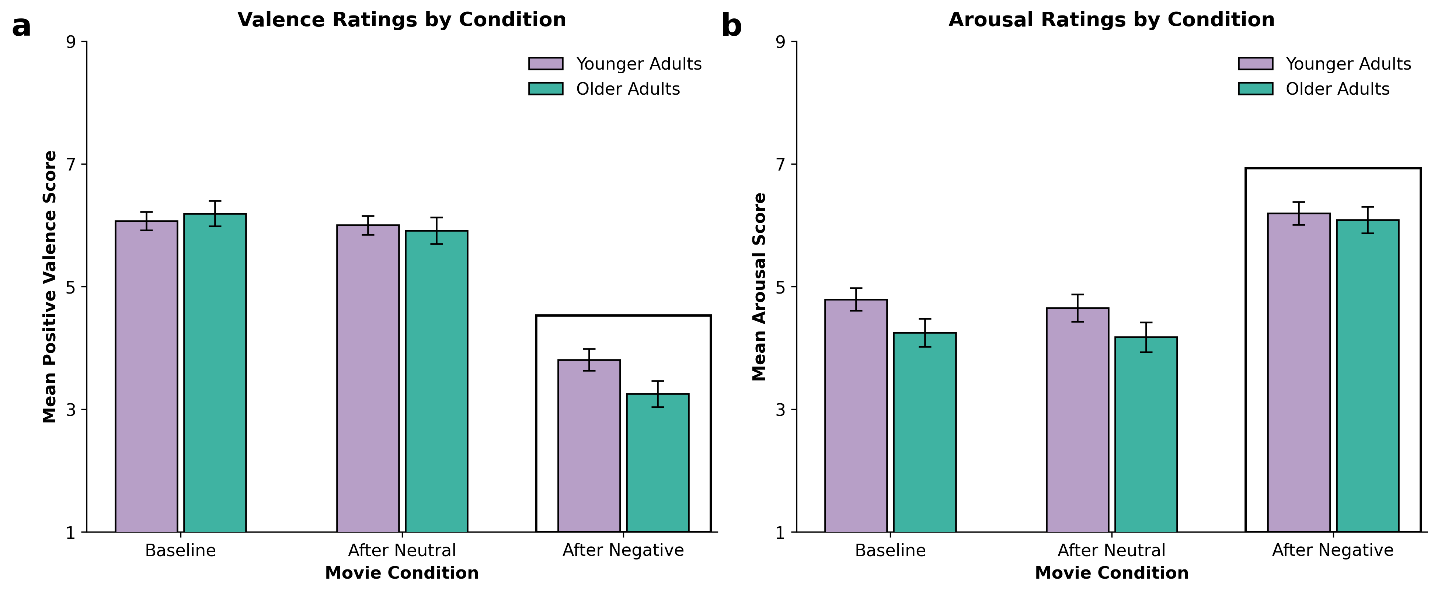
**Supplementary Figure 1**

**Self-reported valence and arousal across movie conditions in younger and older adults.** Panels depict (a) valence and (b) arousal ratings assessed using the 9-point Self-Assessment Manikin at baseline, after neutral movie viewing, and after negative movie viewing. A 3 (Condition: baseline, neutral, negative) × 2 (Age: young, older) mixed ANOVA showed a robust main effect of condition for valence, F(2, 276) = 159.75, p < 0.001, η²ₚ = .54, and arousal, F(2, 276) = 80.04, p < 0.001, η²ₚ = .37. Post-hoc paired t-tests indicated that positive valence decreased markedly after negative movie viewing relative to baseline (t = 15.05, p < 0.001, d = 1.27) and neutral movie viewing (t = 13.13, p < 0.001, d = 1.11), with no difference between baseline and neutral conditions (ps > 0.10). For arousal, negative movie viewing elicited significantly higher ratings compared with baseline (t = 10.12, p < 0.001, d = 0.86) and neutral movie viewing (t = 9.75, p < 0.001, d = 0.82), again with no difference between baseline and neutral conditions (ps > 0.10). No Condition × Age interactions were observed for valence, F(2, 276) = 2.27, p = .11, η²ₚ = 0.016, or arousal, F(2, 276) = 1.18, p = .31, η²ₚ = 0.008, indicating comparable emotional modulation across age groups. Across participants, arousal was negatively correlated with positive valence (r = −0.31, p < 0.001), confirming that heightened arousal was systematically coupled with reduced positive affect following negative movie exposure.

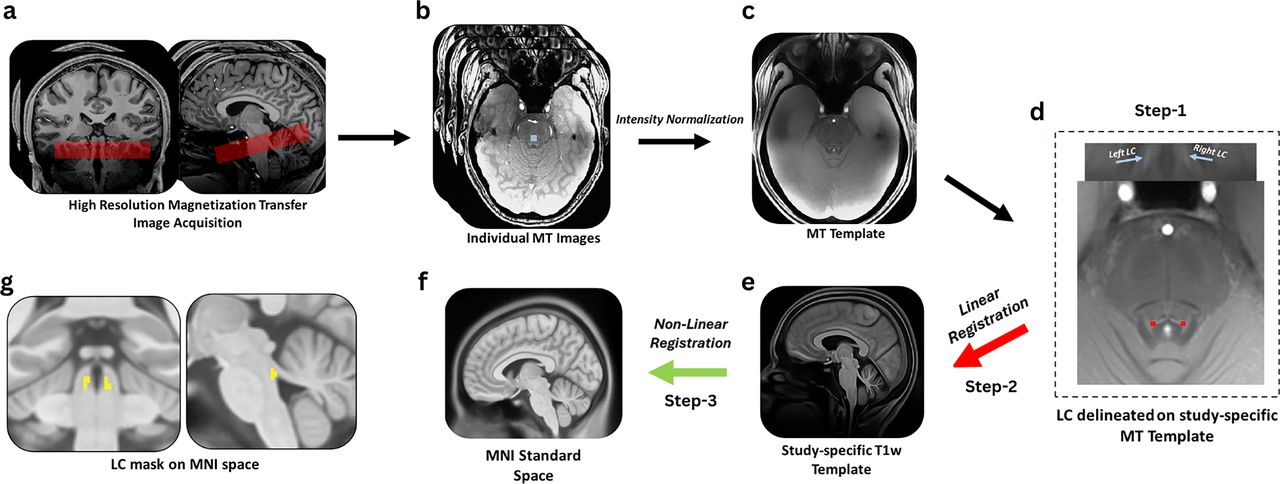
**Supplementary Figure 2**

**Locus coeruleus delineation pipeline.** MT-TFL preprocessing and spatial transformation pipeline. a, High-resolution MT-TFL images acquired by positioning the approximately perpendicular to the pons, encompassing the region between the inferior colliculus and the inferior border of the pons. Panel **b** shows an example of individual MT-TFL images acquired using this sequence with 10 × 10 voxel (blue) placed on pons. Panel **c** demonstrates a study-specific MT-TFL template created after intensity normalizing the individual MT-TFL images. Panel **d** shows left, and right LC delineated on subject-specific MT-TFL template using ITK-SNAP. Panel **e** depicts study-specific T1w template created to act as a bridge to transform LC mask to MNI space; the red arrow indicates MT template and was rigidly registered to the whole-brain study-specific T1w template. Panel **f** shows MNI space where T1w template was nonlinearly registered (green arrow). Panel **g** depicts left and right LC mask transformed to MNI space after using the transformation matrices obtained from Step 2 and Step 3 which was then used to extract the timeseries. Figure used from Dave et al. (2025).

**Supplementary Figure 3**

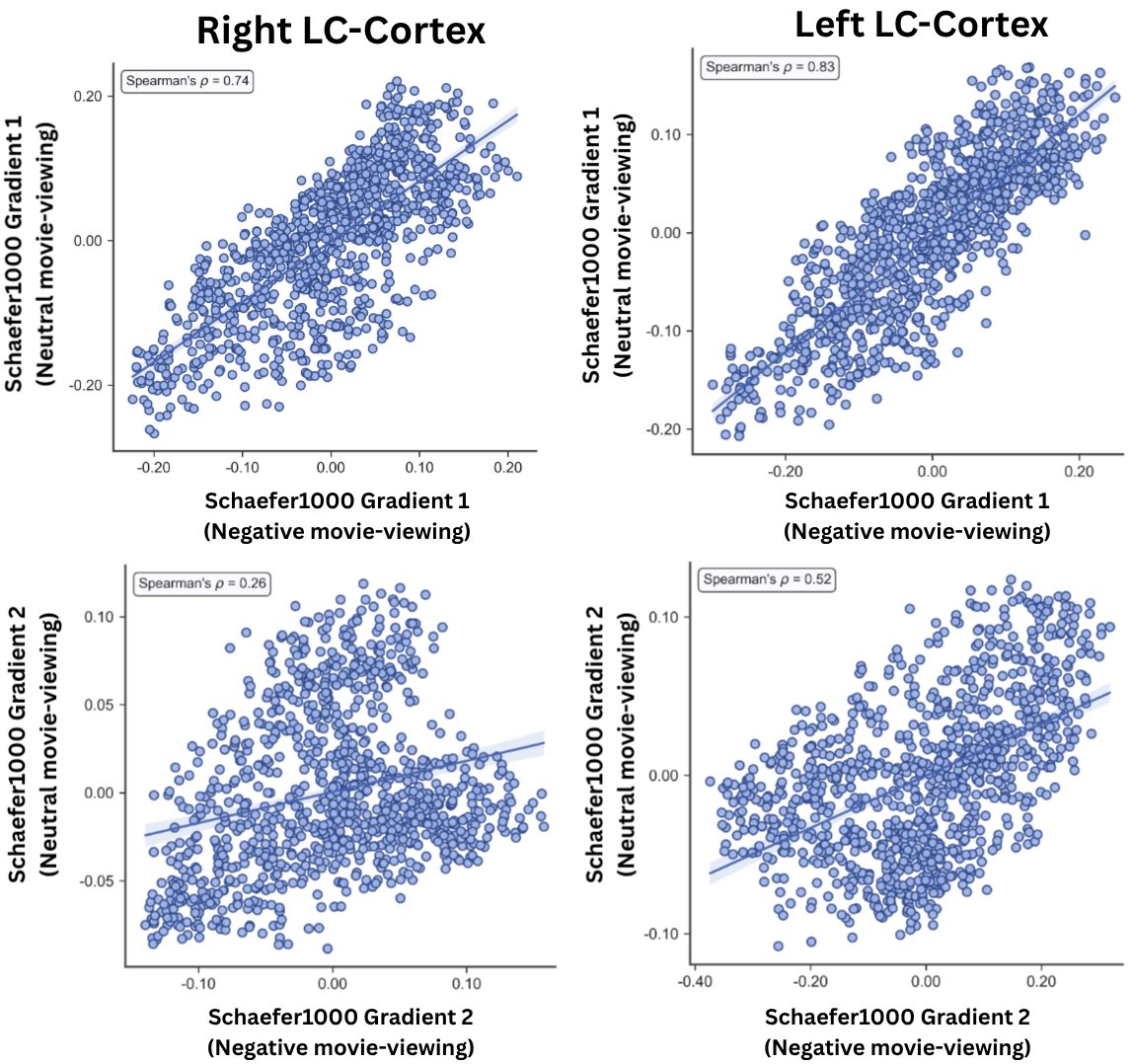

**Cross-condition spatial correspondence of the primary (G1) and secondary (G2) LC–linked cortical gradient between neutral and negative movie-viewing**. Scatter plots show parcel-wise gradient values (Schaefer-1000 atlas) for G1 in the neutral condition (y-axis) plotted against G1 in the negative condition (x-axis) for G1 (top row) and G2 (bottom row), separately for the right LC (left panel) and left LC (right panel). Each point represents one of 1000 cortical parcels. Spearman rank correlations indicate high cross-condition agreement of the primary gradient topography in both hemispheres, confirming that G1 reflects a stable organizational axis preserved across emotional valence. Agreement was considerably lower for G2, indicating that the secondary gradient is substantially reorganized between conditions, most markedly for the right LC.

**Supplementary Figure 4**

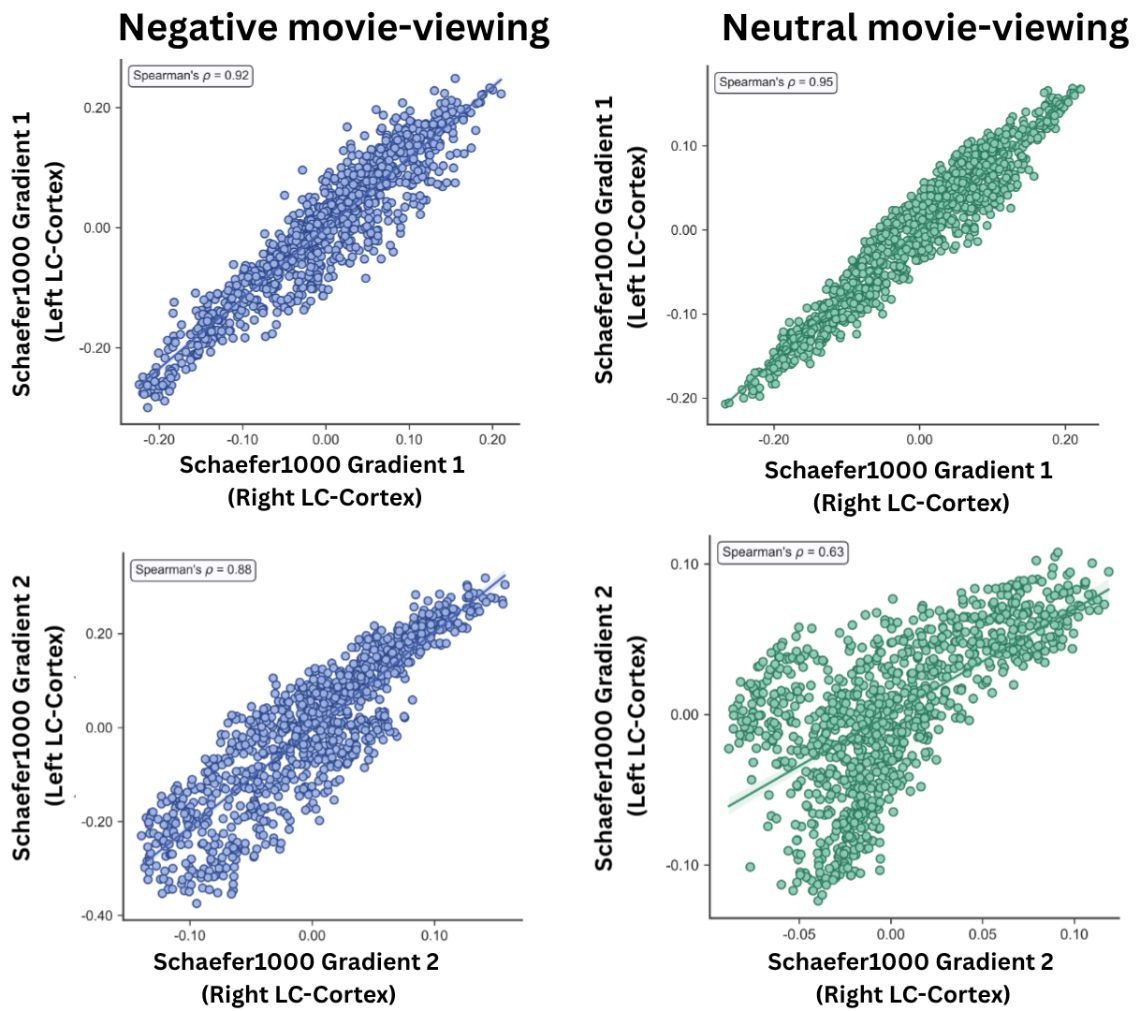

**Cross-hemisphere spatial correspondence of the primary (G1) and secondary (G2) LC–linked cortical gradient.** Scatter plots show parcel-wise gradient values (Schaefer-1000 atlas) for G1 derived from the left LC (y-axis) plotted against G1 derived from the right LC (x-axis), for G1 (top row) and G2 (bottom row), separately for the negative (left panel) and neutral (right panel) movie-viewing conditions. Each point represents one of 1000 cortical parcels. High Spearman rank correlations indicate strong cross-hemispheric agreement of the primary gradient topography in both movie-viewing conditions, confirming that G1 reflects a hemispherical consistent organizational axis across emotional contexts. Cross-hemispheric agreement for G2 was similarly high during negative movie-viewing but lower during neutral movie-viewing, indicating that the secondary gradient topography diverges between hemispheres specifically in the neutral condition.

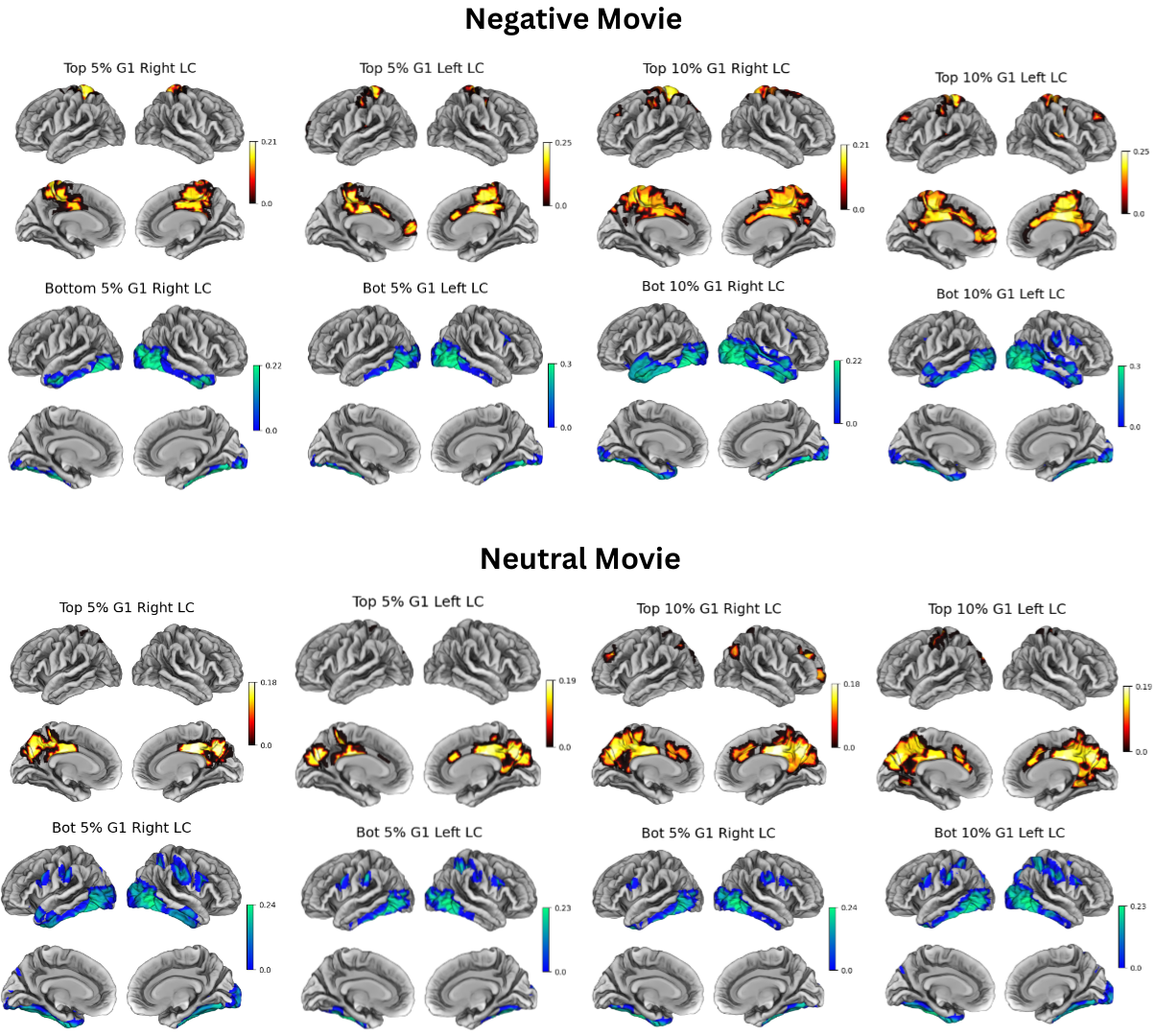
**Supplementary Figure 5**

**Cortical parcels most strongly and weakly expressed along Gradient 1 for left and right LC.** Functional Maps display the top 5% and top 10% cortical parcels with the highest absolute loadings (strongly connected regions) and the lowest absolute loadings (weakly connected regions) on Gradient 1, derived separately for left and right LC connectivity. These parcels indicate the cortical territories that contribute most—and least—to the principal axis of variation in LC–cortical coupling. Color intensity reflects the magnitude of Gradient 1 loadings.

**Supplementary Figure 6**

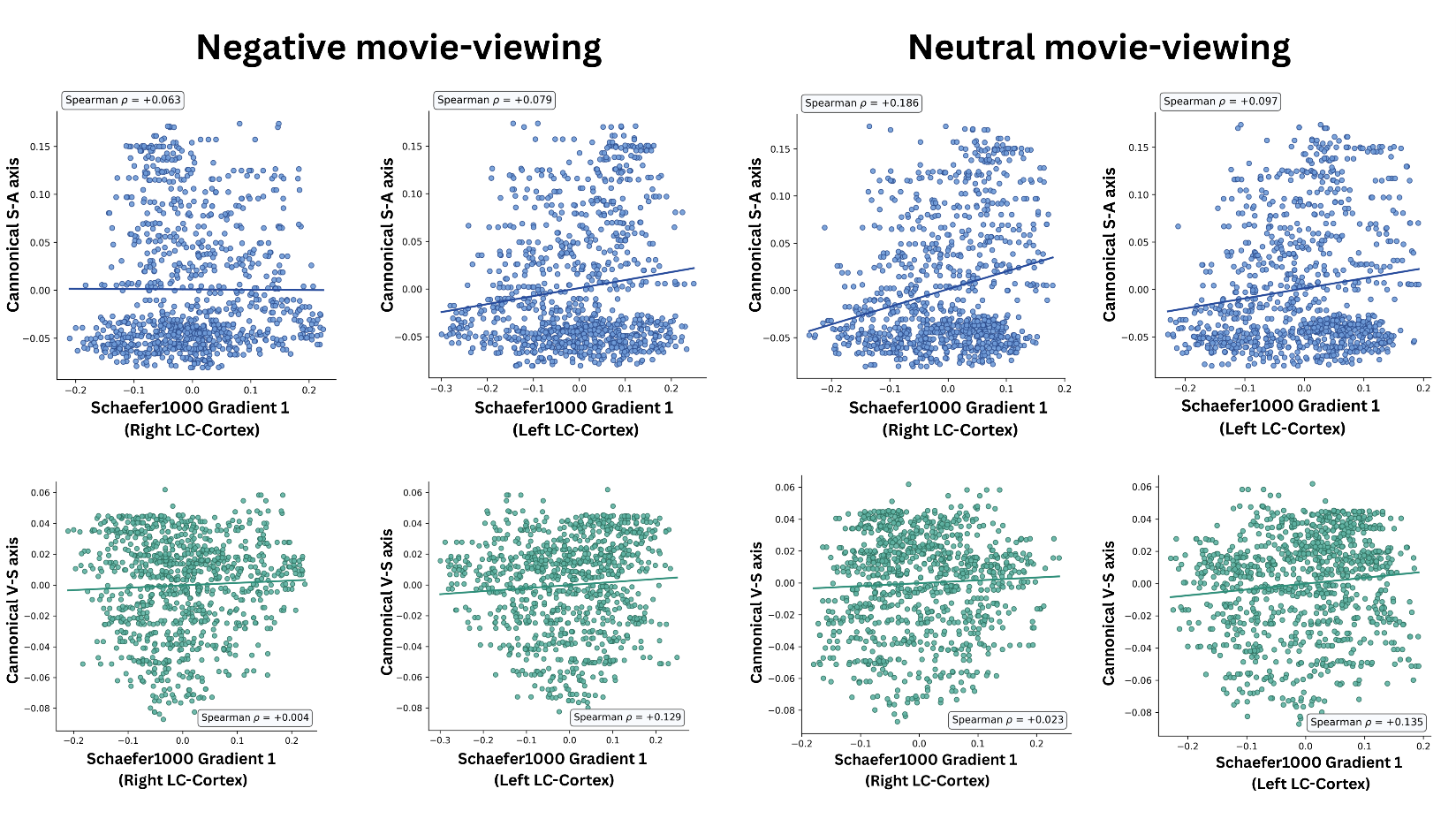

**Spatial alignment of the LC–linked primary cortical gradient (G1) with canonical macroscale cortical axes.** Scatter plots show parcel-wise gradient values for G1 derived from the right LC and the left LC–linked primary cortical gradient plotted against the canonical sensorimotor–association (S–A) axis (top row, blue) and visual–somatomotor (V–S) axis (bottom row, green), separately for negative (left panels) and neutral (right panels) movie-viewing conditions.

**Supplementary Figure 7**

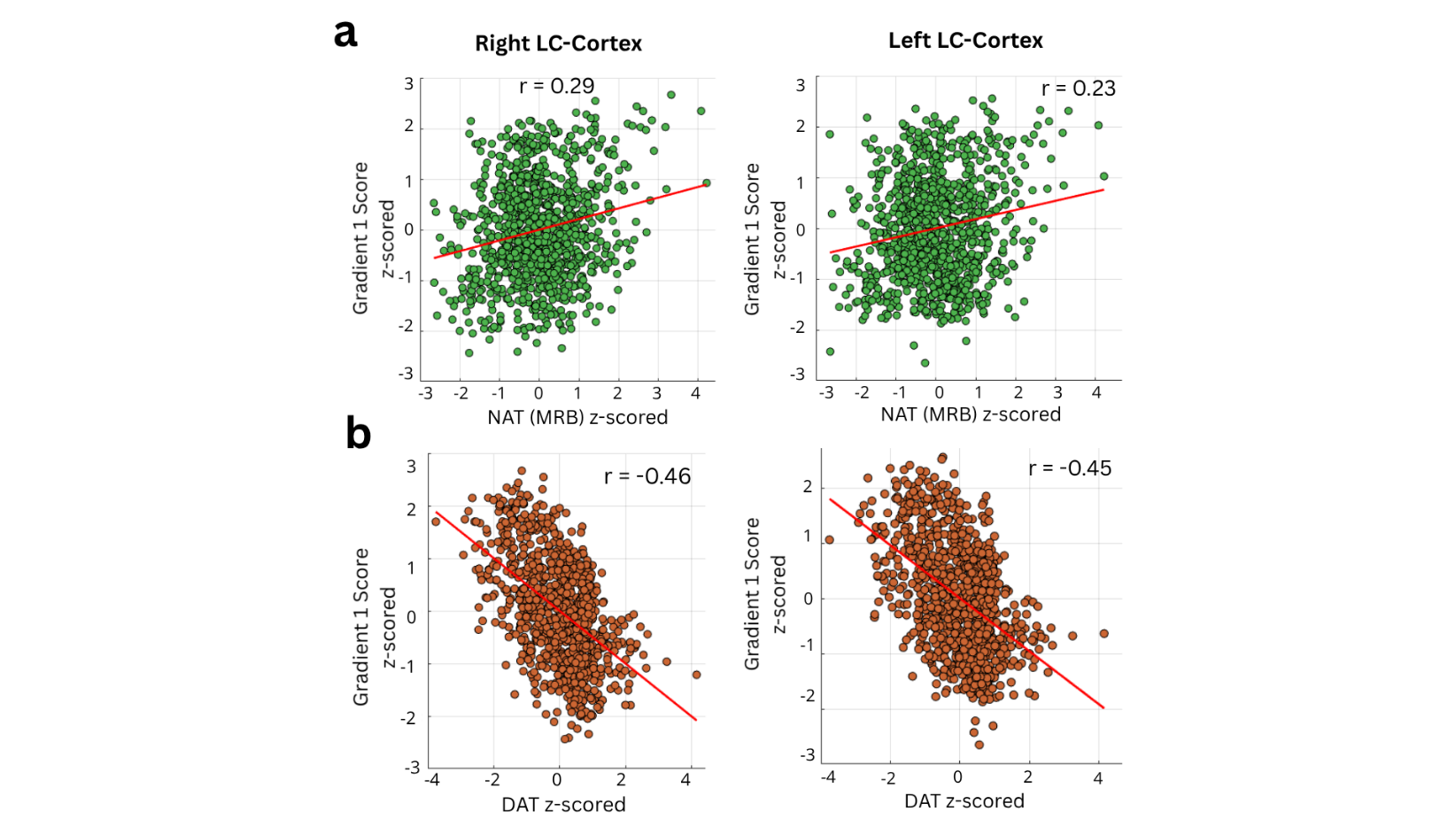

**Correlation of Noradrenaline and Dopamine Transporter with G1.** Right LC–cortical Gradient 1 (z) scores showed a positive correlation with regional noradrenaline transporter (NAT) density (Right LC: r = 0.29, p < 0.001; Left LC: r = 0.22, p < 0.001; Supplementary Fig. 7a). In contrast, dopamine transporter (DAT) density correlated negatively with Gradient 1 (Right LC: r = –0.46, p < 0.001; Left LC: r = –0.45, p < 0.001)

**Supplementary Figure 8**

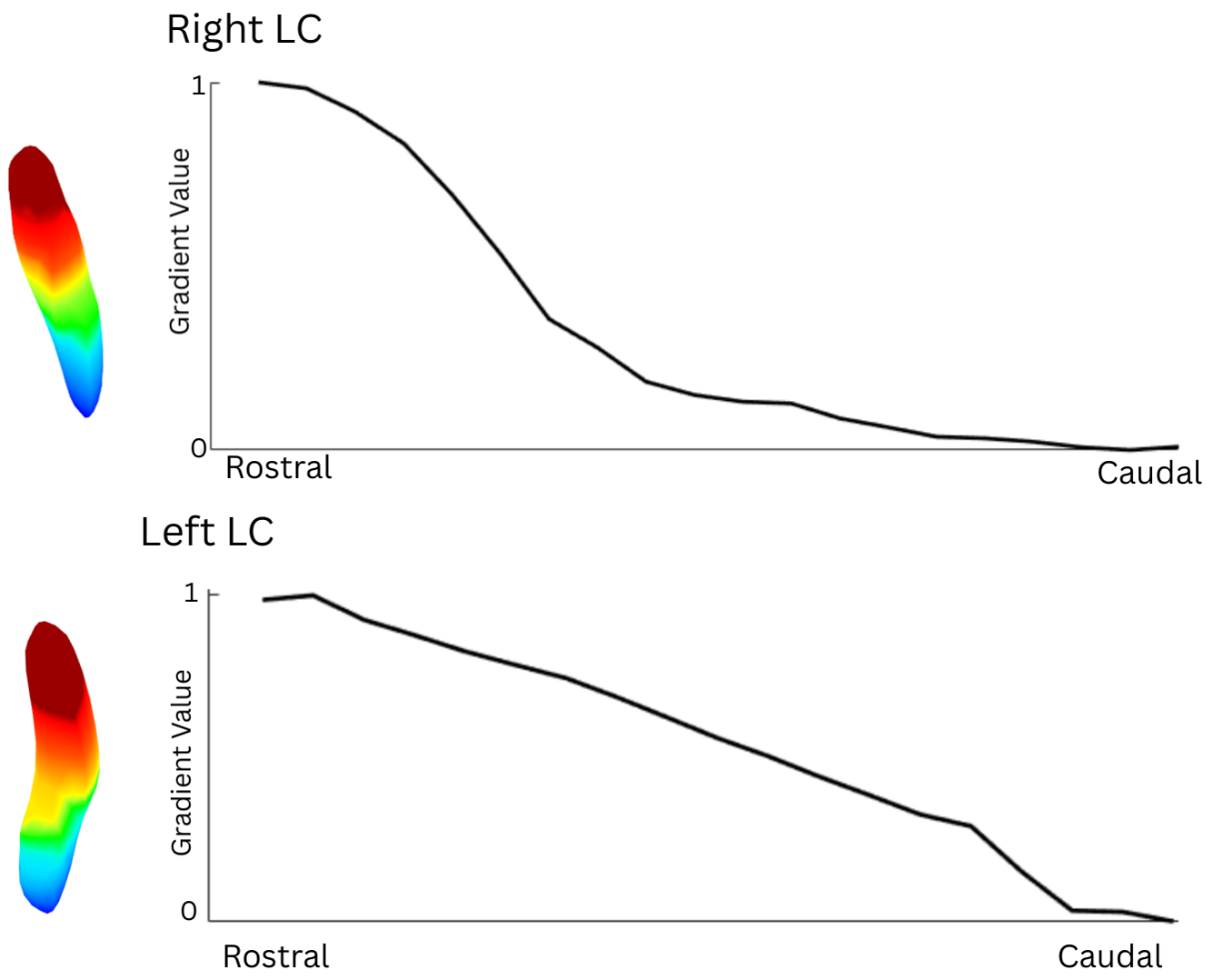

**Topographic gradient of the locus coeruleus based on cortical connectivity.** The first LC gradient, derived from connectopic mapping of LC–cortical connectivity profiles, explained 75.7% (right LC) and 71.0% (left LC) of the variance. Similar colors indicate LC voxels with similar cortical connectivity profiles. Decay plots depict changes in LC–cortical connectivity along the rostro–caudal Z axis.

**Supplementary Figure 9**
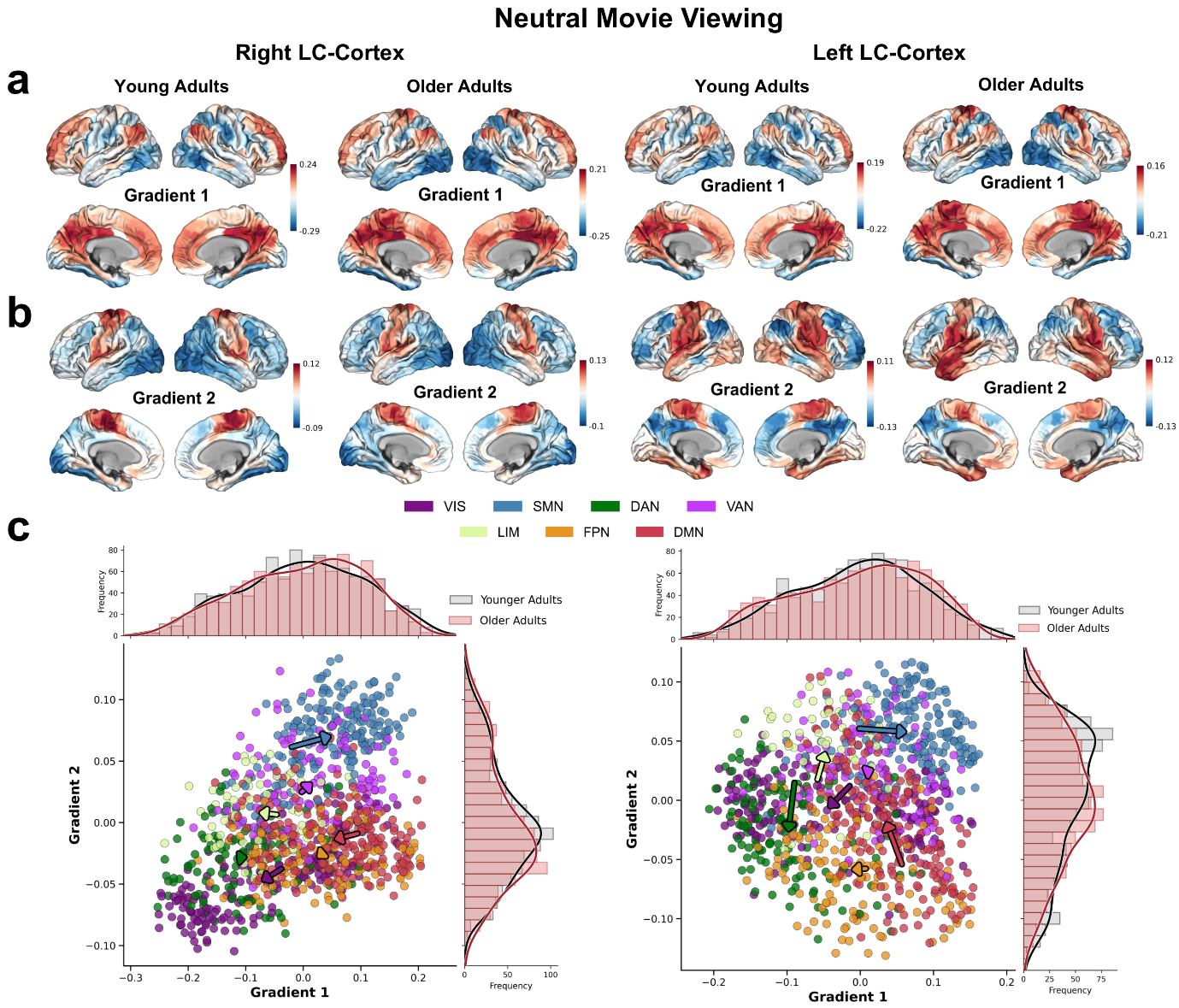

***Age-related network shifts within the LC–linked cortical networks 2D gradient manifold during neutral movie-viewing*.** **(a, b)** Group-averaged cortical maps of Gradient 1 (a) and Gradient 2 (b) for younger and older adults, derived separately from the right LC (left columns) and left LC (right columns) during neutral movie-viewing condition; Color reflects gradient loadings. **(c)** Scatter plot shows older adults G1 × G2 embedding of LC–linked cortical gradient for the right LC (left panel) and left LC (right panel), with parcels colored by their Yeo seven-network assignment (VIS, SMN, DAN, VAN, LIM, FPN, DMN). Marginal histograms above and to the right of each scatter plot show the distribution of younger (grey) and older (red) adult gradient scores along G1 (top) and G2 (right), respectively. Arrows depict age-related centroid displacement: each arrow originates at the younger-adult network centroid and terminates at the older-adult centroid, with arrow length indicating Euclidean displacement and direction indicating the relative ΔG1 and ΔG2 contributions. Network abbreviations: VIS, visual; SMN, somatomotor; DAN, dorsal attention; VAN, ventral attention; LIM, limbic; FPN, frontoparietal; DMN, default mode.

**Supplementary Figure 10**

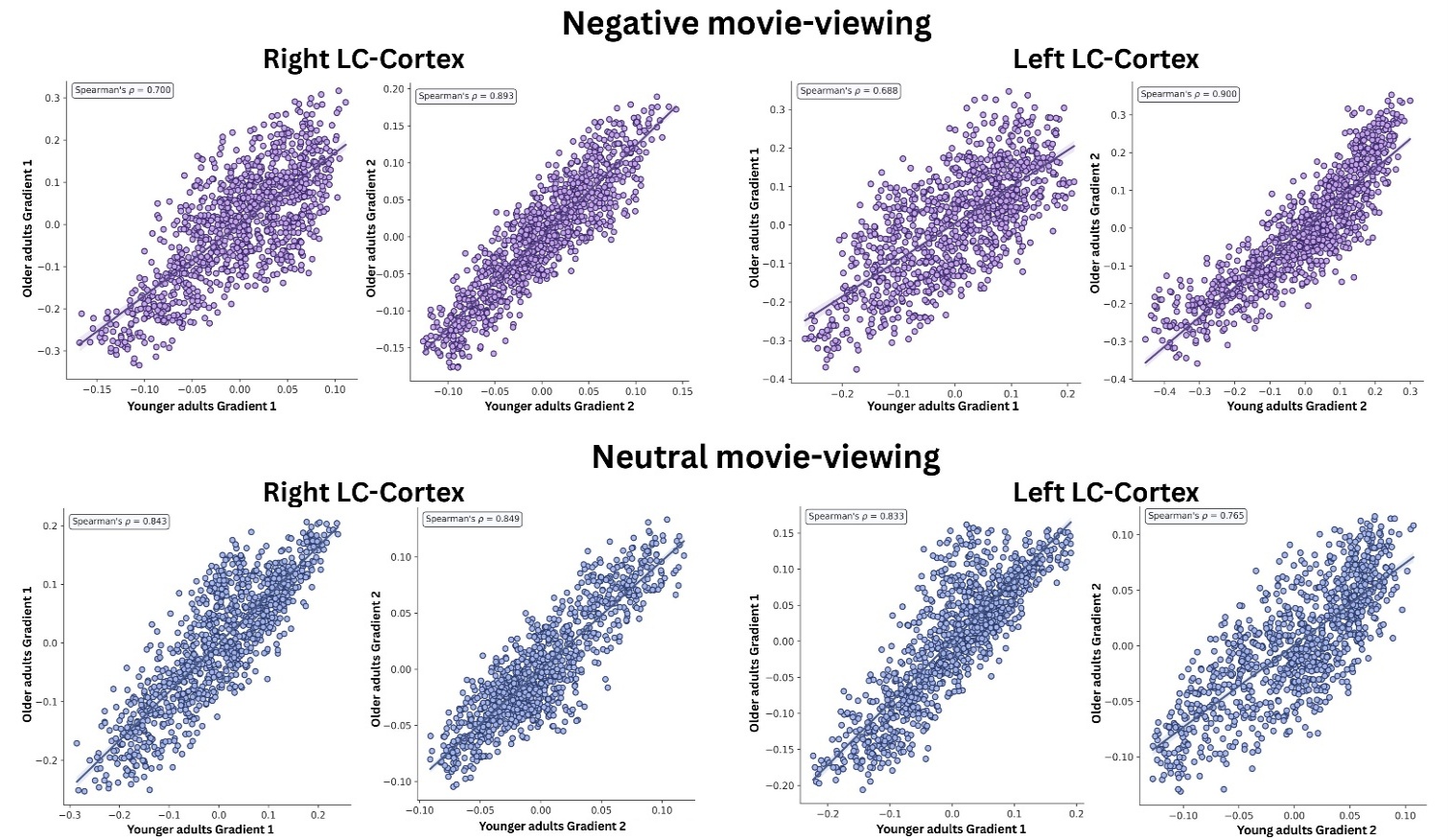

**Age-group correlations of LC–cortex gradient topographies.** Scatter plots showing parcel-wise spatial correlations between younger and older adults' LC–cortex gradient maps, separately for each movie-viewing condition, LC hemisphere, and gradient. Top row: negative movie-viewing condition. Bottom row: neutral movie-viewing condition. Within each row, panels display from left to right: right LC Gradient 1 (G1) and Gradient 2 (G2), and left LC G1 and G2. Each point represents one cortical parcel; the x-axis shows the group-averaged gradient score in younger adults, and the y-axis shows the corresponding score in older adults.

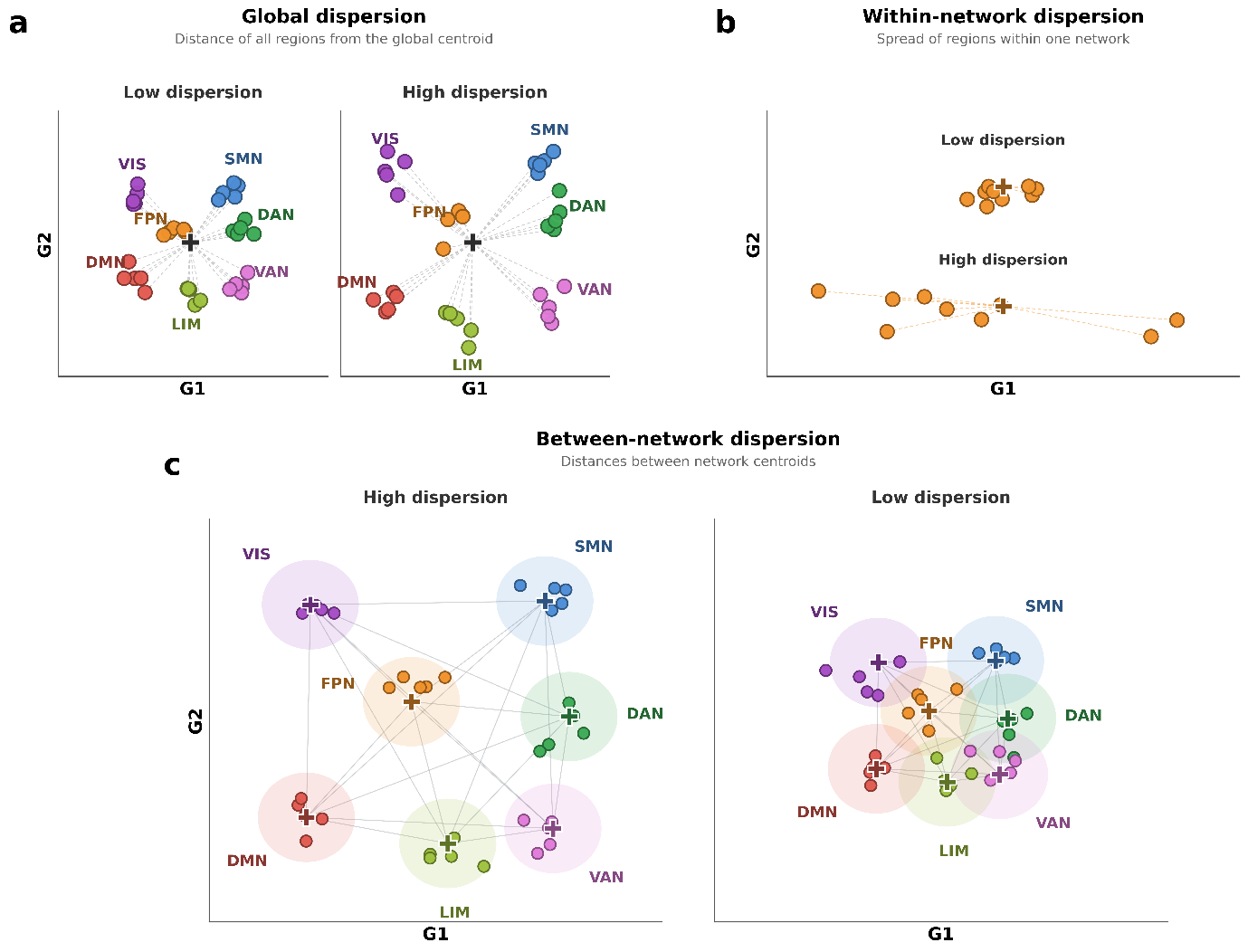
**Supplementary Figure 11**

***Schematic illustration of the three dispersion measures in two-dimensional gradient space.*** *Each panel illustrates how a different dispersion measure is computed in two-dimensional G1 × G2 gradient space, contrasting low- and high-dispersion configurations. Each dot represents a cortical parcel colored by Yeo seven-network assignment, centroids are marked by black crosses, and dashed lines indicate the Euclidean distances being summarized.* ***(a)*** *Global dispersion: mean Euclidean distance of all cortical parcels from the overall cortical centroid. Low dispersion (left) shows parcels clustered near the centroid; high dispersion (right) shows parcels spread further from it.* ***(b)*** *Within-network dispersion: mean Euclidean distance of parcels belonging to a single network from that network's centroid (FPN shown here). Low dispersion (top) shows tight clustering around the FPN centroid; high dispersion (bottom) shows greater spread.* ***(c) Between-network dispersion:*** *mean pairwise Euclidean distance between Yeo network centroids. High dispersion (left) shows widely separated centroids; low dispersion (right) shows centroids clustered more tightly.*

**Supplementary Figure 12**

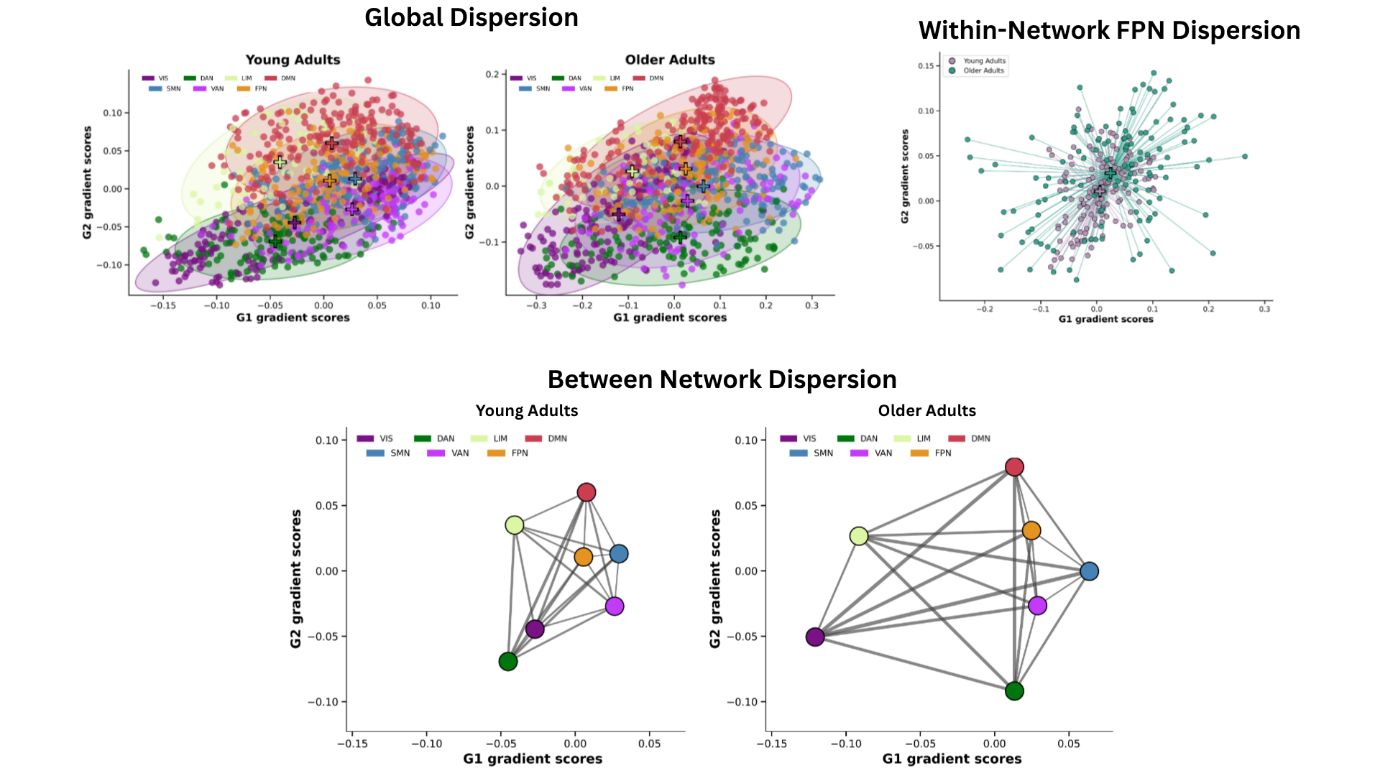

***Empirical visualization of dispersion measures in the right LC–linked cortical gradient embedding for younger and older adults****. Subject-averaged parcel coordinates in two-dimensional G1 × G2 gradient space, with parcels colored by Yeo seven-network assignment. Top left: Global dispersion in younger adults (left sub-panel) and older adults (right sub-panel). Top right: Within-network FPN dispersion showing frontoparietal network parcels from both age groups overlaid in a single plot, with lines connecting each parcel to its age-group FPN centroid; tighter clustering in younger adults (purple) contrasts with the more dispersed distribution in older adults (teal). Bottom: Between-network dispersion in younger adults (left) and older adults (right); each dot represents one network centroid in G1 × G2 space, with lines indicating pairwise Euclidean distances between centroids. Network abbreviations: VIS, visual; SMN, somatomotor; DAN, dorsal attention; VAN, ventral attention; LIM, limbic; FPN, frontoparietal; DMN, default mode.*

**Supplementary Figure 13**

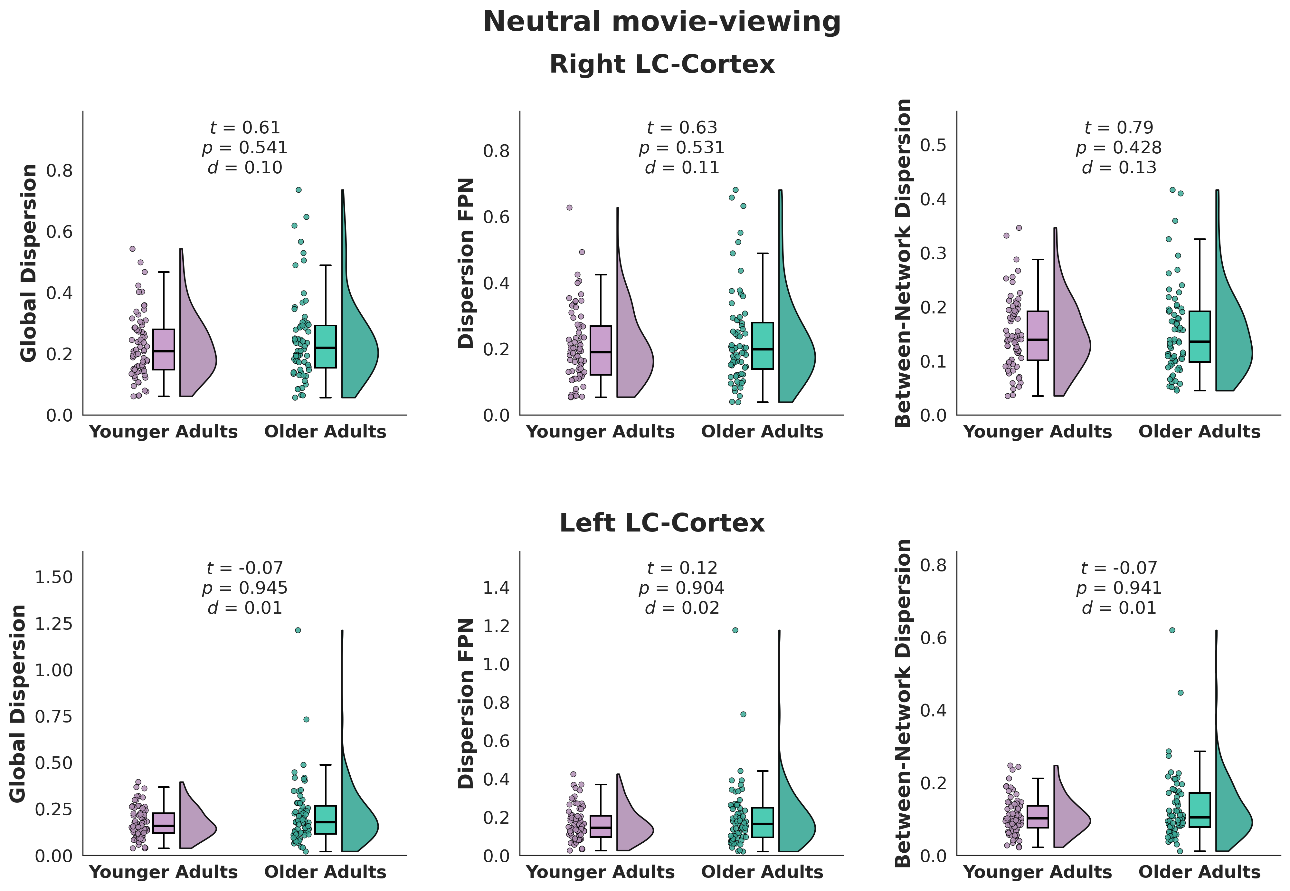

***Age-related differences in LC–linked cortical dispersion during neutral movie-viewing.*** *Raincloud plots show subject-level global dispersion (left column), within-network FPN dispersion (middle column), and between-network dispersion (right column) for younger adults (purple) and older adults (teal), separately for the right LC–cortex (top row) and left LC–cortex (bottom row) embeddings.*

**Supplementary Figure 14**

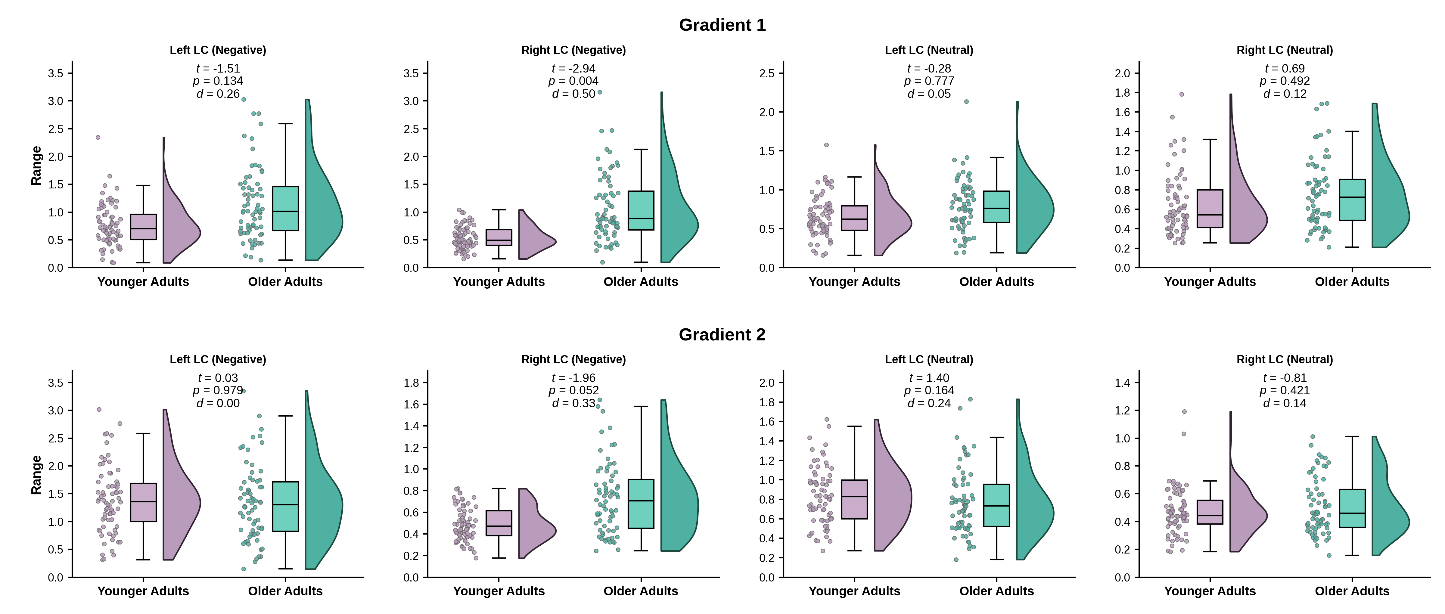

**Age-related differences in per-axis gradient range for the right and left LC, by movie-viewing condition.** Raincloud plots showing the parcel-wise range of gradient scores along Gradient 1 (top row) and Gradient 2 (bottom row) in younger adults (purple) and older adults (teal). From left to right within each row: left LC negative, right LC negative, left LC neutral, right LC neutral. Each dot represents one participant; box plots display the median and interquartile range, and density curves show the distribution shape. Inset values report independent-samples t-statistics, p-values, and Cohen's d for the between-group comparison.

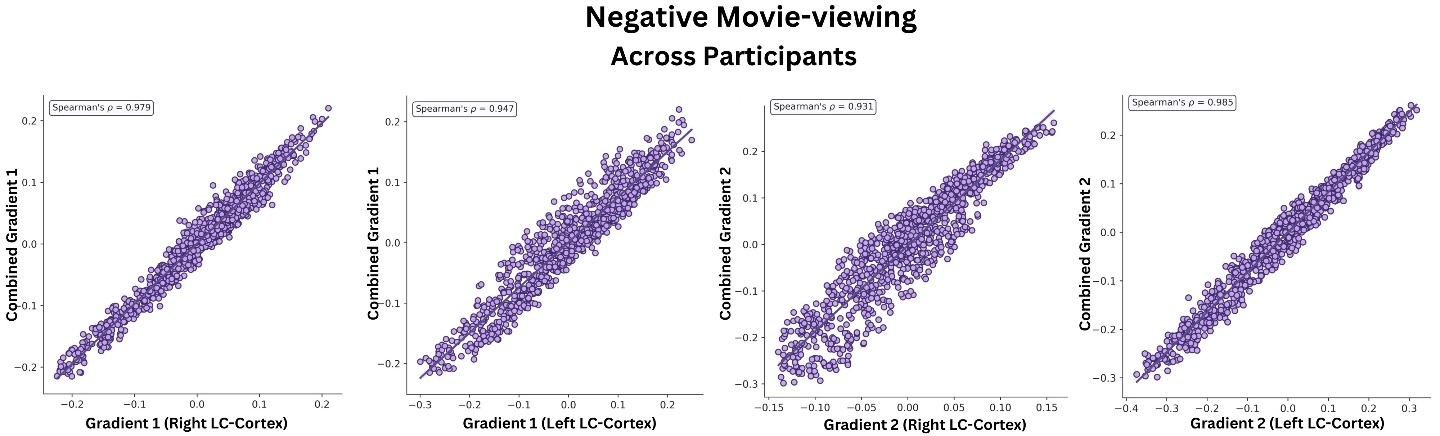
**Supplementary Figure 15**

***Bilateral consistency of the primary and secondary LC–linked cortical gradients across participants.*** *Scatter plots showing parcel-wise spatial correlations between hemisphere-specific and combined bilateral LC–linked cortical gradients in the negative movie-viewing condition. Each point represents one cortical parcel; the x-axis shows the gradient scores derived from hemisphere-specific seed and y-axis gradient scores derived from the combined bilateral LC seed.*

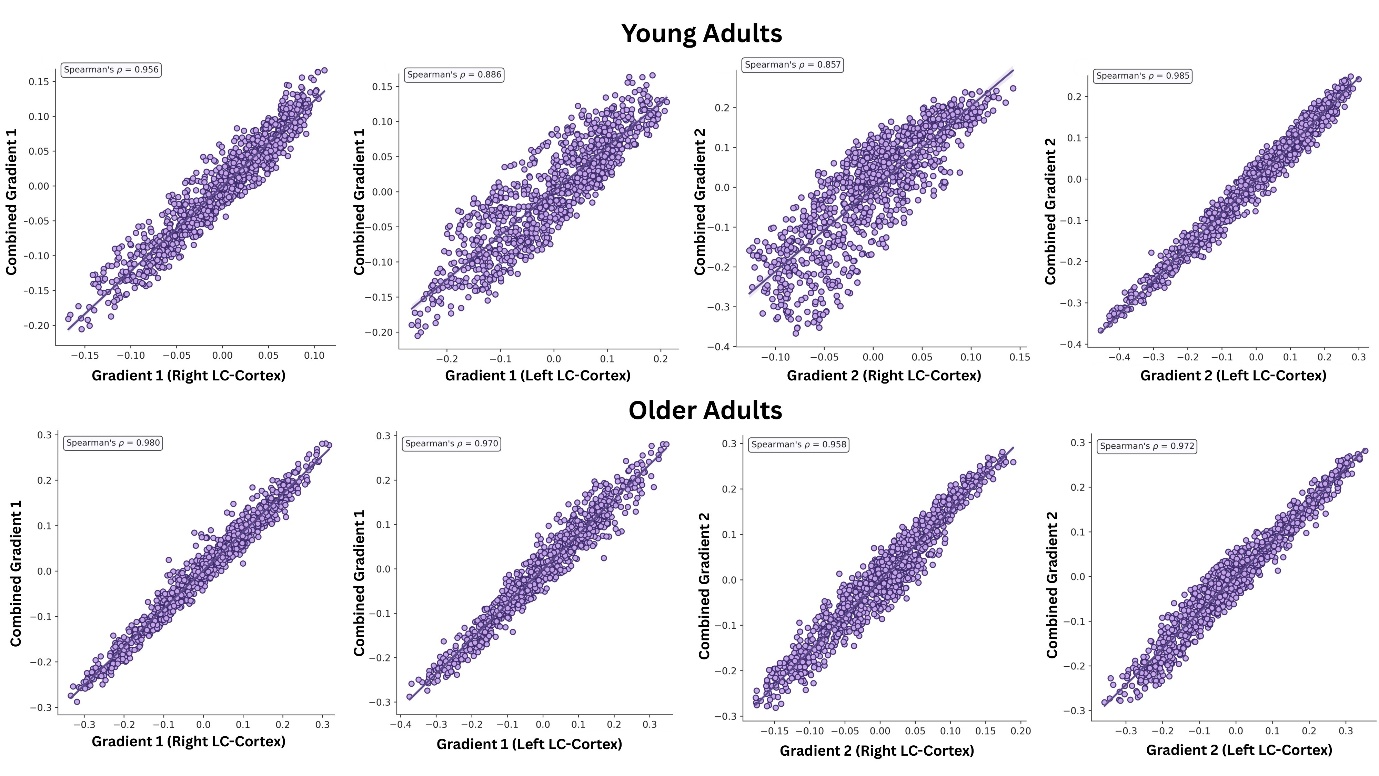
**Supplementary Figure 16**

**Bilateral consistency of the primary and secondary LC–linked cortical gradients in young and older adults.** Scatter plots showing parcel-wise spatial correlations between hemisphere-specific and combined bilateral LC–linked cortical gradients in the negative movie-viewing condition, separately for younger adults (top row) and older adults (bottom row).

**Supplementary Figure 17**

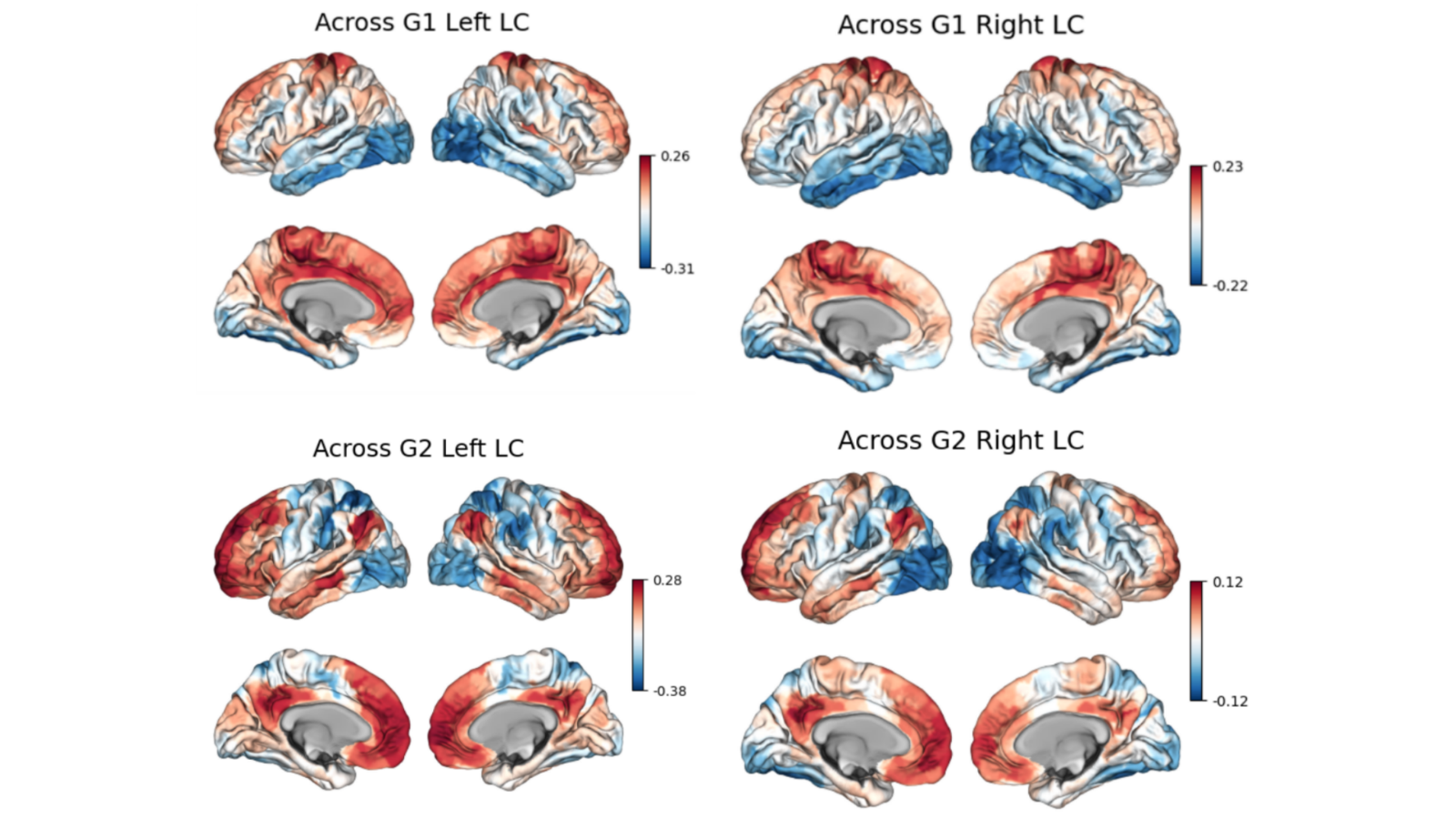

**LC–cortex connectivity patterns in 980 cortical parcels.** Surface maps displaying locus coeruleus (LC)–cortical connectivity patterns derived from an independent 980-region cortical parcellation. To confirm that our findings were not driven by parcellation choice, we replicated the primary analyses in negative movie condition and compared the resulting gradient maps with those obtained from the original parcellation. Voxelwise correlations between the two parcellation-derived maps demonstrated near-identical spatial topographies for G1 (right LC–cortex: r = 0.97; left LC–cortex: r = 0.98), and and G2 (right LC–cortex: r = 0.96; left LC–cortex: r = 0.95) confirming the robustness and reproducibility of the observed patterns.

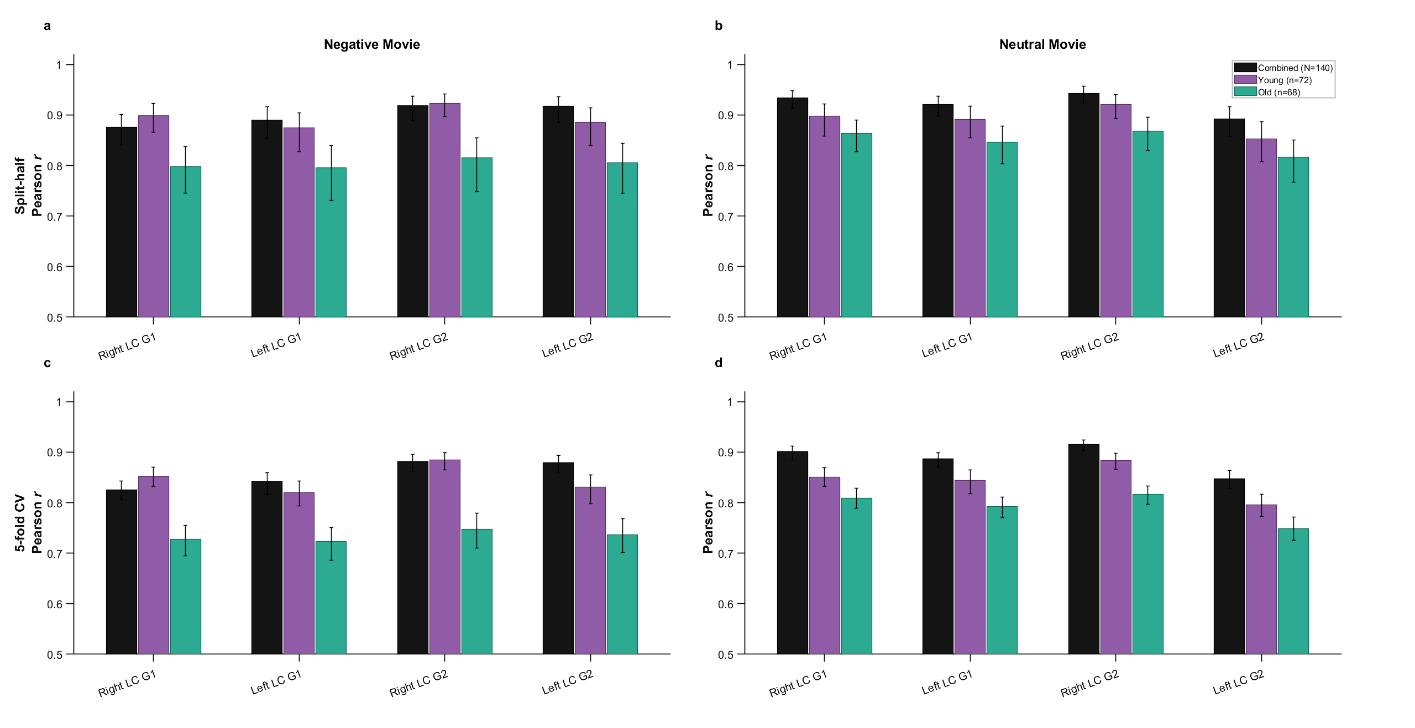
**Supplementary Figure 18**

***Cross-validation reliability of LC–linked cortical gradients across age groups and movie-viewing conditions.*** *Bar plots show Pearson correlation coefficients quantifying gradient reliability for the combined sample (N = 140, black), younger adults (n = 72, purple), and older adults (n = 68, teal), separately for the right LC G1, left LC G1, right LC G2, and left LC G2 (x-axis). Top row (****a, b****): split-half cross-validation (1,000 iterations). Bottom row (****c, d****): 5-fold cross-validation (200 iterations). Left column (****a, c****): negative movie-viewing condition. Right column (****b, d****): neutral movie-viewing condition. Error bars indicate 95% confidence intervals across iterations.*

**Supplementary Figure 1**
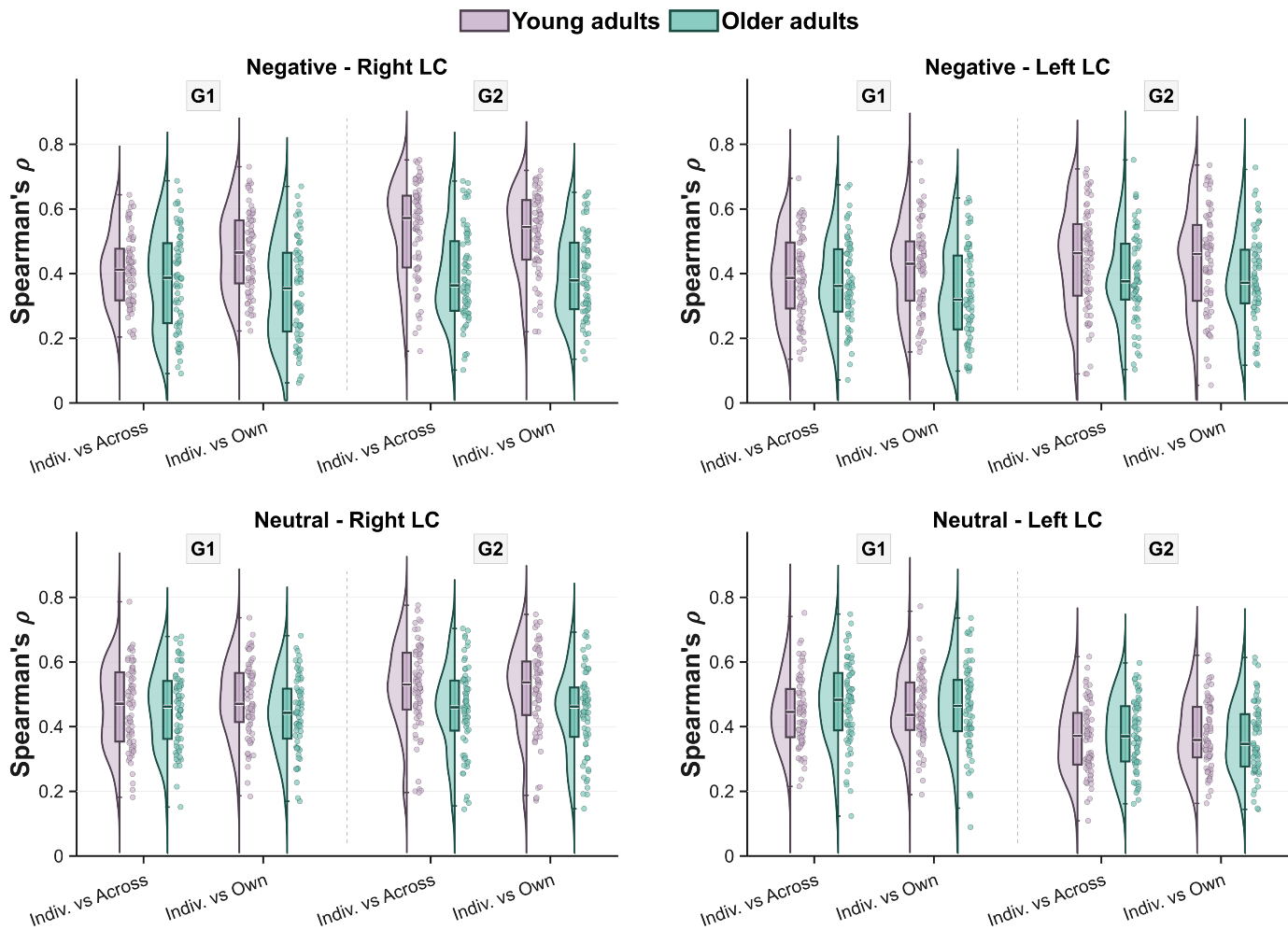
**9**

***Individual-to-group LC–cortex gradient similarity at the participant level, across conditions, hemispheres, gradients, and age group****s. Raincloud plots showing the distribution of leave-one-out (LOO) Spearman correlations between each participant's parcel-wise gradient vector and a reference group-mean gradient computed from the remaining participants. Top row: negative movie-viewing; bottom row: neutral movie-viewing, each with right LC (left) and left LC (right). Within each panel, Gradient 1 (G1) and Gradient 2 (G2) are shown side by side, with two reference comparisons: vs Combined — each participant's gradient correlation with the mean of all remaining participants (N − 1 = 139); and vs Own-group — each participant's gradient correlated with the same-age-group LOO mean (younger adults: n − 1 = 71; older adults: n − 1 = 67). Each dot represents one participant; box plots show the median and interquartile range. Younger adults are shown in purple, older adults in teal.*

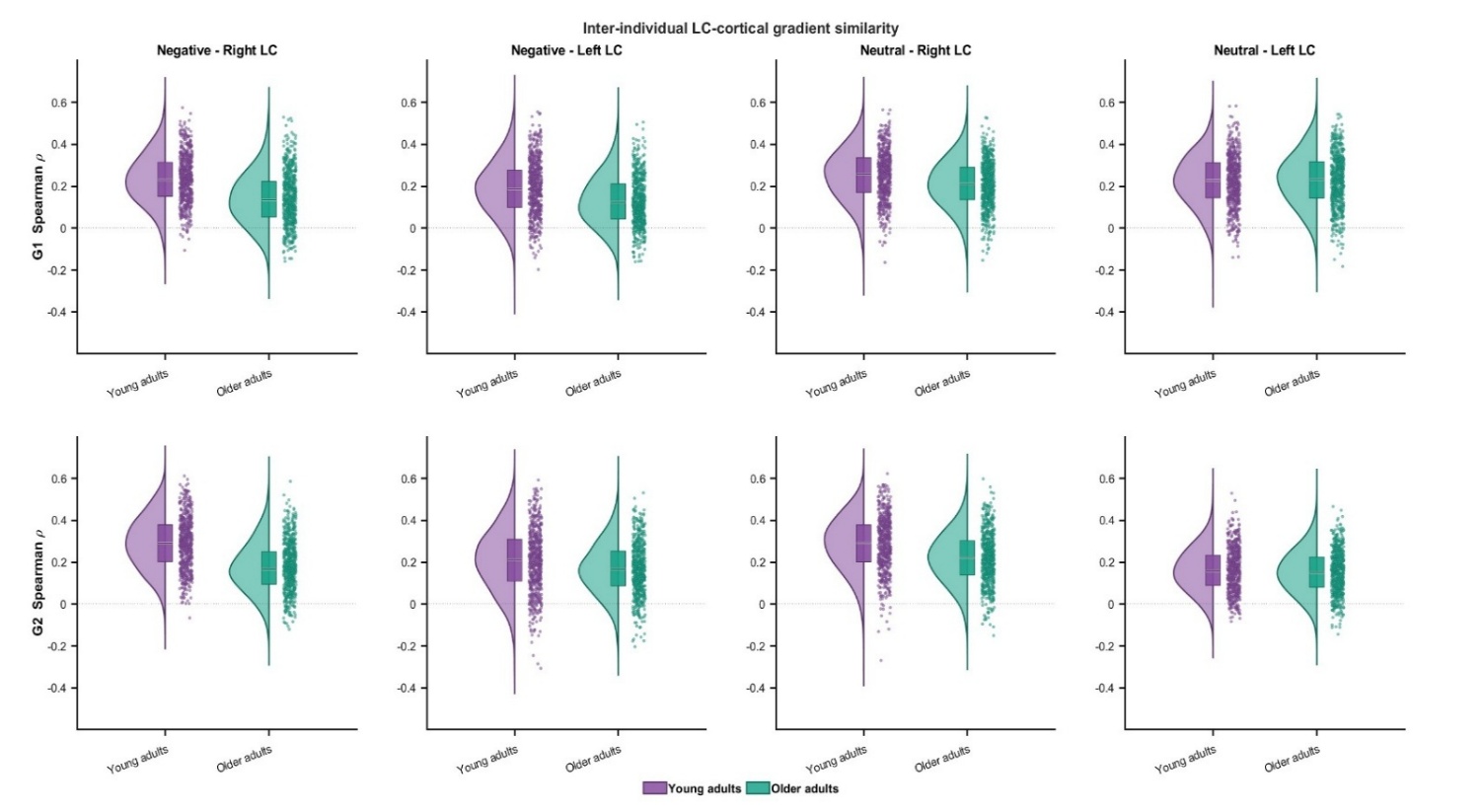
**Supplementary Figure 20**

**Inter-individual similarity of LC–linked cortical gradient organization Visualization of the reliability values reported in Supplementary Table 4.** Raincloud plots showing the distribution of pairwise Spearman correlations between participants' parcel-wise gradient score vectors, computed across all unique within-group participant pairs (younger adults, n = 2,556 pairs, purple; older adults, n = 2,278 pairs, teal). Top row: Gradient 1 (G1). Bottom row: Gradient 2 (G2). Columns from left to right: Negative–Right LC, Negative–Left LC, Neutral–Right LC, Neutral–Left LC. Each dot represents one participant pair; box plots show the median and interquartile range, and density curves show the distribution shape.

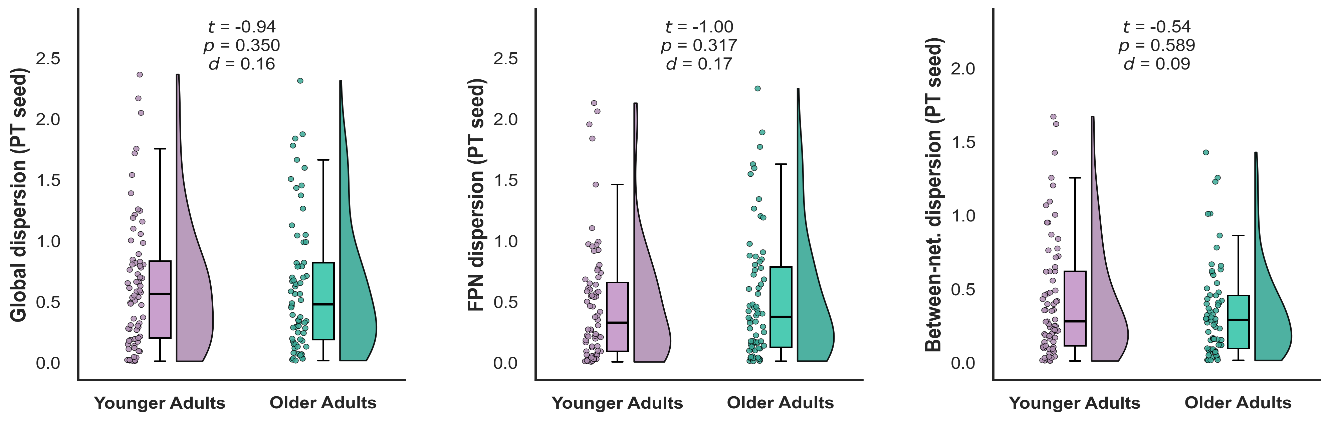
**Supplementary Figure 21**

**Cortical gradient dispersion relative to the pontine control seed between age groups.** Raincloud plots of subject-level dispersion in younger (purple) and older (teal) adults for three dispersion metrics derived from the pontine (PT) control-seed cortical gradient manifold (G1 × G2) during negative movie-viewing. Left: global dispersion. Middle: within-network dispersion for the frontoparietal network. Right: between-network dispersion. Dots are individual participants; boxes show the median and interquartile range; density curves show the distribution. Inset values give the independent-samples t, p and Cohen's d for the age-group comparison on covariate-adjusted values, as in the main LC analyses.

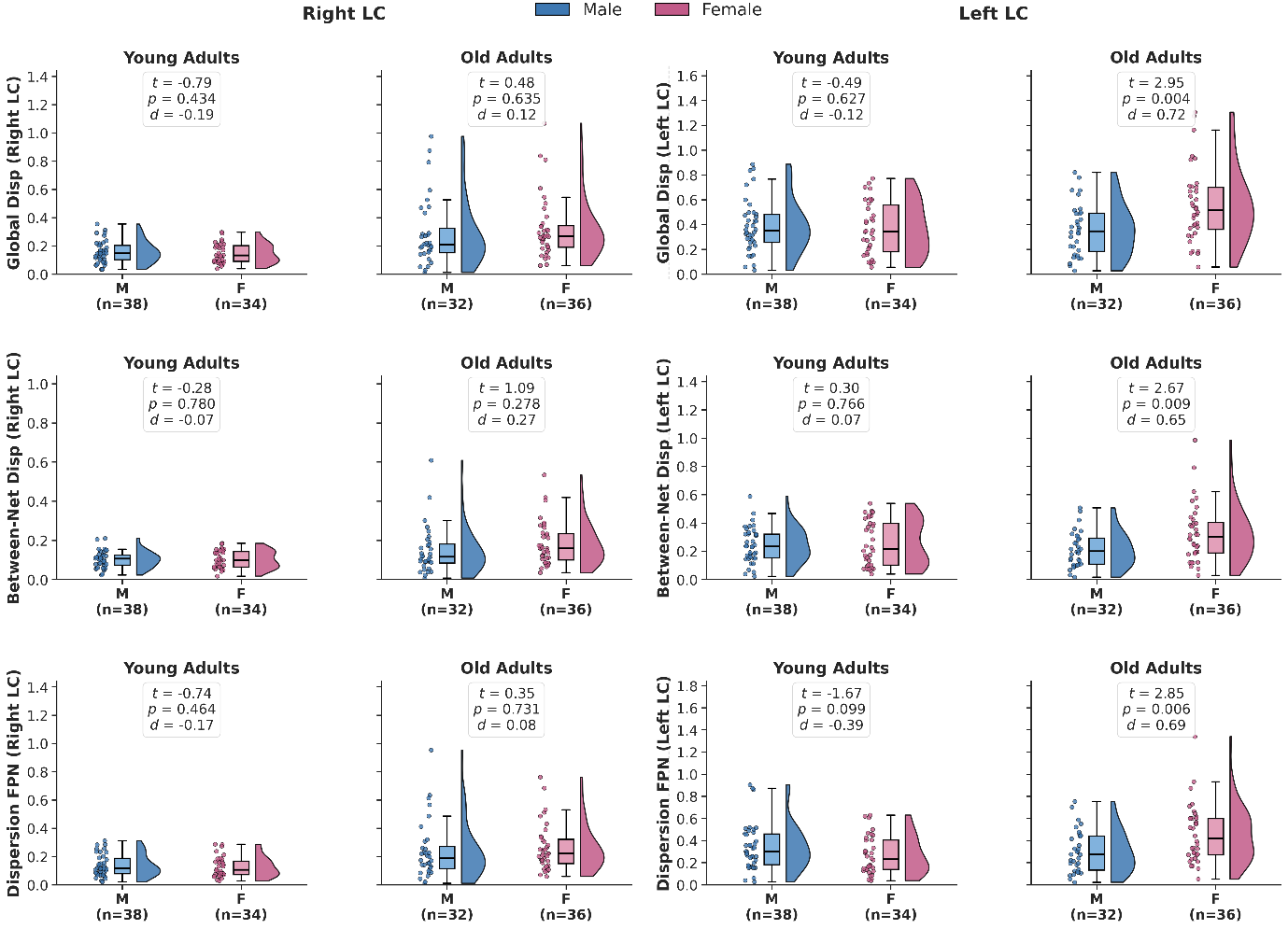
**Supplementary Figure 22**

**Sex differences in LC gradient dispersion within age groups.** Raincloud plots (half-violin, boxplot, and individual points) show six gradient-dispersion measures for males (M, blue) and females (F, pink), separately for younger and older adults. Rows show between-network, global, and within-frontoparietal-network (FPN) dispersion for the left- and right-LC gradients; per-group sample sizes are given below each panel. Sex differences were tested per measure and age group with an ANCOVA adjusting for mean framewise displacement and cortical thickness. Plotted p values are raw; annotated statistics are covariate-adjusted.

**Supplementary Figure 23**

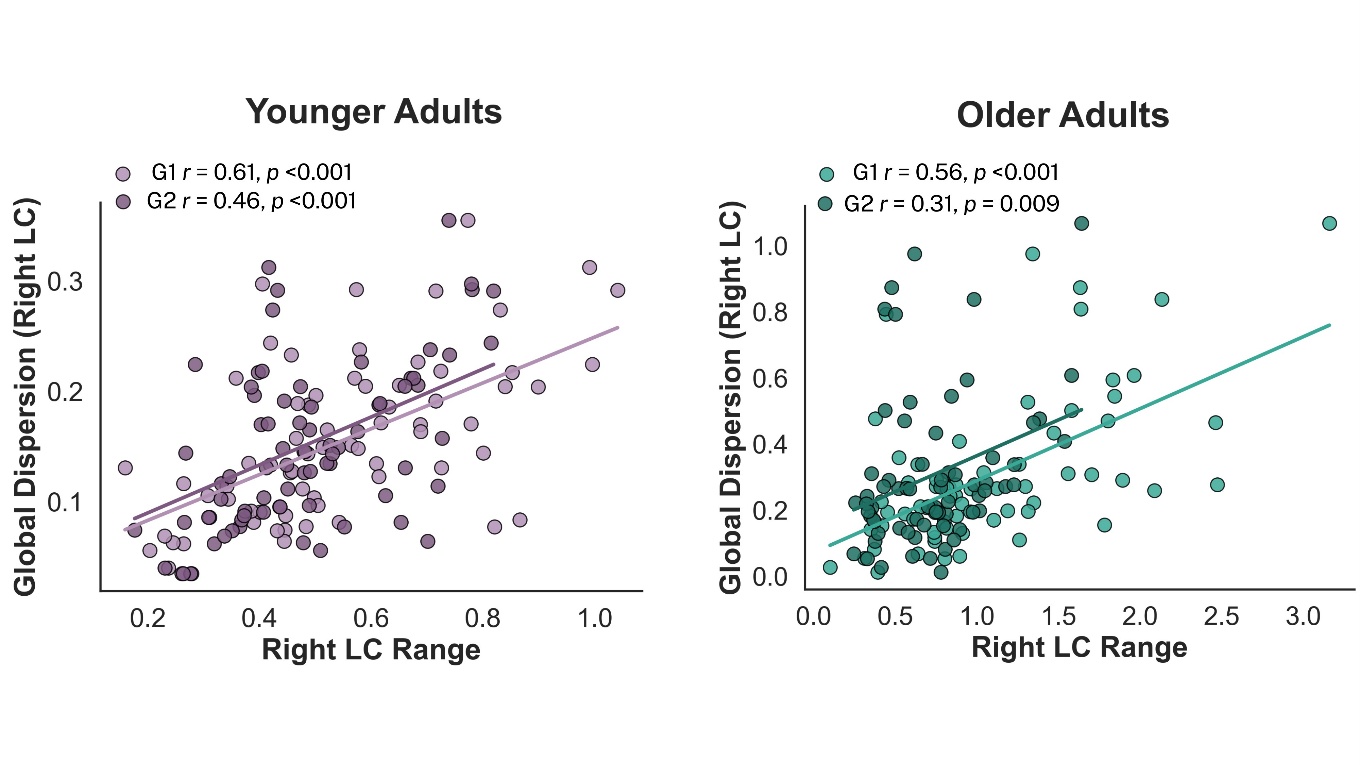

***Within-group associations between per-axis gradient range and 2D right LC–linked cortical dispersion in younger and older adults during negative movie-viewing.*** *Scatter plots showing the association between subject-level gradient range (max − min) and 2D right LC–linked cortical dispersion, separately for Gradient 1 (G1; light shade) and Gradient 2 (G2; dark shade), in younger adults (purple; left) and older adults (teal; right). Each dot represents one participant; solid lines show the linear fit per gradient. In both age groups, G1 and G2 ranges are each significantly and positively associated with 2D dispersion, indicating that both gradient axes contribute independently to the 2D dispersion measure within each age group, ruling out single-axis dominance.*

**Supplementary Figure 24**

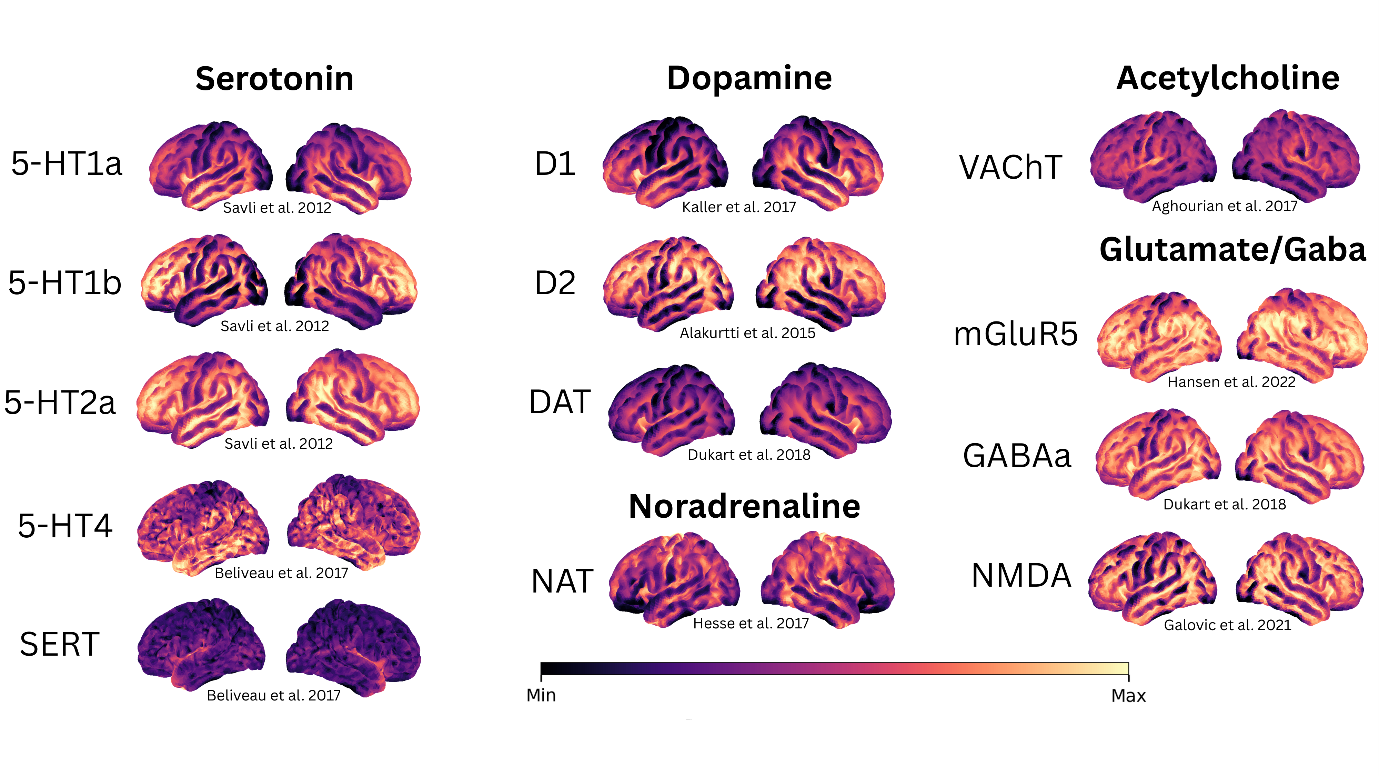

**PET-derived neurotransmitter receptor and transporter density maps used for chemoarchitectural decoding.** Cortical surface projections of the 13 PET-derived receptor and transporter density maps used to characterize the neurochemical organization underlying the LC–linked cortical gradients. Maps are grouped by neurotransmitter system: serotonin (5-HT₁ₐ, 5-HT₁ᵦ, 5-HT₂ₐ, 5-HT₄, SERT), dopamine (D₁, D₂, DAT), noradrenaline (NAT), acetylcholine (VAChT), and glutamate/GABA (mGluR₅, GABAₐ, NMDA). Each map represents the group-averaged density distribution across an independent sample of healthy adults (**Supplementary table 1**). Color reflects normalized receptor or transporter density (min–max scaled per map; shared colorbar).

**Supplementary Figure 25**

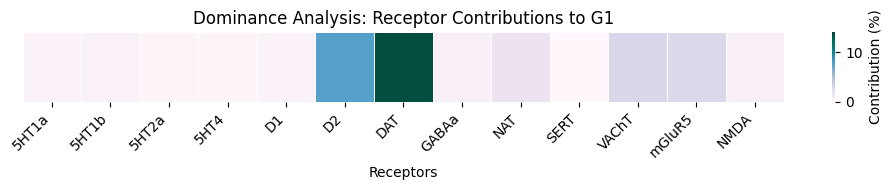
**a. Right LC-Cortex Negative**

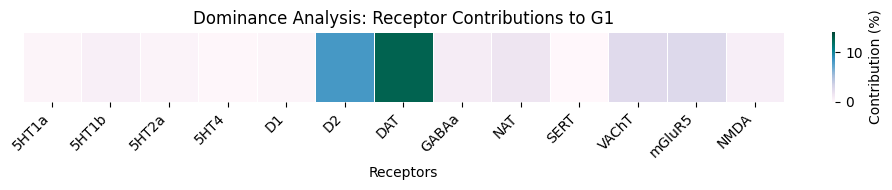
**b. Left LC-Cortex Negative**

***Interactional Dominance Results Negative Movie.*** *Figure illustrates the interactional dominance contributions of 13 cortical neurotransmitter receptor and transporter profiles to the LC–cortical Gradient 1 during negative movie-viewing. Interactional dominance quantifies the unique change in explained variance when a given neurochemical variable is added to the regression model after all other predictors are already included, capturing its distinct contribution beyond shared variance in a) Right LC-Cortex and b) Left LC-Cortex gradient map.*

**Supplementary Figure 26**

a. **Right LC-Cortex Neutral**

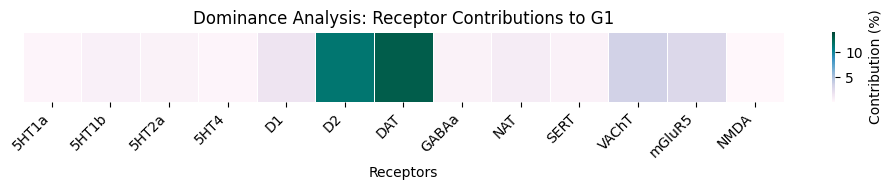

b.
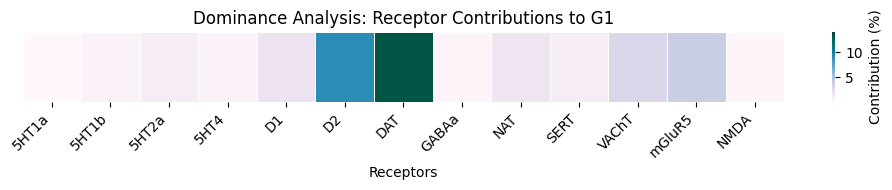
**Left LC-Cortex Neutral**

***Interactional Dominance Results Neutral Movie.*** *Figure illustrates the interactional dominance contributions of 13 cortical neurotransmitter receptor and transporter profiles to the LC–cortical Gradient 1 during neutral movie-viewing. Interactional dominance quantifies the unique change in explained variance when a given neurochemical variable is added to the regression model after all other predictors are already included, capturing its distinct contribution beyond shared variance in a) Right LC-Cortex and b) Left LC-Cortex gradient map.*

**Supplementary Tables**

*Supplementary Table 1: Methodological details of receptor*–*transporters density maps used in spatial correlation/dominance analyses*

| **Receptor Map** | **Neurotransmitter** | **Tracer** | **Modality** | **N Healthy Volunteers** | **Sex** |
| --- | --- | --- | --- | --- | --- |
| D1 | Dopamine | (^11^C) SCH23390 | PET | 13 | 7 Females |
| D2 | Dopamine | (^11^C) Raclopride | PET | 7 | All Males |
| DAT* | Dopamine | (^123^I) FP-CIT | SPECT | 174 | 65 Females |
| NAT* | Noradrenaline | (^11^C) MRB | PET | 10 | 4 Females |
| 5-HT1a | Serotonin | (^11^C) WAY-100635 | PET | 35 | 17 Females |
| 5-HT1b | Serotonin | (^11^C) P943 | PET | 23 | 8 Females |
| 5-HT2a | Serotonin | (^18^F) altanserin | PET | 19 | 8 Females |
| 5-HT4 | Serotonin | [^11^C] SB207145 | PET | 59 | 18 Females |
| SERT* | Serotonin | (^11^C) DASB | PET | 100 | 71 Females |
| VAChT | Acetylcholine | (^18^F) FEOBV | PET | 18 | 13 Females |
| mGluR5 | Glutamate | [^11^C] ABP688 | PET | 22 | 10 Females |
| NMDA | Glutamate | [^18^F] GE-179 | PET | 29 | 8 Females |
| GABAa | GABA | (^11^C) flumazenil | PET | 6 | All Males |

*Tracer abbreviations: DAT = dopamine transporter; NAT = noradrenaline transporter; SERT = serotonin transporter; VAChT = vesicular acetylcholine transporter; mGluR5 = metabotropic glutamate receptor subtype 5; NMDA = N-methyl-D-aspartate receptor; GABAa = gamma-aminobutyric acid type A receptor.* Asterisks* indicate transporters

*Supplementary Table 2: Age-related centroid displacement of Yeo networks in the LC–cortical 2D gradient manifold during negative movie-viewing.*

| ***Network*** | ***Younger adults*** | | ***Older adults*** | | ***Centroid shift (older − younger)*** | | ***\|d\|*** |
| --- | --- | --- | --- | --- | --- | --- | --- |
|  | ***G1*** | ***G2*** | ***G1*** | ***G2*** | ***ΔG1*** | ***ΔG2*** |  |
| ***Right hemisphere*** | | | | | | | |
| **VIS** | -0.0272 | -0.0445 | -0.1208 | -0.0505 | -0.0936 | -0.0060 | **0.0938** |
| **SMN** | 0.0294 | 0.0132 | 0.0637 | -0.0003 | +0.0343 | -0.0135 | **0.0369** |
| **DAN** | -0.0452 | -0.0693 | 0.0134 | -0.0919 | +0.0586 | -0.0226 | **0.0628** |
| **VAN** | 0.0264 | -0.0270 | 0.0289 | -0.0265 | +0.0025 | +0.0004 | **0.0025** |
| **LIM** | -0.0409 | 0.0351 | -0.0913 | 0.0266 | -0.0504 | -0.0085 | **0.0512** |
| **FPN** | 0.0055 | 0.0106 | 0.0248 | 0.0308 | +0.0193 | +0.0202 | **0.0279** |
| **DMN** | 0.0076 | 0.0601 | 0.0134 | 0.0795 | +0.0058 | +0.0194 | **0.0203** |
| ***Left hemisphere*** | | | | | | | |
| **VIS** | -0.0458 | -0.0258 | -0.1298 | -0.0668 | -0.0841 | -0.0410 | **0.0935** |
| **SMN** | 0.0425 | -0.0665 | 0.0926 | -0.0714 | +0.0501 | -0.0049 | **0.0503** |
| **DAN** | -0.1239 | -0.2210 | -0.0318 | -0.1745 | +0.0922 | +0.0465 | **0.1032** |
| **VAN** | -0.0032 | -0.0940 | 0.0433 | -0.0542 | +0.0465 | +0.0398 | **0.0612** |
| **LIM** | -0.0405 | 0.0898 | -0.0682 | 0.0346 | -0.0276 | -0.0553 | **0.0618** |
| **FPN** | 0.0075 | 0.0662 | 0.0247 | 0.0821 | +0.0172 | +0.0159 | **0.0234** |
| **DMN** | 0.0576 | 0.1734 | 0.0153 | 0.1729 | -0.0423 | -0.0005 | **0.0423** |

*For each Yeo seven-network parcellation, columns report the network centroid coordinates in (G1, G2) gradient space for younger adults and older adults. The per-axis displacement components ΔG1 and ΔG2 (computed as older minus younger), and the Euclidean magnitude |d| = √(ΔG1² + ΔG2²) of the displacement vector. Results are presented separately for the right LC and left LC. Positive ΔG1 and ΔG2 values indicate displacement toward the higher end of each gradient axis with age; |d| indexes the total magnitude of repositioning irrespective of direction. Network abbreviations: VIS, visual; SMN, somatomotor; DAN, dorsal attention; VAN, ventral attention; LIM, limbic; FPN, frontoparietal; DMN, default mode*

*Supplementary Table 3: Age-related centroid displacement of Yeo networks in the LC–linked cortical 2D gradient manifold during neutral movie-viewing.*

| ***Network*** | ***Younger adults*** | | ***Older adults*** | | ***Centroid shift (older − younger)*** | | ***\|d\|*** |
| --- | --- | --- | --- | --- | --- | --- | --- |
|  | ***G1*** | ***G2*** | ***G1*** | ***G2*** | ***ΔG1*** | ***ΔG2*** |  |
| ***Right hemisphere*** | | | | | | | |
| **VIS** | -0.0377 | -0.0377 | -0.0720 | -0.0477 | -0.0343 | -0.0100 | **0.0357** |
| **SMN** | -0.0203 | 0.0615 | 0.0483 | 0.0703 | +0.0686 | +0.0088 | **0.0692** |
| **DAN** | -0.1120 | -0.0287 | -0.0977 | -0.0281 | +0.0143 | +0.0007 | **0.0143** |
| **VAN** | -0.0027 | 0.0247 | 0.0152 | 0.0325 | +0.0179 | +0.0077 | **0.0195** |
| **LIM** | -0.0460 | 0.0065 | -0.0770 | 0.0081 | -0.0309 | +0.0016 | **0.0310** |
| **FPN** | 0.0351 | -0.0245 | 0.0242 | -0.0288 | -0.0109 | -0.0043 | **0.0118** |
| **DMN** | 0.0944 | -0.0090 | 0.0538 | -0.0139 | -0.0407 | -0.0049 | **0.0409** |
| ***Left hemisphere*** | | | | | | | |
| **VIS** | -0.0144 | 0.0121 | -0.0435 | -0.0066 | -0.0291 | -0.0187 | **0.0346** |
| **SMN** | -0.0015 | 0.0605 | 0.0636 | 0.0576 | +0.0650 | -0.0029 | **0.0651** |
| **DAN** | -0.0898 | 0.0149 | -0.0981 | -0.0267 | -0.0083 | -0.0417 | **0.0425** |
| **VAN** | 0.0130 | 0.0250 | 0.0182 | 0.0263 | +0.0053 | +0.0013 | **0.0054** |
| **LIM** | -0.0584 | 0.0176 | -0.0474 | 0.0411 | +0.0110 | +0.0235 | **0.0259** |
| **FPN** | 0.0079 | -0.0578 | -0.0118 | -0.0587 | -0.0197 | -0.0009 | **0.0198** |
| **DMN** | 0.0569 | -0.0541 | 0.0354 | -0.0182 | -0.0216 | +0.0359 | **0.0419** |

*For each Yeo seven-network parcellation, columns report the network centroid coordinates in (G1, G2) gradient space for younger adults and older adults. The per-axis displacement components ΔG1 and ΔG2 (computed as older minus younger), and the Euclidean magnitude |d| = √(ΔG1² + ΔG2²) of the displacement vector. Results are presented separately for the right LC and left LC. Positive ΔG1 and ΔG2 values indicate displacement toward the higher end of each gradient axis with age; |d| indexes the total magnitude of repositioning irrespective of direction. Network abbreviations: VIS, visual; SMN, somatomotor; DAN, dorsal attention; VAN, ventral attention; LIM, limbic; FPN, frontoparietal; DMN, default mode*

*Supplementary Table 4: Reliability of LC–linked cortical gradients across validation methods, age groups, and movie-viewing conditions*

| **Method** | **Group** | **Right LC G1** | **Right LC G2** | **Left LC G1** | **Left LC G2** |
| --- | --- | --- | --- | --- | --- |
| **Negative Movie** | | | | | |
| **Split-half** | **Combined** | 0.88 [0.84, 0.90] | 0.92 [0.89, 0.94] | 0.89 [0.85, 0.92] | 0.92 [0.89, 0.94] |
| **(1000 iter)** | Young | 0.90 [0.87, 0.92] | 0.92 [0.90, 0.94] | 0.88 [0.83, 0.91] | 0.88 [0.83, 0.91] |
|  | Old | 0.80 [0.74, 0.83] | 0.82 [0.75, 0.86] | 0.80 [0.74, 0.84] | 0.81 [0.75, 0.85] |
| **5-fold CV** | **Combined** | 0.83 [0.81, 0.85] | 0.88 [0.86, 0.90] | 0.84 [0.82, 0.86] | 0.88 [0.86, 0.89] |
| **(200 iter)** | Young | 0.85 [0.83, 0.87] | 0.89 [0.87, 0.90] | 0.82 [0.80, 0.84] | 0.83 [0.81, 0.86] |
|  | Old | 0.73 [0.70, 0.76] | 0.75 [0.72, 0.78] | 0.72 [0.69, 0.75] | 0.74 [0.70, 0.76] |
| **Inter-individual variability** | **Combined** | 0.17 [−0.25, 0.60] | 0.23 [−0.19, 0.64] | 0.14 [−0.40, 0.61] | 0.19 [−0.31, 0.62] |
|  | Young | 0.23 [−0.14, 0.60] | 0.29 [−0.09, 0.64] | 0.19 [−0.29, 0.61] | 0.21 [−0.31, 0.62] |
|  | Old | 0.14 [−0.21, 0.55] | 0.18 [−0.17, 0.59] | 0.13 [−0.22, 0.55] | 0.17 [−0.22, 0.59] |
| **Neutral Movie** | | | | | |
| **Split-half** | **Combined** | 0.93 [0.91, 0.95] | 0.94 [0.92, 0.96] | 0.92 [0.90, 0.94] | 0.89 [0.86, 0.91] |
| **(1000 iter)** | Young | 0.90 [0.86, 0.92] | 0.92 [0.89, 0.94] | 0.89 [0.85, 0.92] | 0.85 [0.81, 0.88] |
|  | Old | 0.86 [0.83, 0.89] | 0.87 [0.83, 0.90] | 0.85 [0.81, 0.88] | 0.82 [0.77, 0.85] |
| **5-fold CV** | **Combined** | 0.90 [0.89, 0.91] | 0.92 [0.90, 0.93] | 0.89 [0.87, 0.90] | 0.85 [0.83, 0.87] |
| **(200 iter)** | Young | 0.85 [0.83, 0.87] | 0.88 [0.87, 0.90] | 0.84 [0.82, 0.86] | 0.80 [0.78, 0.82] |
|  | Old | 0.81 [0.79, 0.83] | 0.82 [0.80, 0.84] | 0.79 [0.77, 0.81] | 0.75 [0.72, 0.77] |
| **Inter-individual variability** | **Combined** | 0.22 [−0.29, 0.61] | 0.24 [−0.27, 0.62] | 0.22 [−0.42, 0.60] | 0.14 [−0.25, 0.53] |
|  | Young | 0.25 [−0.20, 0.60] | 0.29 [−0.27, 0.62] | 0.23 [−0.26, 0.58] | 0.16 [−0.14, 0.53] |
|  | Old | 0.21 [−0.18, 0.56] | 0.22 [−0.19, 0.60] | 0.23 [−0.18, 0.60] | 0.15 [−0.17, 0.53] |

*Supplementary Table 5. Leave-one-out individual-to-group gradient spatial similarity across participants, age groups, gradients, and movie-viewing conditions.*

| **Condition** | **Gradient** | **Reference** | **Group** | **Mean *ρ*** | **Median *ρ*** | **SD** | **Min *ρ*** | **Max *ρ*** |
| --- | --- | --- | --- | --- | --- | --- | --- | --- |
| **Negative Movie — Right LC** | | | | | | | | |
|  | **G1** | *Individual vs across-group* | **Young** | 0.41 | 0.41 | 0.11 | 0.20 | 0.64 |
|  |  | *Individual vs across-group* | Old | 0.38 | 0.39 | 0.15 | 0.09 | 0.69 |
|  |  | *Individual vs Own-group* | **Young** | 0.47 | 0.47 | 0.13 | 0.22 | 0.73 |
|  |  | *Individual vs Own-group* | Old | 0.35 | 0.36 | 0.16 | 0.06 | 0.67 |
|  | **G2** | *Individual* vs across-group | **Young** | 0.54 | 0.57 | 0.15 | 0.16 | 0.75 |
|  |  | *Individual* vs across-group | Old | 0.39 | 0.36 | 0.14 | 0.10 | 0.69 |
|  |  | *Individual vs Own-group* | **Young** | 0.53 | 0.54 | 0.12 | 0.22 | 0.72 |
|  |  | *Individual vs Own-group* | Old | 0.39 | 0.38 | 0.13 | 0.14 | 0.65 |
| **Negative Movie — Left LC** | | | | | | | | |
|  | **G1** | *Individual* vs across-group | **Young** | 0.39 | 0.39 | 0.13 | 0.14 | 0.70 |
|  |  | *Individual* vs across-group | Old | 0.37 | 0.36 | 0.13 | 0.07 | 0.68 |
|  |  | *Individual vs Own-group* | **Young** | 0.42 | 0.43 | 0.14 | 0.16 | 0.75 |
|  |  | *Individual vs Own-group* | Old | 0.34 | 0.32 | 0.15 | 0.10 | 0.63 |
|  | **G2** | *Individual* vs across-group | **Young** | 0.45 | 0.46 | 0.16 | 0.09 | 0.72 |
|  |  | *Individual* vs across-group | Old | 0.39 | 0.38 | 0.14 | 0.10 | 0.75 |
|  |  | *Individual vs Own-group* | **Young** | 0.44 | 0.46 | 0.16 | 0.06 | 0.74 |
|  |  | *Individual vs Own-group* | Old | 0.39 | 0.37 | 0.14 | 0.12 | 0.73 |
| **Neutral Movie — Right LC** | | | | | | | | |
|  | **G1** | *Individual* vs across-group | Young | 0.46 | 0.47 | 0.13 | 0.18 | 0.79 |
|  |  | *Individual* vs across-group | Old | 0.46 | 0.46 | 0.12 | 0.15 | 0.68 |
|  |  | *Individual vs Own-group* | Young | 0.48 | 0.47 | 0.12 | 0.18 | 0.74 |
|  |  | *Individual vs Own-group* | Old | 0.44 | 0.44 | 0.11 | 0.17 | 0.68 |
|  | **G2** | *Individual* vs across-group | Young | 0.52 | 0.53 | 0.14 | 0.20 | 0.78 |
|  |  | *Individual* vs across-group | Old | 0.45 | 0.46 | 0.13 | 0.15 | 0.70 |
|  |  | *Individual vs Own-group* | Young | 0.52 | 0.54 | 0.14 | 0.17 | 0.75 |
|  |  | *Individual vs Own-group* | Old | 0.44 | 0.46 | 0.13 | 0.15 | 0.69 |
| **Neutral Movie — Left LC** | | | | | | | | |
|  | **G1** | *Individual* vs across-group | Young | 0.45 | 0.45 | 0.11 | 0.22 | 0.75 |
|  |  | *Individual* vs across-group | Old | 0.47 | 0.48 | 0.13 | 0.12 | 0.75 |
|  |  | *Individual vs Own-group* | Young | 0.45 | 0.44 | 0.11 | 0.19 | 0.77 |
|  |  | *Individual vs Own-group* | Old | 0.46 | 0.46 | 0.13 | 0.09 | 0.74 |
|  | **G2** | *Individual* vs across-group | Young | 0.36 | 0.37 | 0.11 | 0.11 | 0.62 |
|  |  | *Individual* vs across-group | Old | 0.38 | 0.37 | 0.11 | 0.16 | 0.60 |
|  |  | *Individual vs Own-group* | Young | 0.45 | 0.45 | 0.11 | 0.22 | 0.75 |
|  |  | *Individual vs Own-group* | Old | 0.37 | 0.35 | 0.11 | 0.14 | 0.61 |

*Supplementary Table 6. Correlations between Right-LC dispersion and individual gradient reliability (leave-one-out ρ) during negative movie-viewing*

| **Dispersion metric** | **Gradient** | **Younger (n = 72) ρ (p)** | **Older (n = 68) ρ (p)** |
| --- | --- | --- | --- |
| Global | G1 | -0.008 (0.949) | 0.189 (0.123) |
| Global | G2 | 0.145 (0.223) | −0.079 (0.522) |
| Between-network | G1 | 0.051 (0.671) | 0.195 (0.110) |
| Between-network | G2 | 0.182 (0.125) | 0.027 (0.830) |
| FPN | G1 | 0.049 (0.686) | 0.180 (0.142) |
| FPN | G2 | 0.124 (0.299) | −0.074 (0.551) |

*Supplementary Table 7.* *Age-group differences in right-LC dispersion with and without individual gradient reliability as a covariate*

| **Dispersion metric** | **Covariates** | ***t*** | ***p*** | ***Cohen's d*** |
| --- | --- | --- | --- | --- |
| Global | sex, FD, thickness | −2.94 | 0.004 | 0.50 |
| **Global** | **+ LOO G1, G2** | **−2.82** | **0.006** | **0.48** |
| Between-network | sex, FD, thickness | −2.42 | 0.017 | 0.41 |
| Between-network | **+ LOO G1, G2** | **−2.35** | **0.020** | **0.40** |
| FPN | sex, FD, thickness | −2.82 | 0.006 | 0.48 |
| FPN | **+ LOO G1, G2** | **−2.68** | **0.008** | **0.46** |

Note. "+ LOO G1, G2": same model with leave-one-out reliability (ρ) of G1 and G2 added as additional covariates

*Supplementary Table 8****.*** *Associations between right-LC dispersion and the Emotional Resilience Index with individual gradient reliability as an additional covariate*

| **Dispersion metric** | **Age × dispersion β [95% CI], *p***  **Original covariates** | **Age × dispersion β [95% CI], *p***  **After extra LOO covariates** |
| --- | --- | --- |
| Global | 0.62 [0.11, 1.14], 0.018 | 0.61 [0.09, 1.14], 0.021 |
| FPN | 0.46 [0.03, 0.89], 0.037 | 0.45 [0.01, 0.89], 0.045 |

*Supplementary Table 9: Sensitivity analysis showing spatial correlations between full-sample and excluded-sample right LC–linked cortical gradients after removing the highest-dispersion individuals in negative movie-viewing.*

| **Comparison** | **Gradient** | **Spatial correlation (ρ)** |
| --- | --- | --- |
| Combined (N = 140) vs Excluded 5% (N = 133) | G1 | 0.97 |
|  | G2 | 0.99 |
| Combined (N = 140) vs Excluded 10% (N = 126) | G1 | 0.94 |
|  | G2 | 0.98 |
| Older adults (N = 68) vs Excluded 5% OA (N = 61) | G1 | 0.93 |
|  | G2 | 0.98 |
| Older adults (N = 68) vs Excluded 10% OA (N = 54) | G1 | 0.88 |
|  | G2 | 0.94 |

*ρ = Spearman spatial correlation between gradient maps. G1 = principal gradient; G2 = secondary gradient. OA = older adults.*

*Supplementary Table 10: Sensitivity analysis comparing dispersion metrics derived from 2D (G1–G2) versus 3D (G1–G3) gradient solutions, for the right LC–cortex coupling during negative movie-viewing.*

| **Metric** | **2D (G1–G2)** | | | | **3D (G1–G3)** | | | |
| --- | --- | --- | --- | --- | --- | --- | --- | --- |
|  | ***t*** | ***p*** | ***d*** | ***95% CI*** | ***t*** | ***p*** | ***d*** | ***95% CI*** |
| **Global Dispersion** | −2.94 | 0.004 | 0.50 | [−0.145, −0.028] | −2.44 | 0.016 | 0.41 | [−0.161, −0.016] |
| **Between-Network** | −2.42 | 0.017 | 0.41 | [−0.066, −0.007] | −2.05 | 0.042 | 0.35 | [−0.060, −0.001] |
| **FPN Within-Network** | −2.82 | 0.006 | 0.48 | [−0.117, −0.021] | −2.27 | 0.025 | 0.38 | [−0.134, −0.009] |

*Supplementary Table 11: Sensitivity analysis comparing dispersion metrics derived from 2D (G1–G2) versus 3D (G1–G3) gradient solutions, for the Left LC–cortex coupling during negative movie-viewing.*

| **Metric** | **2D (G1–G2)** | | | | **3D (G1–G3)** | | | |
| --- | --- | --- | --- | --- | --- | --- | --- | --- |
|  | ***t*** | ***p*** | ***d*** | ***95% CI*** | ***t*** | ***p*** | ***d*** | ***95% CI*** |
| **Global Dispersion** | −1.70 | 0.092 | 0.29 | [−0.152, 0.012] | −1.81 | 0.074 | 0.30 | [−0.157, 0.008] |
| **Between-Network** | −1.44 | 0.152 | 0.24 | [−0.090, 0.014] | −1.51 | 0.136 | 0.25 | [−0.092, 0.012] |
| **FPN Within-Network** | −1.70 | 0.091 | 0.29 | [−0.136, 0.010] | −1.80 | 0.074 | 0.30 | [−0.140, 0.007] |

*Supplementary Table 12: LC–Cortex FC correlation with mean framewise luminance across participants.*

| **Movie** | **LC Side** | **r** | **95% CI** | **p-permutation** |
| --- | --- | --- | --- | --- |
| Negative | Left | +0.049 | [−0.036, +0.212] | 0.508 |
| Negative | Right | −0.012 | [−0.214, +0.171] | 0.917 |
| Neutral | Left | −0.031 | [−0.205, +0.102] | 0.836 |
| Neutral | Right | +0.003 | [−0.204, +0.207] | 0.988 |

*Supplementary Table 13: LC–cortex FC correlation with luminance by age group.*

| **Movie** | **LC Side** | **Group** | **r** | **95% CI** | **p-permutation** |
| --- | --- | --- | --- | --- | --- |
| Negative | Left | Young | −0.102 | [−0.208, +0.180] | 0.458 |
| Negative | Left | Old | +0.158 | [−0.030, +0.294] | 0.192 |
| Negative | Right | Young | −0.034 | [−0.234, +0.209] | 0.836 |
| Negative | Right | Old | +0.008 | [−0.193, +0.206] | 0.949 |
| Neutral | Left | Young | −0.074 | [−0.225, +0.102] | 0.424 |
| Neutral | Left | Old | +0.031 | [−0.221, +0.238] | 0.760 |
| Neutral | Right | Young | −0.011 | [−0.180, +0.165] | 0.911 |
| Neutral | Right | Old | +0.018 | [−0.163, +0.200] | 0.840 |

*Supplementary Table 14***.** *Age × hemisphere interaction for LC–cortical gradient dispersion during negative movie-viewing, with hemispheres as repeated measures within participants*

| **Dispersion metric** | **Age × Hemisphere d [95% CI], *p*** |
| --- | --- |
| Global | 0.07 [−0.26, 0.40], 0.682 |
| Between-network | −0.01 [−0.34, 0.32], 0.934 |
| FPN | 0.03 [−0.30, 0.36], 0.864 |

*Supplementary Table 15: Correlations between right LC–cortical dispersion and mean framewise displacement during negative movie-viewing*

| **Metric** | **Overall *r*** | **Older adults *r*** |
| --- | --- | --- |
| **Global dispersion** | 0.16 | 0.22 |
| **Between-network Dispersion** | 0.17 | 0.34 |
| **FPN Dispersion** | 0.14 | 0.24 |

*Supplementary Table 16: Spearman rank correlations between within-network dispersion and global dispersion during negative movie-viewing, across Yeo seven-networks, hemispheres, and age groups*

| **Network** | **Hemisphere** | **All Participants** | | **Younger Adults** | | **Older Adults** | |
| --- | --- | --- | --- | --- | --- | --- | --- |
|  |  | ***ρ*** | ***n*** | ***ρ*** | ***n*** | ***ρ*** | ***n*** |
| Visual | Left | 0.82 | 140 | 0.87 | 72 | 0.75 | 68 |
|  | Right | 0.90 | 140 | 0.91 | 72 | 0.84 | 68 |
| Somatomotor | Left | 0.92 | 140 | 0.91 | 72 | 0.93 | 68 |
|  | Right | 0.89 | 140 | 0.93 | 72 | 0.87 | 68 |
| Dorsal Attention | Left | 0.84 | 140 | 0.84 | 72 | 0.84 | 68 |
|  | Right | 0.88 | 140 | 0.87 | 72 | 0.84 | 68 |
| Ventral Attention | Left | 0.94 | 140 | 0.95 | 72 | 0.94 | 68 |
|  | Right | 0.93 | 140 | 0.93 | 72 | 0.89 | 68 |
| Limbic | Left | 0.86 | 140 | 0.83 | 72 | 0.85 | 68 |
|  | Right | 0.87 | 140 | 0.80 | 72 | 0.84 | 68 |
| Frontoparietal | Left | 0.89 | 140 | 0.87 | 72 | 0.91 | 68 |
|  | Right | 0.95 | 140 | 0.94 | 72 | 0.93 | 68 |
| Default Mode | Left | 0.78 | 140 | 0.68 | 72 | 0.83 | 68 |
|  | Right | 0.90 | 140 | 0.82 | 72 | 0.89 | 68 |

### *Supplementary Table 17: Neurosynth functional decoding for the left LC–cortex G1 during neutral movie-viewing, showing spin-test (p_spin_) and FDR-corrected association terms*

| **Term** | **ρ (rho)** | **p_spin** | **p_spin (FDR)** | **Correction** |
| --- | --- | --- | --- | --- |
| **Adaptation** | -0.452 | 5.00e-4 | 0.0154 | **both** |
| **Expertise** | -0.436 | 4.00e-4 | 0.0154 | **both** |
| **Strength** | 0.435 | 4.00e-4 | 0.0154 | **both** |
| **Discrimination** | -0.413 | 4.00e-4 | 0.0154 | **both** |
| **Inference** | 0.291 | 0.0014 | 0.0344 | **both** |
| Objectrecognition | -0.448 | 0.0182 | 0.1078 | p_spin only |
| Risk | 0.429 | 0.0082 | 0.0776 | p_spin only |
| Stress | 0.404 | 0.0113 | 0.0927 | p_spin only |
| Perception | -0.402 | 0.0316 | 0.1589 | p_spin only |
| Reading | -0.395 | 0.0057 | 0.0701 | p_spin only |
| Gaze | -0.393 | 0.0134 | 0.1030 | p_spin only |
| Psychosis | 0.383 | 0.0045 | 0.0701 | p_spin only |
| Association | 0.374 | 0.0029 | 0.0594 | p_spin only |
| Anticipation | 0.372 | 0.0150 | 0.1078 | p_spin only |
| Hyperactivity | 0.346 | 0.0056 | 0.0701 | p_spin only |
| Salience | 0.345 | 0.0076 | 0.0776 | p_spin only |
| Inhibition | 0.343 | 0.0184 | 0.1078 | p_spin only |
| Loss | 0.325 | 0.0336 | 0.1589 | p_spin only |
| Induction | 0.319 | 0.0173 | 0.1078 | p_spin only |
| Sleep | 0.318 | 0.0080 | 0.0776 | p_spin only |
| Mentalimagery | -0.304 | 0.0378 | 0.1682 | p_spin only |
| Selectiveattention | -0.289 | 0.0308 | 0.1589 | p_spin only |
| Naming | -0.270 | 0.0053 | 0.0701 | p_spin only |
| Balance | 0.269 | 0.0206 | 0.1152 | p_spin only |
| Consolidation | 0.268 | 0.0336 | 0.1589 | p_spin only |
| Insight | 0.266 | 0.0166 | 0.1078 | p_spin only |
| Competition | -0.213 | 0.0383 | 0.1682 | p_spin only |
| Concept | 0.204 | 0.0093 | 0.0817 | p_spin only |

### *Supplementary Table 18: Neurosynth functional decoding for the left LC–cortex G2 during neutral movie-viewing, showing spin-test (p_spin_) and FDR-corrected association terms*

| **Term** | **ρ (rho)** | **p_spin** | **p_spin (FDR)** | **Correction** |
| --- | --- | --- | --- | --- |
| **Reasoning** | -0.704 | 4.00e-4 | 0.0035 | **both** |
| **Cognitivecontrol** | -0.698 | 4.00e-4 | 0.0035 | **both** |
| **Memory** | -0.683 | 4.00e-4 | 0.0035 | **both** |
| **Memoryretrieval** | -0.666 | 4.00e-4 | 0.0035 | **both** |
| **Judgment** | -0.633 | 4.00e-4 | 0.0035 | **both** |
| **Retrieval** | -0.623 | 4.00e-4 | 0.0035 | **both** |
| **Strategy** | -0.569 | 4.00e-4 | 0.0035 | **both** |
| **Workingmemory** | -0.564 | 4.00e-4 | 0.0035 | **both** |
| **Perception** | 0.556 | 4.00e-4 | 0.0035 | **both** |
| **Rhythm** | 0.534 | 5.00e-4 | 0.0035 | **both** |
| **Decision** | -0.531 | 4.00e-4 | 0.0035 | **both** |
| **Maintenance** | -0.527 | 6.00e-4 | 0.0035 | **both** |
| **Decisionmaking** | -0.517 | 6.00e-4 | 0.0035 | **both** |
| **Episodicmemory** | -0.513 | 5.00e-4 | 0.0035 | **both** |
| **Rule** | -0.502 | 6.00e-4 | 0.0035 | **both** |
| **Intention** | -0.500 | 8.00e-4 | 0.0038 | **both** |
| **Updating** | -0.497 | 4.00e-4 | 0.0035 | **both** |
| **Monitoring** | -0.490 | 7.00e-4 | 0.0037 | **both** |
| **Facialexpression** | 0.489 | 4.00e-4 | 0.0035 | **both** |
| **Navigation** | -0.483 | 8.00e-4 | 0.0038 | **both** |
| **Goal** | -0.481 | 6.00e-4 | 0.0035 | **both** |
| **Integration** | 0.463 | 7.00e-4 | 0.0037 | **both** |
| **Uncertainty** | -0.450 | 5.00e-4 | 0.0035 | **both** |
| **Speechproduction** | 0.446 | 0.0026 | 0.0097 | **both** |
| **Impulsivity** | -0.444 | 4.00e-4 | 0.0035 | **both** |
| **Multisensory** | 0.438 | 0.0033 | 0.0116 | **both** |
| **Attention** | -0.434 | 1.00e-3 | 0.0044 | **both** |
| **Thought** | -0.425 | 0.0018 | 0.0071 | **both** |
| **Autobiographicalmemory** | -0.409 | 0.0046 | 0.0149 | **both** |
| **Intelligence** | -0.407 | 9.00e-4 | 0.0041 | **both** |
| **Recall** | -0.388 | 0.0015 | 0.0061 | **both** |
| **Responseinhibition** | -0.371 | 4.00e-4 | 0.0035 | **both** |
| **Taskdifficulty** | -0.369 | 0.0042 | 0.0140 | **both** |
| **Knowledge** | -0.367 | 8.00e-4 | 0.0038 | **both** |
| **Recognition** | -0.356 | 0.0155 | 0.0433 | **both** |
| **Retention** | -0.354 | 0.0028 | 0.0101 | **both** |
| **Expectancy** | -0.335 | 0.0020 | 0.0077 | **both** |
| **Familiarity** | -0.333 | 0.0035 | 0.0120 | **both** |
| **Interference** | -0.330 | 0.0137 | 0.0401 | **both** |
| **Emotionregulation** | -0.322 | 0.0155 | 0.0433 | **both** |
| **Efficiency** | -0.295 | 0.0180 | 0.0492 | **both** |
| **Localization** | 0.279 | 0.0068 | 0.0214 | **both** |
| **Distraction** | -0.273 | 0.0014 | 0.0059 | **both** |
| **Naming** | 0.263 | 0.0130 | 0.0390 | **both** |
| **Inference** | -0.215 | 0.0103 | 0.0317 | **both** |
| Movement | 0.395 | 0.0230 | 0.0589 | p_spin only |
| Pain | 0.359 | 0.0407 | 0.0927 | p_spin only |
| Coordination | 0.357 | 0.0372 | 0.0880 | p_spin only |
| Morphology | 0.320 | 0.0490 | 0.1096 | p_spin only |
| Speechperception | 0.319 | 0.0240 | 0.0602 | p_spin only |
| Risk | -0.307 | 0.0358 | 0.0863 | p_spin only |
| Induction | 0.295 | 0.0214 | 0.0572 | p_spin only |
| Belief | -0.279 | 0.0337 | 0.0829 | p_spin only |
| Competition | -0.266 | 0.0230 | 0.0589 | p_spin only |
| Consciousness | -0.254 | 0.0388 | 0.0900 | p_spin only |

### *Supplementary Table 19: Neurosynth functional decoding for the right LC–cortex G1 during neutral movie-viewing, showing spin-test (p_spin_) and FDR-corrected association terms*

| **Term** | **ρ (rho)** | **p_spin** | **p_spin (FDR)** | **Correction** |
| --- | --- | --- | --- | --- |
| **Perception** | -0.557 | 4.00e-4 | 0.0123 | **both** |
| **Discrimination** | -0.499 | 4.00e-4 | 0.0123 | **both** |
| **Adaptation** | -0.484 | 4.00e-4 | 0.0123 | **both** |
| **Risk** | 0.479 | 0.0061 | 0.0441 | **both** |
| **Cognitivecontrol** | 0.442 | 0.0022 | 0.0283 | **both** |
| **Reading** | -0.417 | 0.0023 | 0.0283 | **both** |
| **Strength** | 0.402 | 4.00e-4 | 0.0123 | **both** |
| **Hyperactivity** | 0.393 | 8.00e-4 | 0.0164 | **both** |
| **Psychosis** | 0.386 | 0.0042 | 0.0397 | **both** |
| **Expertise** | -0.382 | 0.0017 | 0.0261 | **both** |
| **Memoryretrieval** | 0.378 | 0.0033 | 0.0359 | **both** |
| **Association** | 0.369 | 0.0054 | 0.0441 | **both** |
| **Salience** | 0.354 | 0.0069 | 0.0447 | **both** |
| **Naming** | -0.345 | 0.0012 | 0.0211 | **both** |
| **Inference** | 0.333 | 6.00e-4 | 0.0148 | **both** |
| **Belief** | 0.319 | 0.0068 | 0.0447 | **both** |
| **Uncertainty** | 0.295 | 0.0053 | 0.0441 | **both** |
| **Strategy** | 0.254 | 0.0060 | 0.0441 | **both** |
| **Concept** | 0.221 | 0.0035 | 0.0359 | **both** |
| Objectrecognition | -0.493 | 0.0166 | 0.0851 | p_spin only |
| Multisensory | -0.468 | 0.0403 | 0.1524 | p_spin only |
| Integration | -0.456 | 0.0191 | 0.0940 | p_spin only |
| Decisionmaking | 0.428 | 0.0119 | 0.0697 | p_spin only |
| Gaze | -0.418 | 0.0202 | 0.0956 | p_spin only |
| Impulsivity | 0.413 | 0.0148 | 0.0827 | p_spin only |
| Monitoring | 0.385 | 0.0317 | 0.1300 | p_spin only |
| Inhibition | 0.380 | 0.0114 | 0.0697 | p_spin only |
| Visualperception | -0.379 | 0.0284 | 0.1247 | p_spin only |
| Facialexpression | -0.371 | 0.0457 | 0.1606 | p_spin only |
| Mentalimagery | -0.315 | 0.0409 | 0.1524 | p_spin only |
| Selectiveattention | -0.303 | 0.0228 | 0.1039 | p_spin only |
| Responseinhibition | 0.298 | 0.0158 | 0.0845 | p_spin only |
| Reasoning | 0.290 | 0.0375 | 0.1488 | p_spin only |
| Sleep | 0.274 | 0.0456 | 0.1606 | p_spin only |
| Thought | 0.271 | 0.0317 | 0.1300 | p_spin only |

### *Supplementary Table 20: Neurosynth functional decoding for the right LC–cortex G2 during neutral movie-viewing, showing spin-test (p_spin_) and FDR-corrected association terms*

| **Term** | **ρ (rho)** | **p_spin** | **p_spin (FDR)** | **Correction** |
| --- | --- | --- | --- | --- |
| **Pain** | 0.752 | 5.00e-4 | 0.0068 | **both** |
| **Judgment** | -0.673 | 4.00e-4 | 0.0061 | **both** |
| **Recognition** | -0.638 | 4.00e-4 | 0.0061 | **both** |
| **Memory** | -0.600 | 4.00e-4 | 0.0061 | **both** |
| **Navigation** | -0.579 | 7.00e-4 | 0.0078 | **both** |
| **Retrieval** | -0.528 | 4.00e-4 | 0.0061 | **both** |
| **Induction** | 0.518 | 1.00e-3 | 0.0088 | **both** |
| **Attention** | -0.514 | 0.0031 | 0.0166 | **both** |
| **Encoding** | -0.492 | 0.0016 | 0.0116 | **both** |
| **Workingmemory** | -0.488 | 4.00e-4 | 0.0061 | **both** |
| **Rhythm** | 0.477 | 0.0087 | 0.0345 | **both** |
| **Expertise** | -0.471 | 6.00e-4 | 0.0074 | **both** |
| **Knowledge** | -0.467 | 8.00e-4 | 0.0082 | **both** |
| **Reasoning** | -0.458 | 9.00e-4 | 0.0085 | **both** |
| **Updating** | -0.449 | 4.00e-4 | 0.0061 | **both** |
| **Episodicmemory** | -0.447 | 0.0015 | 0.0115 | **both** |
| **Spatialattention** | -0.445 | 0.0028 | 0.0166 | **both** |
| **Anticipation** | 0.419 | 0.0052 | 0.0229 | **both** |
| **Familiarity** | -0.417 | 4.00e-4 | 0.0061 | **both** |
| **Balance** | 0.416 | 0.0046 | 0.0218 | **both** |
| **Intention** | -0.410 | 0.0028 | 0.0166 | **both** |
| **Loss** | 0.400 | 0.0017 | 0.0116 | **both** |
| **Search** | -0.397 | 0.0030 | 0.0166 | **both** |
| **Competition** | -0.393 | 4.00e-4 | 0.0061 | **both** |
| **Memoryretrieval** | -0.388 | 0.0033 | 0.0169 | **both** |
| **Maintenance** | -0.387 | 0.0054 | 0.0229 | **both** |
| **Mentalimagery** | -0.380 | 0.0029 | 0.0166 | **both** |
| **Priming** | -0.366 | 0.0039 | 0.0192 | **both** |
| **Taskdifficulty** | -0.336 | 0.0054 | 0.0229 | **both** |
| **Strategy** | -0.333 | 0.0015 | 0.0115 | **both** |
| **Selectiveattention** | -0.322 | 0.0122 | 0.0469 | **both** |
| **Distraction** | -0.215 | 0.0083 | 0.0340 | **both** |
| Stress | 0.417 | 0.0163 | 0.0607 | p_spin only |
| Objectrecognition | -0.407 | 0.0288 | 0.0886 | p_spin only |
| Visualattention | -0.406 | 0.0192 | 0.0673 | p_spin only |
| Facerecognition | -0.368 | 0.0326 | 0.0955 | p_spin only |
| Reading | -0.366 | 0.0184 | 0.0666 | p_spin only |
| Motorcontrol | 0.363 | 0.0468 | 0.1251 | p_spin only |
| Morphology | 0.351 | 0.0279 | 0.0886 | p_spin only |
| Retention | -0.307 | 0.0225 | 0.0748 | p_spin only |
| Efficiency | -0.305 | 0.0197 | 0.0673 | p_spin only |
| Wordrecognition | -0.298 | 0.0346 | 0.0990 | p_spin only |
| Strength | 0.282 | 0.0413 | 0.1154 | p_spin only |
| Thought | -0.276 | 0.0283 | 0.0886 | p_spin only |
| Association | 0.262 | 0.0302 | 0.0906 | p_spin only |
| Concept | 0.193 | 0.0451 | 0.1233 | p_spin only |

### *Supplementary Table 21: Neurosynth functional decoding for the left LC–cortex G1 during negative movie-viewing, showing spin-test (p_spin_) and FDR-corrected association terms*

| **Term** | **ρ (rho)** | **p_spin** | **p_spin (FDR)** | **Correction** |
| --- | --- | --- | --- | --- |
| **Pain** | 0.568 | 4.00e-4 | 0.0109 | **both** |
| **Reading** | -0.529 | 6.00e-4 | 0.0109 | **both** |
| **Anticipation** | 0.495 | 8.00e-4 | 0.0109 | **both** |
| **Discrimination** | -0.478 | 4.00e-4 | 0.0109 | **both** |
| **Adaptation** | -0.469 | 4.00e-4 | 0.0109 | **both** |
| **Expertise** | -0.426 | 8.00e-4 | 0.0109 | **both** |
| **Induction** | 0.411 | 8.00e-4 | 0.0109 | **both** |
| **Wordrecognition** | -0.397 | 0.0041 | 0.0420 | **both** |
| **Hyperactivity** | 0.390 | 0.0047 | 0.0445 | **both** |
| **Strength** | 0.353 | 4.00e-4 | 0.0109 | **both** |
| **Naming** | -0.308 | 0.0011 | 0.0135 | **both** |
| **Concept** | 0.282 | 5.00e-4 | 0.0109 | **both** |
| **Competition** | -0.275 | 0.0039 | 0.0420 | **both** |
| Language | -0.458 | 0.0065 | 0.0566 | p_spin only |
| Perception | -0.451 | 0.0412 | 0.2460 | p_spin only |
| Inhibition | 0.436 | 0.0069 | 0.0566 | p_spin only |
| Decisionmaking | 0.434 | 0.0363 | 0.2460 | p_spin only |
| Sleep | 0.319 | 0.0278 | 0.2011 | p_spin only |
| Communication | -0.307 | 0.0474 | 0.2460 | p_spin only |
| Judgment | -0.269 | 0.0471 | 0.2460 | p_spin only |
| Balance | 0.266 | 0.0234 | 0.1799 | p_spin only |

### *Supplementary Table 22: Neurosynth functional decoding for the left LC–cortex G2 during negative movie-viewing, showing spin-test (p_spin_) and FDR-corrected association terms*

| **Term** | **ρ (rho)** | **p_spin** | **p_spin (FDR)** | **Correction** |
| --- | --- | --- | --- | --- |
| **Movement** | -0.669 | 4.00e-4 | 0.0019 | **both** |
| **Multisensory** | -0.666 | 4.00e-4 | 0.0019 | **both** |
| **Autobiographicalmemory** | 0.663 | 4.00e-4 | 0.0019 | **both** |
| **Imagery** | -0.638 | 4.00e-4 | 0.0019 | **both** |
| **Coordination** | -0.614 | 4.00e-4 | 0.0019 | **both** |
| **Risk** | 0.613 | 5.00e-4 | 0.0019 | **both** |
| **Recall** | 0.610 | 4.00e-4 | 0.0019 | **both** |
| **Action** | -0.603 | 4.00e-4 | 0.0019 | **both** |
| **Memoryretrieval** | 0.603 | 4.00e-4 | 0.0019 | **both** |
| **Mood** | 0.576 | 4.00e-4 | 0.0019 | **both** |
| **Psychosis** | 0.565 | 5.00e-4 | 0.0019 | **both** |
| **Semanticmemory** | 0.565 | 4.00e-4 | 0.0019 | **both** |
| **Perception** | -0.564 | 4.00e-4 | 0.0019 | **both** |
| **Socialcognition** | 0.559 | 4.00e-4 | 0.0019 | **both** |
| **Emotion** | 0.556 | 4.00e-4 | 0.0019 | **both** |
| **Emotionregulation** | 0.554 | 4.00e-4 | 0.0019 | **both** |
| **Episodicmemory** | 0.541 | 5.00e-4 | 0.0019 | **both** |
| **Manipulation** | -0.540 | 4.00e-4 | 0.0019 | **both** |
| **Gaze** | -0.515 | 4.00e-4 | 0.0019 | **both** |
| **Motorcontrol** | -0.513 | 9.00e-4 | 0.0026 | **both** |
| **Impulsivity** | 0.513 | 6.00e-4 | 0.0020 | **both** |
| **Retrieval** | 0.508 | 4.00e-4 | 0.0019 | **both** |
| **Context** | 0.500 | 4.00e-4 | 0.0019 | **both** |
| **Valence** | 0.495 | 8.00e-4 | 0.0025 | **both** |
| **Thought** | 0.492 | 5.00e-4 | 0.0019 | **both** |
| **Decision** | 0.485 | 6.00e-4 | 0.0020 | **both** |
| **Spatialattention** | -0.463 | 0.0024 | 0.0063 | **both** |
| **Discrimination** | -0.458 | 5.00e-4 | 0.0019 | **both** |
| **Rhythm** | -0.457 | 0.0046 | 0.0115 | **both** |
| **Fixation** | -0.456 | 6.00e-4 | 0.0020 | **both** |
| **Adaptation** | -0.450 | 4.00e-4 | 0.0019 | **both** |
| **Planning** | -0.449 | 0.0079 | 0.0174 | **both** |
| **Inference** | 0.449 | 4.00e-4 | 0.0019 | **both** |
| **Cognitivecontrol** | 0.445 | 5.00e-4 | 0.0019 | **both** |
| **Association** | 0.433 | 5.00e-4 | 0.0019 | **both** |
| **Objectrecognition** | -0.430 | 0.0168 | 0.0337 | **both** |
| **Belief** | 0.424 | 4.00e-4 | 0.0019 | **both** |
| **Reasoning** | 0.416 | 7.00e-4 | 0.0023 | **both** |
| **Salience** | 0.410 | 5.00e-4 | 0.0019 | **both** |
| **Selectiveattention** | -0.409 | 5.00e-4 | 0.0019 | **both** |
| **Judgment** | 0.408 | 9.00e-4 | 0.0026 | **both** |
| **Visualperception** | -0.404 | 0.0011 | 0.0031 | **both** |
| **Decisionmaking** | 0.400 | 0.0077 | 0.0172 | **both** |
| **Visualattention** | -0.393 | 0.0212 | 0.0414 | **both** |
| **Stress** | 0.392 | 0.0069 | 0.0159 | **both** |
| **Knowledge** | 0.389 | 8.00e-4 | 0.0025 | **both** |
| **Intention** | 0.388 | 5.00e-4 | 0.0019 | **both** |
| **Skill** | -0.387 | 0.0070 | 0.0159 | **both** |
| **Memory** | 0.382 | 0.0086 | 0.0186 | **both** |
| **Anxiety** | 0.381 | 0.0170 | 0.0337 | **both** |
| **Mentalimagery** | -0.379 | 0.0028 | 0.0072 | **both** |
| **Addiction** | 0.362 | 0.0122 | 0.0254 | **both** |
| **Localization** | -0.354 | 1.00e-3 | 0.0029 | **both** |
| **Strategy** | 0.351 | 4.00e-4 | 0.0019 | **both** |
| **Uncertainty** | 0.348 | 5.00e-4 | 0.0019 | **both** |
| **Languagecomprehension** | 0.345 | 0.0059 | 0.0142 | **both** |
| **Hyperactivity** | 0.333 | 0.0061 | 0.0144 | **both** |
| **Strength** | 0.305 | 6.00e-4 | 0.0020 | **both** |
| **Utility** | 0.303 | 0.0057 | 0.0140 | **both** |
| **Naming** | -0.302 | 0.0018 | 0.0049 | **both** |
| **Reinforcementlearning** | 0.300 | 0.0132 | 0.0271 | **both** |
| **Intelligence** | 0.260 | 0.0104 | 0.0221 | **both** |
| **Concept** | 0.249 | 0.0020 | 0.0053 | **both** |
| Responseselection | -0.350 | 0.0438 | 0.0792 | p_spin only |
| Detection | -0.328 | 0.0459 | 0.0818 | p_spin only |
| Loss | 0.319 | 0.0277 | 0.0532 | p_spin only |
| Sentencecomprehension | 0.259 | 0.0333 | 0.0621 | p_spin only |
| Consolidation | 0.259 | 0.0283 | 0.0535 | p_spin only |
| Learning | -0.229 | 0.0377 | 0.0692 | p_spin only |

### *Supplementary Table 23: Neurosynth functional decoding for the right LC–cortex G1 during negative movie-viewing, showing spin-test (p_spin_) and FDR-corrected association terms*

| **Term** | **ρ (rho)** | **p_spin** | **p_spin (FDR)** | **Correction** |
| --- | --- | --- | --- | --- |
| Language | 0.541 | 0.0065 | 0.0888 | p_spin only |
| Recognition | 0.510 | 0.0292 | 0.2762 | p_spin only |
| Pain | -0.492 | 0.0086 | 0.1058 | p_spin only |
| Reading | 0.483 | 0.0056 | 0.0861 | p_spin only |
| Meaning | 0.449 | 0.0029 | 0.0713 | p_spin only |
| Wordrecognition | 0.447 | 0.0041 | 0.0840 | p_spin only |
| Facialexpression | 0.436 | 0.0356 | 0.2919 | p_spin only |
| Knowledge | 0.436 | 1.00e-3 | 0.0713 | p_spin only |
| Priming | 0.419 | 0.0027 | 0.0713 | p_spin only |
| Judgment | 0.399 | 0.0028 | 0.0713 | p_spin only |
| Communication | 0.394 | 0.0049 | 0.0861 | p_spin only |
| Induction | -0.356 | 0.0022 | 0.0713 | p_spin only |
| Competition | 0.241 | 0.0248 | 0.2542 | p_spin only |
| Strength | -0.234 | 0.0098 | 0.1096 | p_spin only |
| Concept | -0.146 | 0.0345 | 0.2919 | p_spin only |

### *Supplementary Table 24: Neurosynth functional decoding for the right LC–cortex G2 during negative movie-viewing, showing spin-test (p_spin_) and FDR-corrected association terms*

| **Term** | **ρ (rho)** | **p_spin** | **p_spin (FDR)** | **Correction** |
| --- | --- | --- | --- | --- |
| **Multisensory** | -0.699 | 4.00e-4 | 0.0020 | **both** |
| **Perception** | -0.619 | 4.00e-4 | 0.0020 | **both** |
| **Gaze** | -0.589 | 4.00e-4 | 0.0020 | **both** |
| **Spatialattention** | -0.584 | 4.00e-4 | 0.0020 | **both** |
| **Discrimination** | -0.580 | 4.00e-4 | 0.0020 | **both** |
| **Adaptation** | -0.572 | 4.00e-4 | 0.0020 | **both** |
| **Risk** | 0.568 | 4.00e-4 | 0.0020 | **both** |
| **Objectrecognition** | -0.551 | 4.00e-4 | 0.0020 | **both** |
| **Mood** | 0.539 | 4.00e-4 | 0.0020 | **both** |
| **Manipulation** | -0.510 | 4.00e-4 | 0.0020 | **both** |
| **Stress** | 0.506 | 4.00e-4 | 0.0020 | **both** |
| **Action** | -0.506 | 0.0031 | 0.0089 | **both** |
| **Autobiographicalmemory** | 0.504 | 4.00e-4 | 0.0020 | **both** |
| **Selectiveattention** | -0.500 | 4.00e-4 | 0.0020 | **both** |
| **Visualattention** | -0.497 | 5.00e-4 | 0.0022 | **both** |
| **Psychosis** | 0.490 | 7.00e-4 | 0.0027 | **both** |
| **Fixation** | -0.486 | 4.00e-4 | 0.0020 | **both** |
| **Mentalimagery** | -0.485 | 4.00e-4 | 0.0020 | **both** |
| **Memoryretrieval** | 0.485 | 4.00e-4 | 0.0020 | **both** |
| **Impulsivity** | 0.480 | 5.00e-4 | 0.0022 | **both** |
| **Imagery** | -0.479 | 0.0014 | 0.0045 | **both** |
| **Emotionregulation** | 0.479 | 4.00e-4 | 0.0020 | **both** |
| **Attention** | -0.473 | 7.00e-4 | 0.0027 | **both** |
| **Recall** | 0.463 | 7.00e-4 | 0.0027 | **both** |
| **Association** | 0.456 | 4.00e-4 | 0.0020 | **both** |
| **Movement** | -0.451 | 0.0183 | 0.0409 | **both** |
| **Decision** | 0.448 | 4.00e-4 | 0.0020 | **both** |
| **Visualperception** | -0.447 | 0.0013 | 0.0043 | **both** |
| **Detection** | -0.445 | 6.00e-4 | 0.0025 | **both** |
| **Belief** | 0.441 | 4.00e-4 | 0.0020 | **both** |
| **Socialcognition** | 0.430 | 4.00e-4 | 0.0020 | **both** |
| **Inference** | 0.429 | 5.00e-4 | 0.0022 | **both** |
| **Decisionmaking** | 0.426 | 0.0015 | 0.0046 | **both** |
| **Semanticmemory** | 0.424 | 0.0161 | 0.0374 | **both** |
| **Thought** | 0.420 | 4.00e-4 | 0.0020 | **both** |
| **Coordination** | -0.418 | 0.0219 | 0.0473 | **both** |
| **Loss** | 0.417 | 0.0011 | 0.0038 | **both** |
| **Addiction** | 0.415 | 0.0016 | 0.0048 | **both** |
| **Context** | 0.406 | 0.0029 | 0.0085 | **both** |
| **Hyperactivity** | 0.395 | 9.00e-4 | 0.0033 | **both** |
| **Cognitivecontrol** | 0.391 | 1.00e-3 | 0.0035 | **both** |
| **Episodicmemory** | 0.386 | 0.0152 | 0.0360 | **both** |
| **Strength** | 0.363 | 4.00e-4 | 0.0020 | **both** |
| **Concept** | 0.356 | 4.00e-4 | 0.0020 | **both** |
| **Salience** | 0.349 | 0.0032 | 0.0089 | **both** |
| **Utility** | 0.344 | 8.00e-4 | 0.0030 | **both** |
| **Retrieval** | 0.342 | 0.0106 | 0.0256 | **both** |
| **Uncertainty** | 0.329 | 4.00e-4 | 0.0020 | **both** |
| **Reasoning** | 0.327 | 0.0048 | 0.0123 | **both** |
| **Expertise** | -0.322 | 0.0045 | 0.0120 | **both** |
| **Reinforcementlearning** | 0.304 | 0.0206 | 0.0452 | **both** |
| **Naming** | -0.301 | 0.0015 | 0.0046 | **both** |
| **Consolidation** | 0.299 | 0.0054 | 0.0136 | **both** |
| **Localization** | -0.282 | 0.0174 | 0.0396 | **both** |
| **Intention** | 0.273 | 0.0048 | 0.0123 | **both** |
| **Strategy** | 0.252 | 0.0044 | 0.0120 | **both** |
| **Intelligence** | 0.244 | 0.0102 | 0.0251 | **both** |
| Emotion | 0.427 | 0.0261 | 0.0553 | p_spin only |
| Valence | 0.422 | 0.0267 | 0.0557 | p_spin only |
| Integration | -0.375 | 0.0459 | 0.0911 | p_spin only |
| Sleep | 0.274 | 0.0276 | 0.0566 | p_spin only |
| Interference | -0.264 | 0.0417 | 0.0841 | p_spin only |
